# Vasohibin1, a new IRES trans-acting factor for induction of (lymph)angiogenic factors in early hypoxia

**DOI:** 10.1101/260364

**Authors:** Fransky Hantelys, Anne-Claire Godet, Florian David, Florence Tatin, Edith Renaud-Gabardos, Françoise Pujol, Leila Diallo, Isabelle Ader, Laetitia Ligat, Anthony K. Henras, Yasufumi Sato, Angelo Parini, Eric Lacazette, Barbara Garmy-Susini, Anne-Catherine Prats

**Affiliations:** UMR 1048-I2MC, Inserm, Université de Toulouse, UPS, Toulouse, France; UMR 1031-STROMALAB, Inserm, CNRS ERL5311, Etablissement Français du Sang-Occitanie (EFS), National Veterinary School of Toulouse (ENVT), Université de Toulouse, UPS, Toulouse, France; UMR 1037-CRCT, Inserm, CNRS, Université de Toulouse, UPS, Pôle Technologique-Plateau Protéomique, Toulouse, France; UMR 5099-LBME, CBI, CNRS, Université de Toulouse, UPS, Toulouse, France; Department of Vascular Biology, Institute of Development, Aging and Cancer, Tohoku University, Sendai, Japan

**Keywords:** cardiomyocyte, hypoxia, IRES, translational control, vasohibin

## Abstract

Hypoxia, a major inducer of angiogenesis, is known to trigger major changes of gene expression at the transcriptional level. Furthermore, global protein synthesis is blocked while internal ribosome entry sites (IRES) allow specific mRNAs to be translated. Here we report the transcriptome and translatome signatures of (lymph)angiogenic genes in hypoxic HL-1 cardiomyocytes: most genes are not induced at the transcriptome-, but at the translatome level, including all IRES-containing mRNAs. Our data reveal activation of (lymph)angiogenic mRNA IRESs in early hypoxia. We identify vasohibin1 (VASH1) as an IRES trans-acting factor (ITAF) able to activate FGF1 and VEGFD IRESs in hypoxia while it inhibits several IRESs in normoxia. Thus this new ITAF may have opposite effects on IRES activities. These data suggest a generalized process of IRES-dependent translational induction of (lymph)angiogenic growth factors expression in early hypoxia, whose pathophysiological relevance is to trigger formation of new functional vessels in ischemic heart. VASH1 is not always required, indicating that the IRESome composition is variable, thus allowing subgroups of IRESs to be activated under the control of different ITAFs.

## INTRODUCTION

Hypoxia constitutes a major stress in different pathologies including cancer, as well as ischemic pathologies where artery occlusion leads to hypoxic conditions. In all these pathologies, hypoxia induces a cell response that stimulates angiogenesis to re-feed starved cells with oxygen and nutrients (1). Recently it has been shown that lymphangiogenesis is also induced by hypoxia (2). Hypoxia-induced (lymph)angiogenesis is mediated by strong modification of gene expression at both transcriptional and post-transcriptional levels (1, 3). A major way of gene expression regulation is mediated at the transcriptional level by the hypoxia inducible factor 1 (HIF1), a transcription factor stabilized by oxygen deprivation, that activates transcription from promoters containing hypoxia responsive elements (HRE). One of the well-described HIF1 targets is vascular endothelial growth factor A (VEGFA), a major angiogenic factor (4, 5). However, two other major angiogenic or lymphangiogenic growth factors, fibroblast growth factor 2 (FGF2) and VEGFC, respectively, are induced by hypoxia in a HIF-independent manner by a translational mechanism, indicating the importance of the post-transcriptional regulation of gene expression in this process (2, 6).

Translational control of gene expression plays a crucial role in the stress response. In particular, translation of most mRNAs, occurring by the classical cap-dependent mechanism, is silenced whereas alternative translation mechanisms allow enhanced expression of a small group of messengers involved in the control of cell survival (3, 7, 8). One of the major alternative mechanisms able to overcome this global inhibition of translation by stress depends on internal ribosome entry sites (IRESs) that correspond to RNA structural elements allowing the direct recruitment of the ribosome on mRNA. As regards the molecular mechanisms of IRES activation by stress, several studies have reported the involvement of RNA binding proteins, called IRES trans-acting factors (ITAFs), able to stabilize the adequate RNA conformation allowing ribosome recruitment (9–13). Interestingly, subcellular relocalization of ITAFs plays a critical role in IRES-dependent translation (14). Indeed, many RNA-binding proteins are known to shuttle between nucleus and cytoplasm, and it has been reported that cytoplasmic relocalization of ITAFs such as PTB, PCBP1, RBM4 or nucleolin is critical to activate IRES-dependent translation (10, 13, 14). In contrast, other ITAFs such as hnRNPA1, may have a negative impact on IRES activity when accumulating in the cytoplasm (15). However, how ITAFs participate in the regulation of the hypoxic response remains a challenging question to address.

IRESs are present in the mRNAs of several (lymph)angiogenic growth factors in the FGF and VEGF families, suggesting that the IRES-dependent mechanism might be a major way to activate angiogenesis and lymphangiogenesis during stress (2, 10, 13, 16–19). However, most studies of the role of hypoxia in gene expression regulation have been performed in tumoral hypoxia, while it has been reported that tumoral angiogenesis leads to formation of abnormal vessels that are non functional, which strongly differs from non tumoral angiogenesis that induces formation of functional vessels (20). This suggests that gene expression regulation in response to hypoxia may be different in cancer versus ischemic pathologies. In particular, the role of IRESs in the control of gene expression in ischemic heart, the most frequent ischemic pathology, remains to be elucidated.

Here we analyzed the transcriptome and the translatome of (lymph)angiogenic growth factors in hypoxic cardiomyocytes, and studied regulation of IRES activities in early and late hypoxia. Data show that in cardiomyocyte, (lymph)angiogenic growth factors are mostly regulated at the translational level. Interestingly, FGF and VEGF mRNA IRESs are sequentially activated at different times of early hypoxia in contrast to IRESs of non angiogenic messengers. We also looked for ITAFs governing IRES activation in hypoxia and identified vasohibin1 (VASH1) as a new ITAF specific of the earliest activated FGF1 IRES in cardiomyocytes. VASH1 knock-down strongly down-regulates the earliest-induced FGF1 IRES but not the other IRESs, revealing that this protein is a new IRES trans-acting factor (ITAF) in cardiomyocytes, specific of early hypoxia.

## RESULTS

### Most (lymph)angiogenic genes are not induced at the transcriptome level of hypoxic cardiomyocytes

In order to analyze expression of angiogenic and lymphangiogenic growth factors in hypoxic cardiomyocytes, the HL-1 cell line was chosen: although immortalized, it keeps the beating phenotype specific to cardiomyocyte (21). HL-1 cells were submitted to increasing times of hypoxia, from 5 minutes to 24 hours and their trancriptome was analyzed on a Fluidigm Deltagene PCR array targeting 96 genes of angiogenesis, lymphangiogenesis and/or stress (Fig. 1, EV Fig. 1, EV Table 1). Data showed a significant increase of *Vegfa*, PAI-1 and apelin (*Apln*) mRNA levels, with a peak at 8h of hypoxia for *Vegfa* and PAI1 and 24h for *Apln*. These three genes are well described HIF1 targets (4, 22, 23). However, only 5-8% of the genes were induced, while the mRNA levels of several major angio- or lymphangiogenic factors, such as FGF2 and VEGFC, were strongly decreased after 4 h or 8 h of hypoxia. These data indicate that the transcriptional response to hypoxia in cardiomyocytes is not the major mechanism controlling expression of (lymph)angiogenic factors, suggesting that post-transcriptional mechanisms are involved.

**Figure 1.**
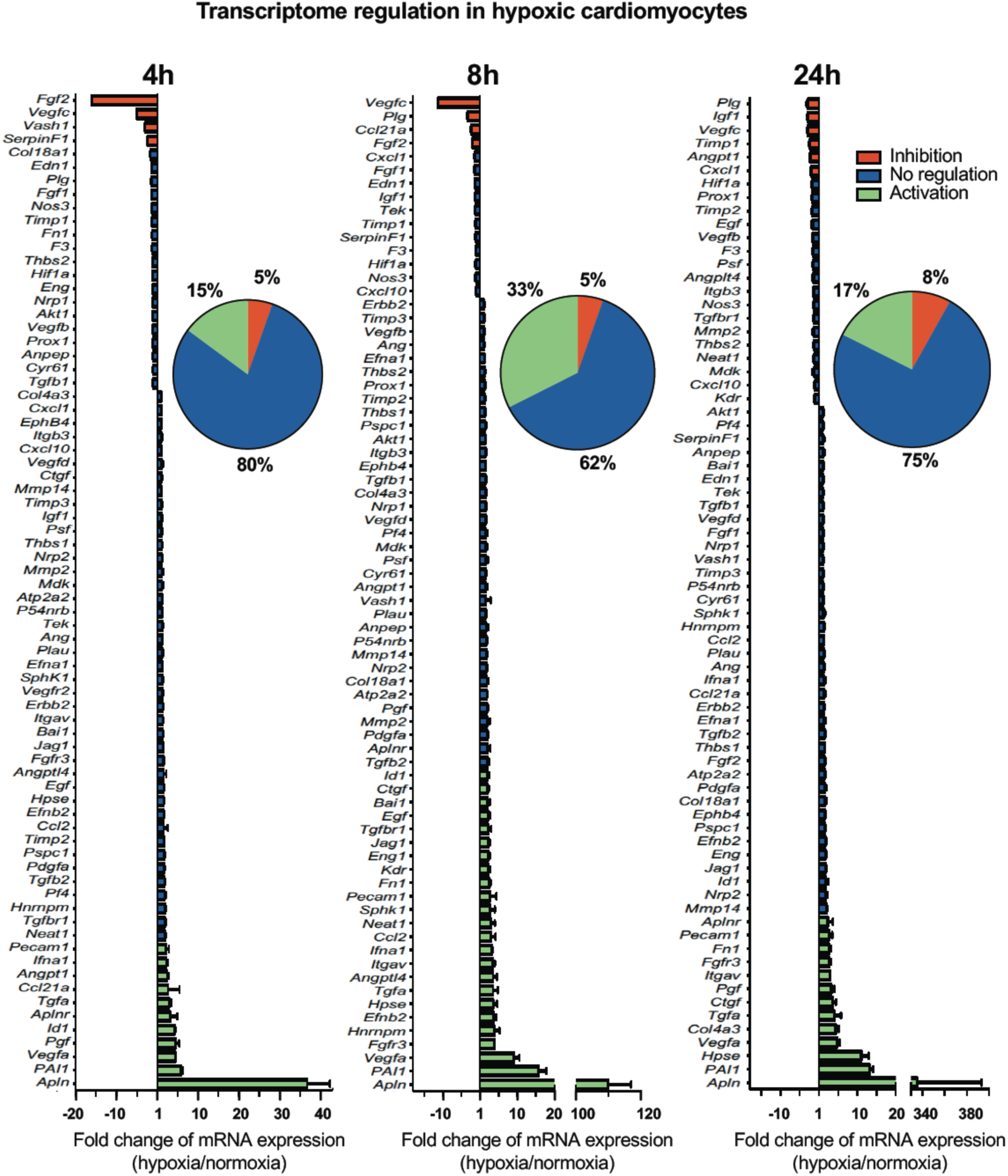
Most (lymph)angiogenic genes are not induced at the transcriptome level in hypoxic cardiomyocytes. Total RNA was purified from HL-1 cardiomyocytes submitted to increasing times from 5 min to 24 h of hypoxia at 1% O_2_, as well as from normoxic cardiomyocytes as a control. cDNAs were synthesized and used for a Fluidigm deltagene PCR array dedicated to genes related to (lymph)angiogenesis or stress (EV Table 6). Relative quantification (RQ) of gene expression during hypoxia was calculated using the 2^-ΔΔCT^ method with normalization to 18S and to normoxia. mRNA levels are presented by histograms for the times of 4 h, 8 h and 24 h, as the fold change of repression (red) or induction (green) normalized to normoxia. Non-regulated mRNAs are represented in blue. When the RQ value is inferior to 1, the fold change is expressed as −1/RQ. The percentage of repressed, induced, and non-regulated mRNAs is indicated for each time. For earlier times of 5 min to 2 hr, the percentages are shown in EV Fig. 1. The detailed values for all the times of the kinetics are presented in EV Table 1.

### mRNAs of most (lymph)angiogenic genes are recruited into polysomes in hypoxic cardiomyocytes

Based on the fact that mRNA present in polysomes are actively translated, we tested the hypothesis of translational induction by analysing the recruitment of mRNAs into polysomes. This experiment was performed in early and late hypoxia. The polysome profile showed that translational activity in normoxic HL-1 cells was low but decreased after 4 h of hypoxia, with a shift of the polysome to monosome ratio from 1,55 to 1,40 (Fig. 2A). 4E-BP1 appeared as a single band and its phosphorylation profile did not change upon hypoxia, suggesting that it is already hypophosphorylated in normoxia in these cells (EV Fig. 2A and 2B). In contrast, translation blockade was confirmed by the strong phosphorylation of eIF2*α* (Fig. 2B, EV Fig.2C). 94% of the genes of the (lymph)angiogenic array showed a more sustained recruitment into polysomes under hypoxic conditions (Fig. 2C, EV Table 2). This translational induction not only targets major angiogenic factors and their receptors (*Vegfa, Fgf1, Pdgfa, Fgfr3, Vegfr2*…), but also genes involved in cardiomyocyte survival in ischemic heart (*Igf1, Igf1R*) or in inflammation (BAI1, *Tgfb*). These data suggest that in cardiomyocytes, the main response to early hypoxia of (lymph)angiogenic genes is not transcriptional, but translational.

**Figure 2.**
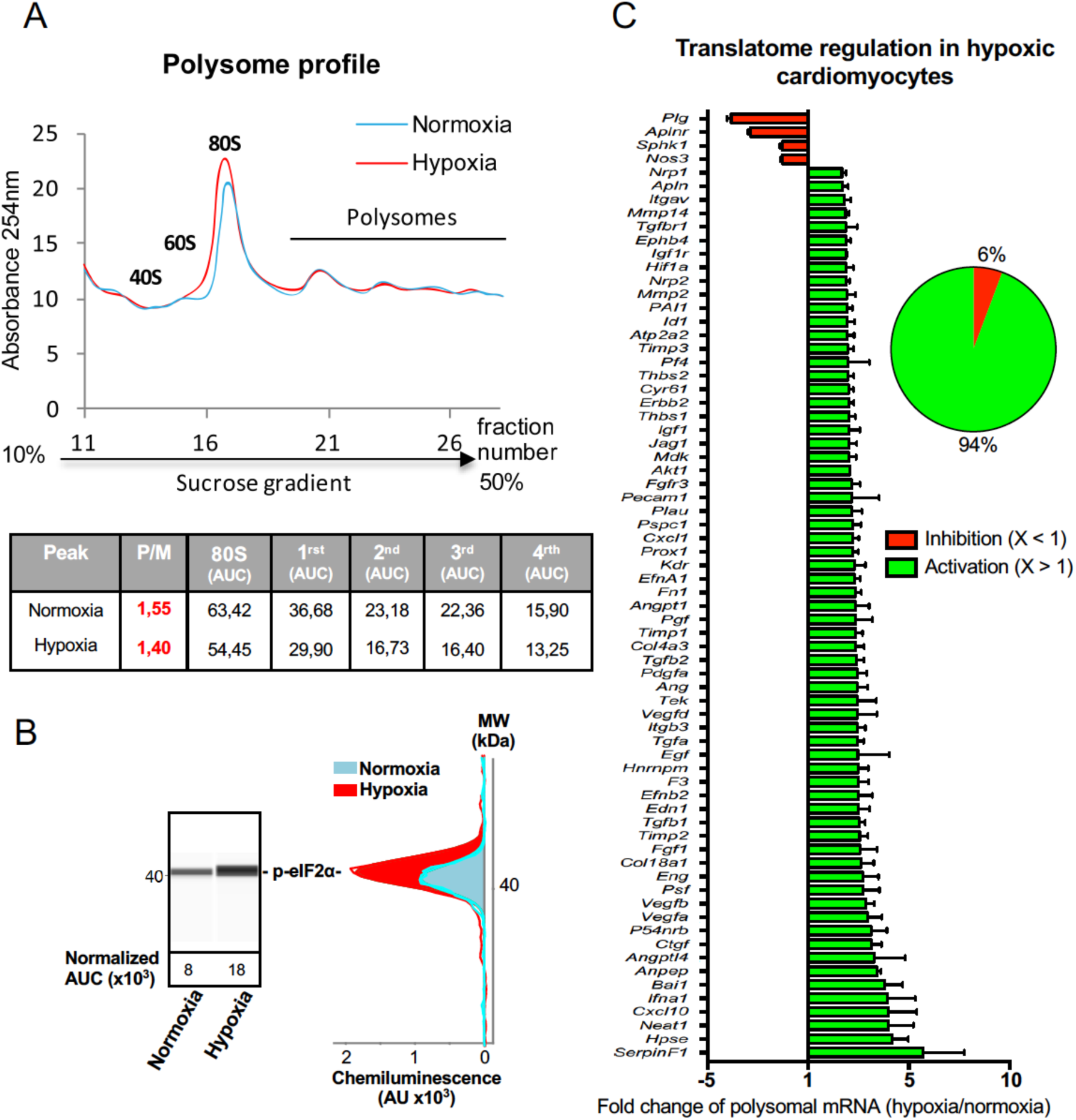
mRNAs of most (lymph)angiogenic genes are mobilized into polysomes in hypoxic cardiomyocytes. A-C In order to isolate translated mRNAs, polysomes were purified on sucrose gradient from HL-1 cardiomyocytes in normoxia or after 4 hr of hypoxia at 1% O_2_, as described in Materials and Methods. P/M ratio (polysome/monosome) was determined by delimiting the 80s and polysome peaks by taking the lowest plateau values between each peak and by calculating the area under the curve (AUC). Then the sum of area values of the four polysome peaks was divided by the area of the 80s peak (A). Translation blockade was measured by eIF2*α* phosphorylation quantification by capillary Simple Western using normalization against total protein as described in Mat. & Meth. (B). RNA was purified from polysome fractions and from cell lysate before loading. cDNA and PCR array were performed as in Figure 1. Polysomes profiles are presented for normoxic and hypoxic cardiomyocytes. Relative quantification (RQ) of gene expression during hypoxia was calculated using the 2^-ΔΔCT^ method with normalization to 18S and to normoxia. mRNA levels (polysomal RNA/total RNA) are shown as fold change of repression (red) or induction (green) in hypoxia normalized to normoxia as in Figure 1 (C). When the RQ value is inferior to 1, the fold change is expressed as −1/RQ. The detailed values are available in EV Table 2.

### IRES-containing mRNAs are more efficiently mobilized into polysomes under hypoxic conditions

IRES-dependent translation has been reported to drive translation of several mRNAs in stress conditions (2, 3, 6, 24). Thus we focused onto the regulation of the different IRES-containing mRNAs present in the Fluidigm array (Fig. 3). Interestingly, the only IRES-containing mRNA to be significantly induced at the transcriptome level by hypoxia was *Vegfa* (Fig. 3A and EV Fig. 3). Expression of the apelin receptor (*Aplnr*), presumably devoid of IRES but transcriptionally induced during hypoxia, is also shown for comparison.

**Figure 3.**
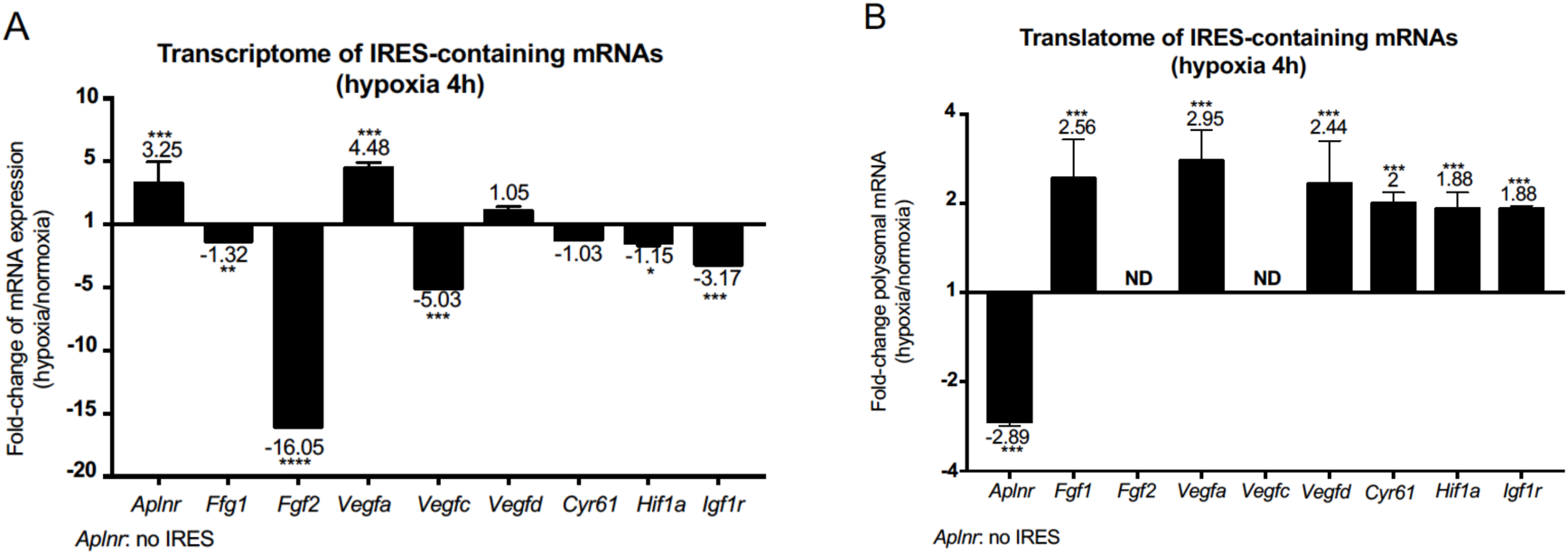
IRES-containing mRNAs are more efficiently associated to polysomes in hypoxic conditions. A-B RQ values for IRES-containing mRNA transcriptome (A) and translatome (B) extracted from the PCR arrays shown in Figures 1 and 2. The gene *Aplnr* (apelin receptor) was chosen as a control without an IRES. *Vegfc* and *Fgf2* mRNAs, repressed in the transcriptome, were below the detection threshold in polysomes (ND). Histograms correspond to means ± standard deviation, *p<0.05, **p<0.01, ***<0.001, ****p<0.0001 compared to normoxia.

Polysome recruitment of these IRES-containing mRNAs is shown in figure 3B. Clearly, *Fgf1, Vegfa, Vegfd, Cyr61*, *Hif1a* and *Igf1r* mRNAs were recruited into polysomes under hypoxia 2 to 3 times more than in normoxia, suggesting an important induction in terms of translation. In contrast, *Aplnr* mRNA recruited into polysomes decreased about three times. The data are not available for *Fgf2* and *Vegfc* mRNAs, which were not detectable. These results indicate that hypoxia in cardiomyocytes, although blocking global cap-dependent translation, induces translation of all detectable IRES-containing angiogenic factor mRNAs. This mechanism occurs as soon as 4 hours after oxygen deprivation, thus corresponding to an early event in the hypoxic response.

### IRESs of (lymph)angiogenic factor mRNAs are activated during early hypoxia

To confirm that the polysome recruitment of IRES-containing mRNAs actually corresponds to a stimulation of IRES-dependent translation, IRESs from FGF and VEGF mRNAs were introduced into a bicistronic dual luciferase gene expression cassette (Fig. 4A). As controls, two IRESs from non angiogenic mRNAs, c-myc and EMCV IRESs, were used. A negative control without IRES was provided by an hairpin inserted between the two cistrons (25). The well established bicistronic vector strategy, previously validated by us and others, allows to measure IRES activity revealed by expression of the second cistron, LucF (2, 25). The bicistronic cassettes were subcloned into lentivectors, as HL-1 cells are not efficiently transfected by plasmids but can be easily transduced by lentivectors with an efficiency of more than 80% (not shown). HL-1 cardiomyocytes were first transduced with the lentivector containing the FGF1 IRES and a kinetics was performed from 1 to 24 hours of hypoxia. Luciferase activities were measured from cell extracts and IRES activities reported as the LucF/LucR luminescence ratio. Data showed an increase of IRES activity between 4 to 8 hours whereas it decreased from 16 h to 24 h (Fig. 4B). Expression of endogenous FGF1 was analyzed after 8 hours of hypoxia. FGF1 protein quantification normalized to total proteins showed that IRES induction correlates with an increased expression of FGF1 protein (Fig. 4C). This is also consistent with the increase of FGF1 mRNA recruitment into polysomes observed above (Fig. 3C, EV Table 2). To determine whether this transient induction could affect other IRESs, HL-1 cells were then transduced by the complete series of lentivectors described above (Fig. 4A) and submitted to 4, 8 or 24 hours of hypoxia. Results showed an increase of all FGF and VEGF IRES activities in early hypoxia (4 hours and/or 8 hours), while the c-myc IRES was activated only in late hypoxia after 24 hours (Fig. 4D). The viral EMCV IRES was activated in both early (4 hours) and late (24 hours) hypoxia. The hairpin control was not induced (EV Table 3J). These data revealed two waves of IRES activation in response to hypoxia: a first wave concerns IRESs from (lymph)angiogenic growth factor mRNAs that are activated during early hypoxia, while a second wave concerns “non-angiogenic” c-myc and EMCV IRESs, that are activated in late hypoxia.

**Figure 4.**
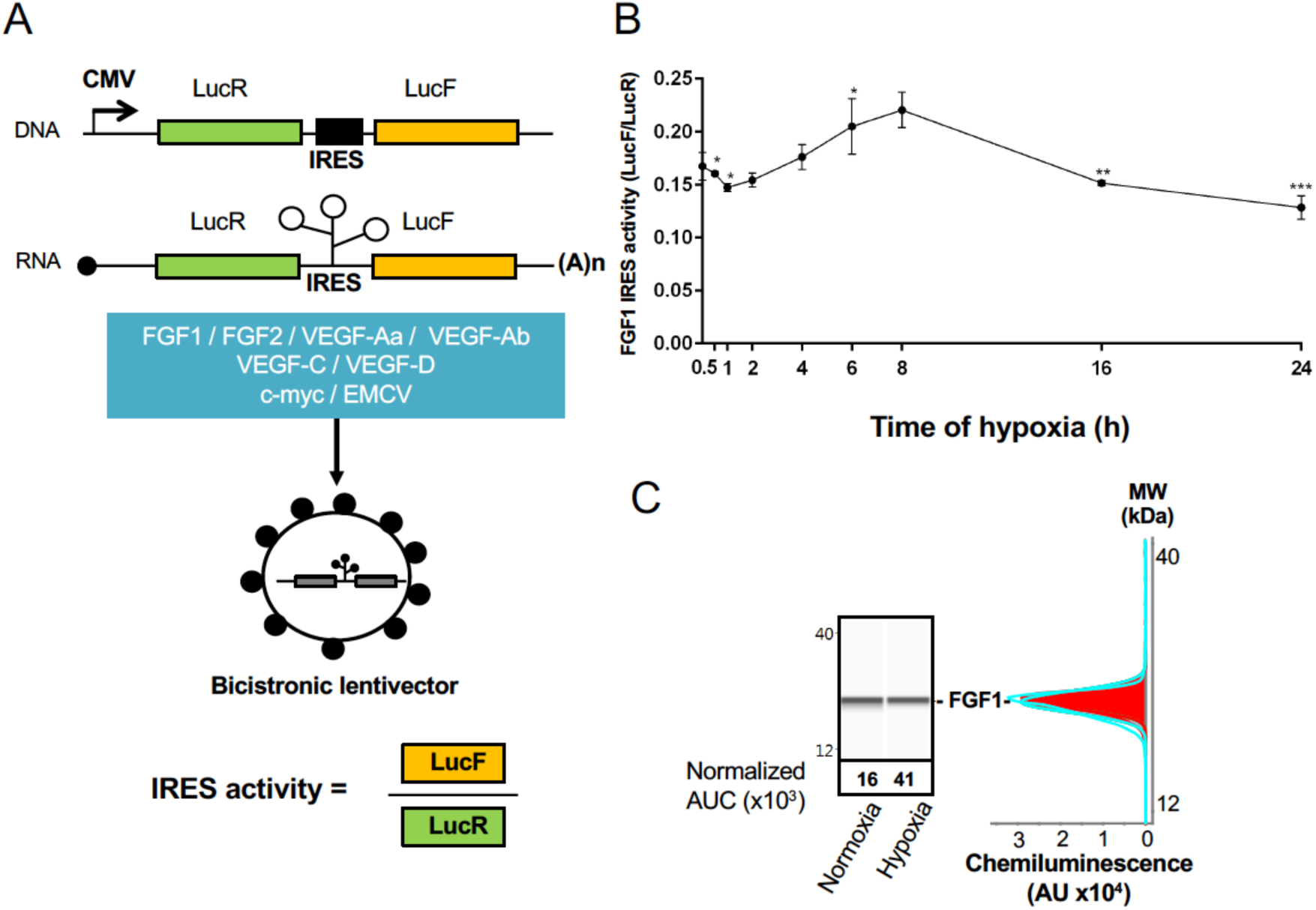
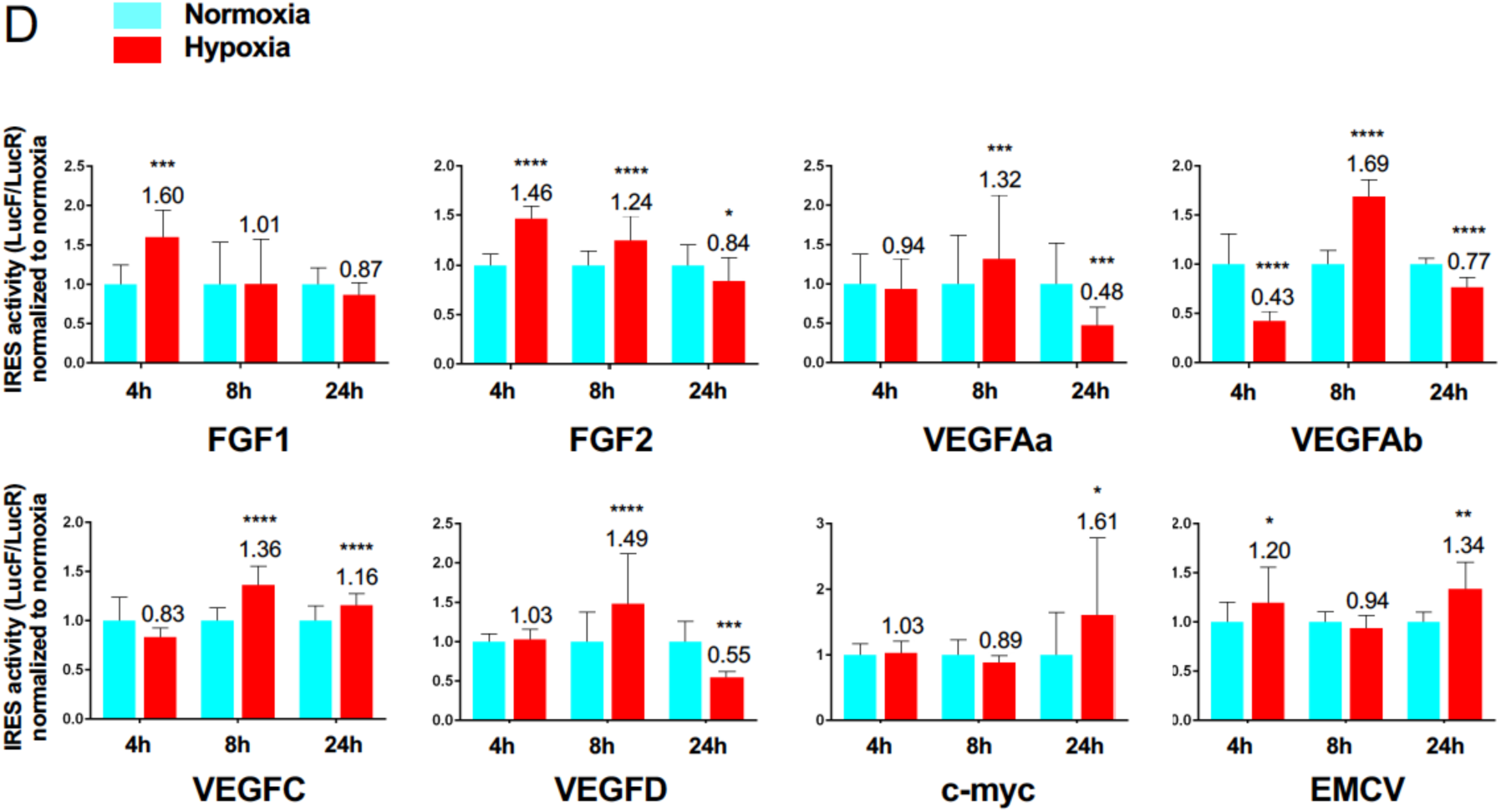
IRESs from (lymph)angiogenic factor mRNAs are activated in early hypoxia. A-D To measure IRES-dependent translation during hypoxia, HL-1 cardiomyocytes were transduced with bicistronic dual luciferase lentivectors (termed “Lucky-Luke”) containing different IRESs cloned between the genes of renilla (LucR) and firefly (LucF) luciferase (A). In bicistronic vectors, translation of the first cistron LucR is cap-dependent whereas translation of the second cistron LucF is IRES-dependent (25). Cardiomyocytes transduced by the CRF1AL+ lentivector Lucky-Luke reporter containing FGF1 IRES were submitted to an hypoxia time-course (0h, 1h, 2h, 4h, 6h, 8h, 16h and 24h) and each time point compared to 0h point with one-tailed t-test (B). Endogenous FGF1 protein expression was measured by capillary Simple Western, from extracts of cardiomyocytes in normoxia or submitted to 8h of hypoxia (C). HL-1 cardiomyocytes transduced by different Lucky-Luke constructs were submitted to 4h, 8h or 24h of hypoxia and luciferase activities measured. IRES activities during hypoxia, expressed as LucF/LucR ratio, are normalized to normoxia. Three groups of activation were identified: FGF IRESs, VEGF IRESs, non-angiogenic IRESs at 4h, 8h or 24h of hypoxia, respectively. Histograms correspond to means ± standard deviation of the mean, with a one-tailed t-test *p<0.05, **p<0.01, ***<0.001, ****p<0.0001, compared to normoxia. For each IRES the mean has been calculated from three independent experiments with three biological replicates (n=9). All detailed values are presented in EV Table 3. A no-IRES control was also performed and values are presented in EV Table 3 J.

### Identification of IRES-bound proteins in hypoxic cardiomyocytes reveals vasohibin1 as a new RNA-binding protein

Early activation of angiogenic factor IRESs during hypoxia suggested that specific ITAFs may be involved between 4 and 8 hours. In an attempt to identify such ITAFs, we used the technology of biomolecular analysis coupled to mass spectrometry (BIA-MS), validated for ITAF identification in two previous studies (13, 26). Biotinylated RNAs corresponding to FGF1 (4 hours activation), VEGFAa (8 hours activation) and EMCV IRESs (24 hours activation) were used as probes for BIA-MS. Hooked proteins from normoxic and hypoxic HL-1 cells were then recovered and identified (Fig. 5A-B, EV Table 4). Surprisingly, except for nucleolin bound to VEGFAa and EMCV IRES in normoxia, no known ITAF was identified as bound to these IRESs in normoxia or in hypoxia. Interestingly, besides several proteins unrelated to (lymph)angiogenesis, we detected the presence of vasohibin1 (VASH1), a protein described as an endothelial cell-produced angiogenesis inhibitor, but also for its role in stress tolerance and cell survival (Fig. 5C) (27, 28). However, this secreted protein has never been reported for any RNA-binding activity. VASH1 bound to the FGF1 IRES under 4 hours or 8 hours of hypoxia, but not under normoxia (EV Table 4). This protein also bound to the EMCV IRES both in normoxia and hypoxia but not to the VEGFA IRES. In order to address the RNA binding potential of VASH1, we performed an *in silico* analysis of VASH1 protein sequence that predicted two conserved RNA-binding domains (RBD) in the N- and C-terminal parts of the full length protein, respectively (Fig. 5C, EV Fig. 4A and 4B). The direct interaction of VASH1 with FGF1, VEGFAa and EMCV IRESs was assessed by surface plasmon resonance using the full length recombinant 44 kDa protein, resulting in the measurement of affinity constants of 6.5 nM, 8.0 nM and 9.6 nM, respectively (Fig. 5D-5F). These data indicate that VASH1 exhibits a significant RNA binding activity.

**Figure 5.**
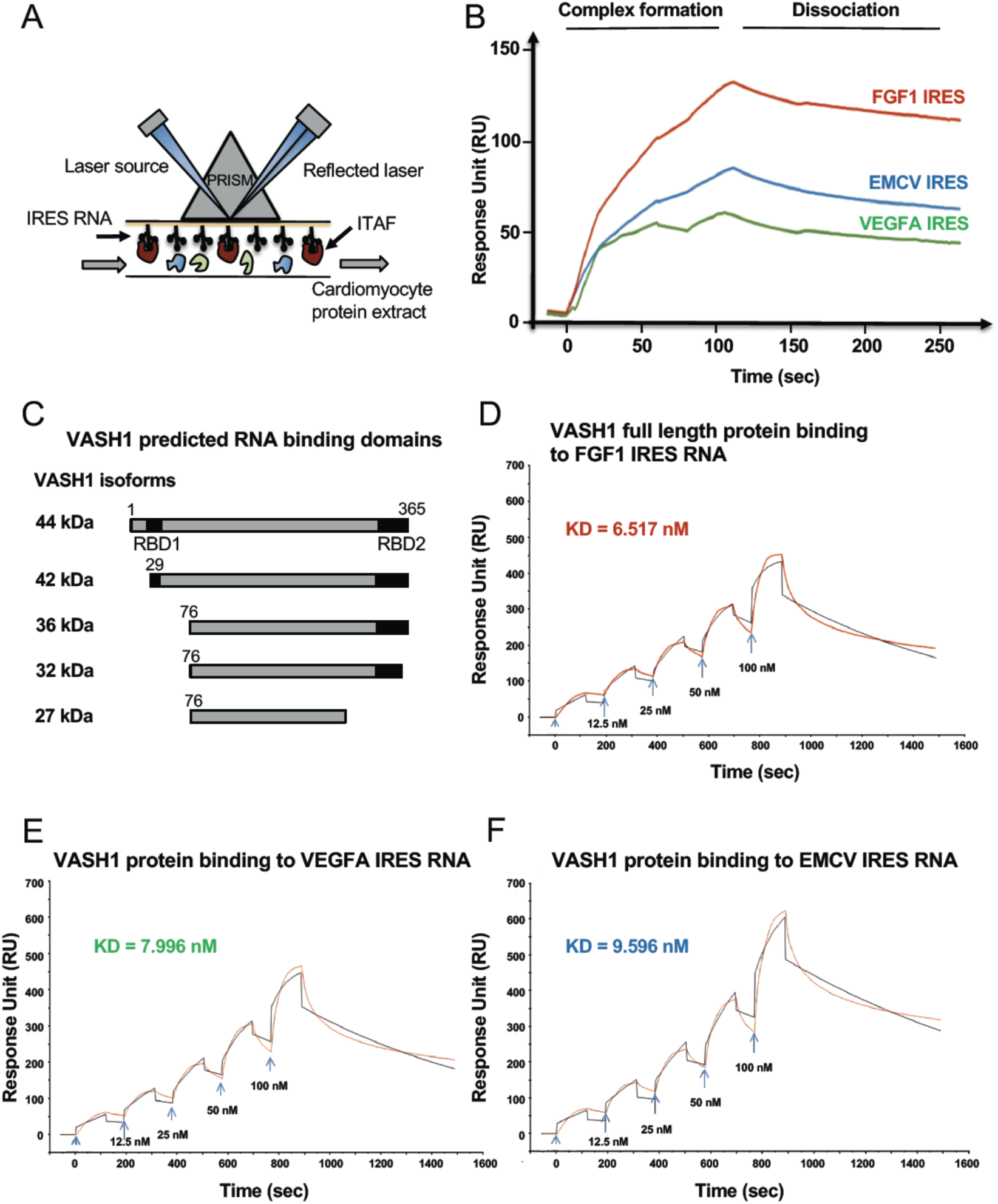
Identification of IRES-bound proteins in hypoxic cardiomyocytes reveals vasohibin1 as a new RNA-binding protein. A-F Biotinylated IRES RNAs were transcribed *in vitro* and immobilized on the sensorchip of the BIAcore T200 optical biosensor device (A). Total cell extracts from normoxic or hypoxic HL-1 cardiomyocytes were injected in the device. Complex formation and dissociation were measured (see Mat. & Meth) (B). Bound proteins were recovered as described in Mat. & Meth. and identified by mass spectrometry (LC-MS/MS) after tryptic digestion. The list of proteins bound in normoxia and hypoxia to FGF1, VEGFAa and EMCV IRESs is shown in EV Table 4. VASH1 protein was identified bound to FGF1 (hypoxia) and EMCV IRESs (hypoxia and normoxia), but not to VEGFA IRES. A diagram of VASH1 RNA-binding properties is shown, with VASH1 isoforms described by Sonoda et al (**37**) (C). The predicted RNA binding domains (RBD1 and RBD2) shown in EV Figure 4, conserved in mouse and human, are indicated (C). Recombinant full-length 44 kDa VASH1 was injected into the Biacore T200 device containing immobilized FGF1 (D), VEGFAa (E) or EMCV (F) IRES as above. The affinity constants (KD) were calculated (D, E, F) with a Single Cycle Kinetics (SCK) strategy.

### Vasohibin1 is translationally induced and nuclearized in early hypoxia

VASH1 has been previously described for its expression in endothelial cells but never in cardiomyocytes (27). The present BIA-MS study provides evidence that it is expressed in HL-1 cardiomyocytes (EV Table 4). We analyzed the regulation of VASH1 expression during hypoxia: *Vash1* mRNA level strongly decreases after 4 hours of hypoxia whereas it is slightly upregulated after 8 hours (EV Table 1, Fig. 6A). In contrast, analysis of *Vash1* mRNA recruitment into polysomes showed a strong increase at 4 hours of hypoxia (about 7 times)(Fig. 6B), whereas it was not detectable in polysomes at 24 hours (EV Table 2). This indicates that *Vash1* mRNA translation is strongly induced in early hypoxia. VASH1 immunodetection confirmed a strong expression of VASH1 at 4 hours of hypoxia, despite the decrease of its mRNA. VASH1 appeared as foci in both cytoplasm and nucleus (Fig. 6C). The number of foci did not change, but their size significantly increased in hypoxia (Fig. 6D and 6E).

**Figure 6.**
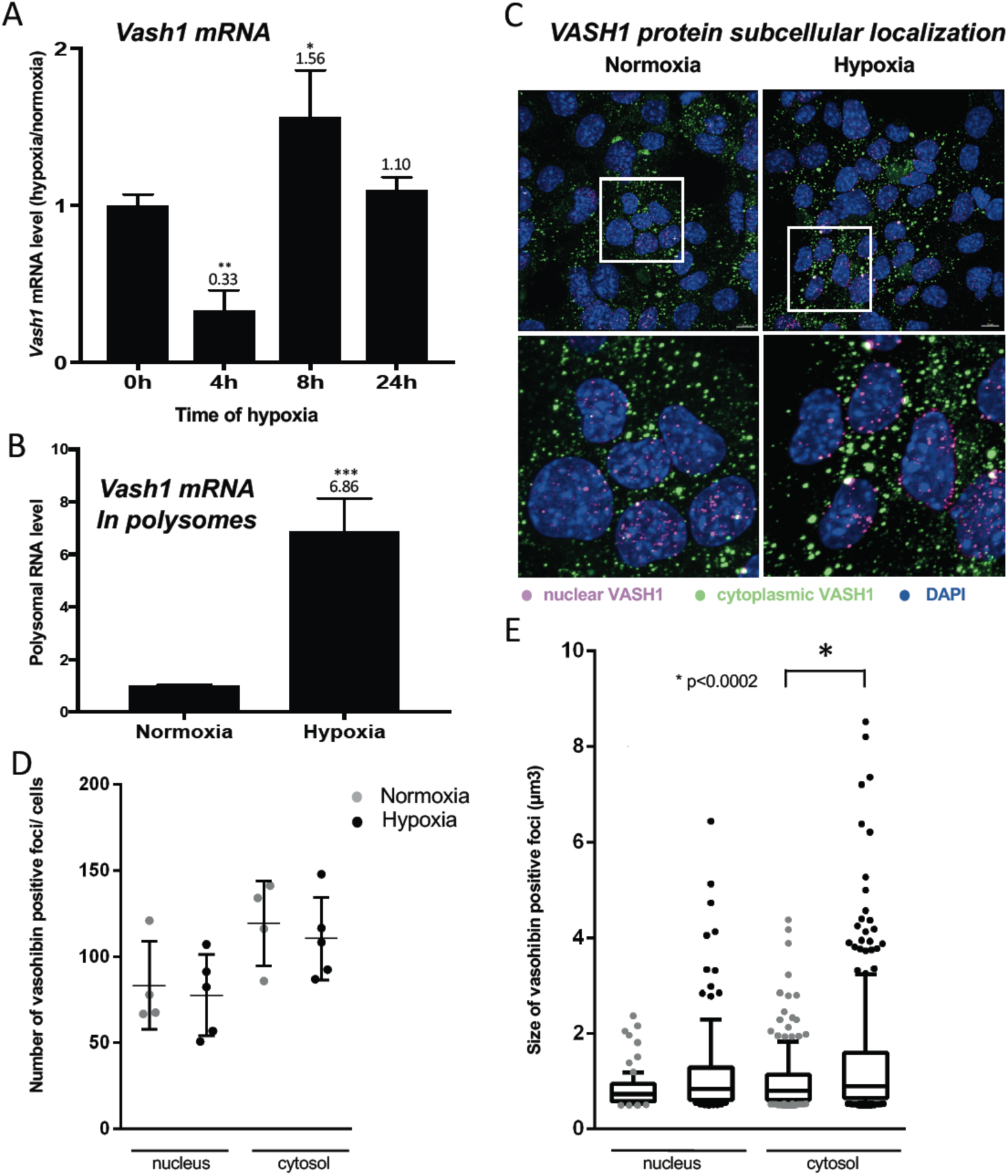
Vasohibin1 is translationally induced and nuclearized in early hypoxia. A-D VASH1 expression was analyzed by RT-qPCR in HL-1 cardiomyocytes in response to hypoxia at the transcriptome and translatome levels. Total RNA was purified from cardiomyocytes in normoxia, or submitted to 4 h, 8 h or 24 h of hypoxia (A). Polysomal RNA was purified from cardiomyocytes in normoxia or after 4 h of hypoxia (B). Histograms correspond to mean ± standard deviation of the mean, with two-tailed t-test, *p<0.05, **p<0.01, ***<0.001, compared to normoxia. Experiments have been reproduced 3 times independently and a representative triplicate experiment is shown. VASH1 was immunodetected in HL-1 cardiomyocytes in normoxia or after 4h of hypoxia (C). DAPI staining allows to detect VASH1 nuclear localization (MERGE). VASH1 foci in the nucleus are shown in purple and in the cytoplasm in green using Imaris software. The number of VASH1 foci was quantified in the nucleus and in the cytoplasm in normoxia and after 4 h of hypoxia (n = 4-5 images with a total cell number of 149 in normoxia and 178 in hypoxia) (D). Boxplots of volume of vasohibin foci in normoxia and hypoxia (E). All foci above 0,5 μm_3_ were counted. Whiskers mark the 10 % and the 90% percentiles with the mean in the center. One-way Anova with Tukey’s comparisons test was applied.

### Vasohibin1 is a new ITAF selectively active in early hypoxia

The putative ITAF function of VASH1 was assessed by a knock-down approach using an siRNA smartpool (siVASH1). Transfection of HL-1 cardiomyocytes with siVASH1 was able to knock-down VASH1 mRNA with an efficiency of 73% (Fig. 7A). The knock-down of VASH1 protein measured by capillary Western was only 59% (Fig. 7B). This moderate knock-down efficiency was probably due to the long half-life of VASH1, superior to 24h (EV Fig. 5). The effect of VASH1 knock-down was analyzed in HL-1 cells transduced with different IRES-containing bicistronic lentivectors in normoxia or after 8 h of hypoxia. In normoxia, VASH1 knock-down generated a moderate increase of activity for several IRESs (13-16%), significant for VEGFD and EMCV IRESs (Fig. 7C). In contrast, in hypoxia, VASH1 knock-down resulted in a significant decrease of FGF1, VEGFD and EMCV IRES activities, by 64%, 12% and 5%, respectively (Fig. 7D). These data showed that VASH1 behaves as an activator ITAF in hypoxia, limited to FGF1, VEGFD and EMCV IRESs, while it has an inhibitory role on the activities of these IRES in normoxia (Fig. 7C).

**Figure 7.**
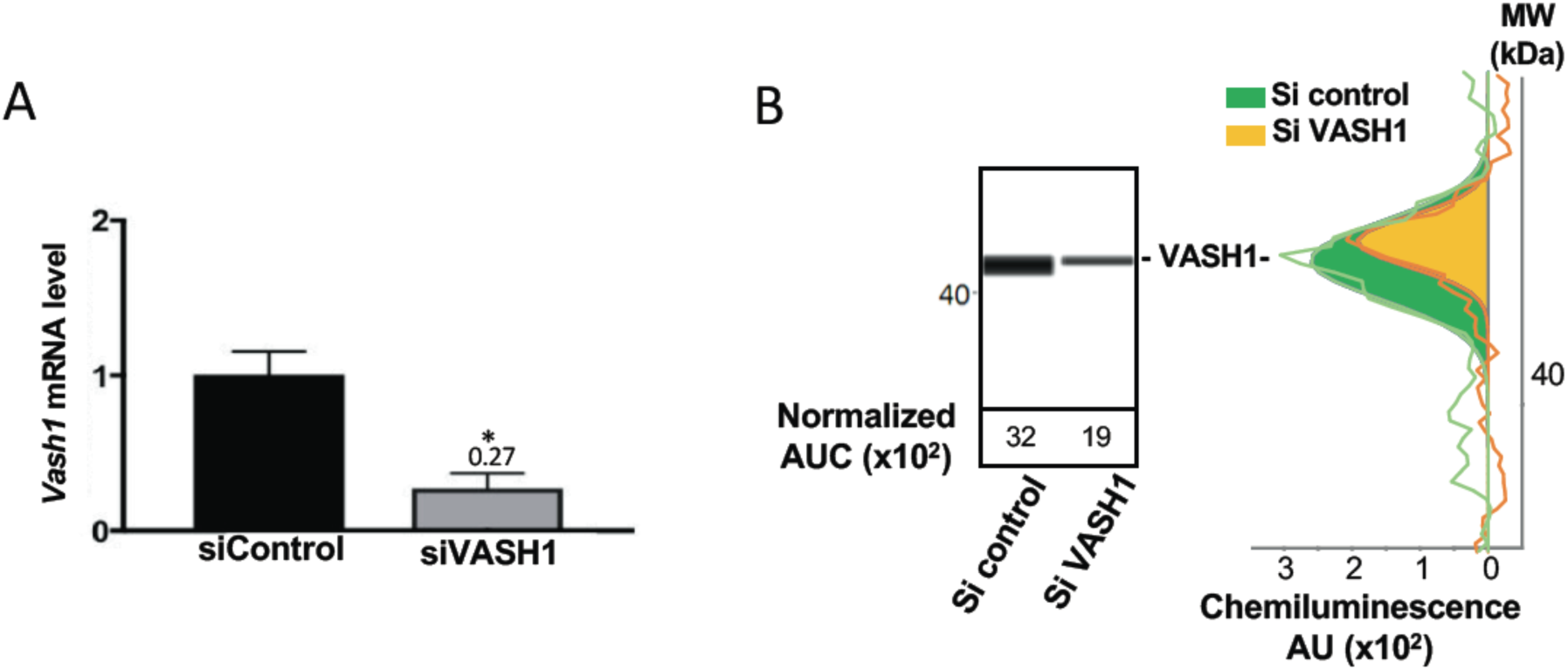
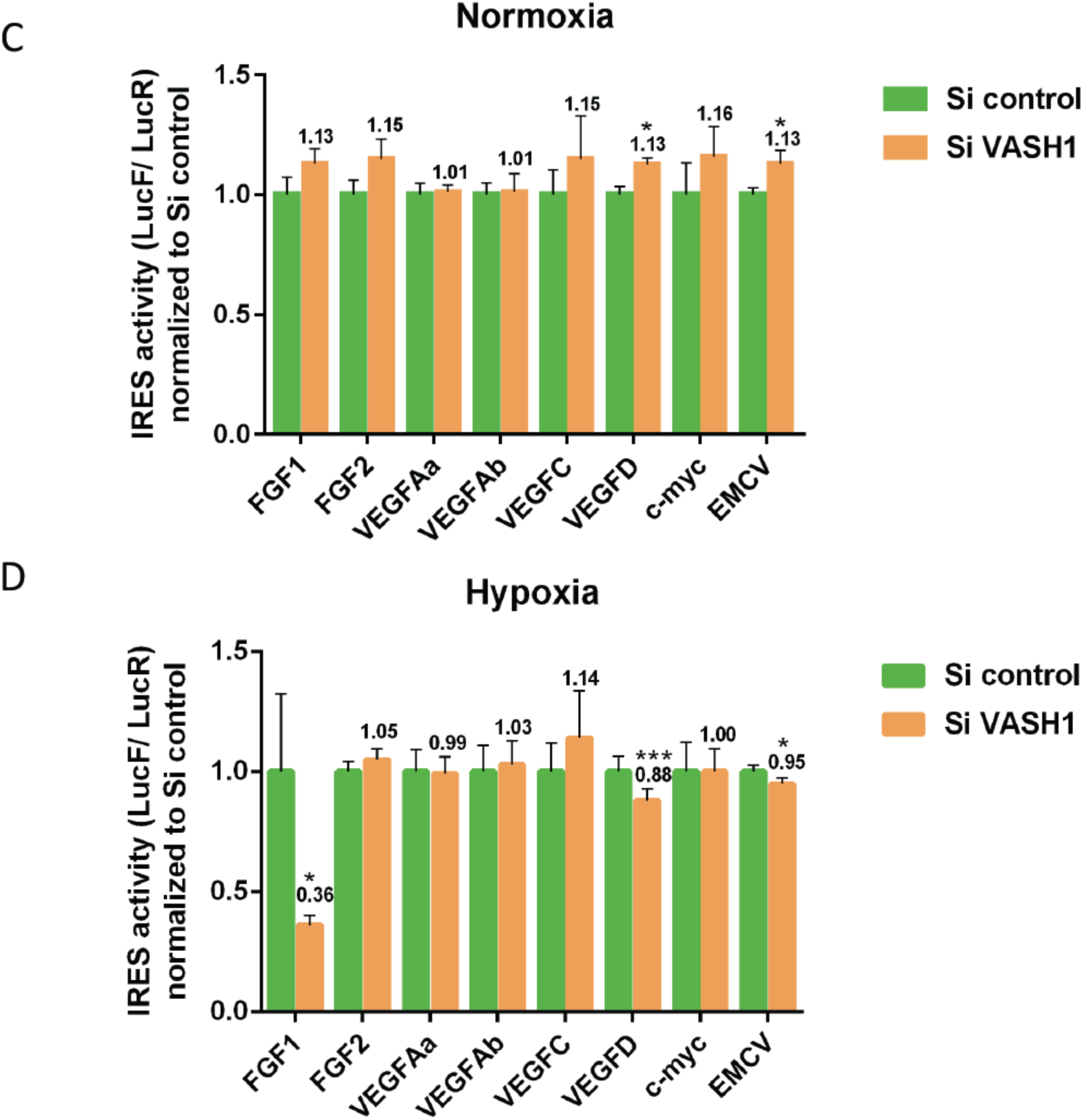
Vasohibin1 is a new ITAF selectively active in early hypoxia. A-B VASH1 knock-down was performed in HL-1 cardiomyocytes using siRNA smartpools targeting VASH1 (siVASH1) or control (siControl). VASH1 mRNA level was measured by RT-qPCR (A) and VASH1 protein expression analyzed by capillary Simple Western method using an anti-VASH1 antibody and quantified by normalization to total proteins. The experiments have been reproduced three times and representative results are shown (B). Knock-down experiment of VASH-1 performed on cardiomyocytes transduced by a set of IRES-containing lentivectors used in Fig. 4. After 8 h of hypoxia, IRES activities (LucF/LucR ratio) were measured in cell extracts. The IRES activity values have been normalized to the control siRNA. Histograms correspond to means ± standard error of the mean of the mean, with a one-tailed t-test *p<0.05, ***<0.001, compared to siControl. For each IRES the mean of three independent experiments with three biological replicates (n=9) is shown in normoxia (C) and hypoxia (D). All detailed values as well as standard deviations are presented in EV Table 5.

## DISCUSSION

The present study highlights the crucial role of translational control in cardiomyocyte response to hypoxia. Up to now, although a few genes had been described for their translational regulation by hypoxia, it was thought that most genes are transcriptionally regulated. Here we show that translational control, revealed by mRNA recruitment in polysomes during hypoxia, concerns the majority of the genes involved in angiogenesis and lymphangiogenesis. IRES-dependent translation appears as a key mechanism in this process, as we show that all the (lymph)angiogenic mRNAs known to contain an IRES are up-regulated. Furthermore, our data reveal that IRESs of angiogenic factor mRNAs are activated during early hypoxia, while non angiogenic mRNA IRESs are activated in late hypoxia. We have identified an angiogenesis- and stress-related protein, vasohibin1, as a new ITAF responsible for the activation of several, but not all IRESs, in early hypoxia.

### Translational control in tumoral versus non tumoral hypoxia

Most studies of gene expression in response to stress have been performed at the transcriptome level in tumoral cells of different origins, whereas the present study is focused on cardiomyocytes. HL-1 cardiomyocytes are immortalized but still exhibit the beating phenotype (21). Thus this cell model, although not perfectly mimicking the cardiomyocyte behavior *in vivo*, is still close to a physiological state. The strong translational response to hypoxia revealed by our data, that differs from the transcriptional response usually observed in tumor cells, may reflect mechanisms occurring in cells that are not engaged into the cell transformation process leading to cancer, or at least not too far. Indeed, HL-1 cells respond to hypoxia very early, whereas various murine or human tumor cell lines described in other reports require a longer time of hypoxia for IRES-dependent translation to be stimulated. In human breast cancer BT474 cells, VEGFA, HIF and EMCV IRESs are all activated after 24h of hypoxia (29). In murine 4T1 and LLC cells (breast and lung tumor, respectively) as well as in human CAPAN-1 pancreatic adenocarcinoma, the VEGFA and VEGFC IRESs are activated after 24 hours of hypoxia whereas the EMCV IRES is not activated (2). The same observation of late activation in 4T1 cells has been made for the FGF1 IRES, while this IRES is activated in early hypoxia in HL-1 cardiomyocytes (Godet AC & Prats AC, unpublished data)(Fig. 4). Also, the VEGFD IRES is differently regulated in HL-1 cardiomyocytes compared to 4T1 tumor cells: only heat shock, but not hypoxia, is able to activate this IRES in 4T1 cells, whereas it is activated by hypoxia in HL-1 cardiomyocytes (Fig. 4)(13). These observations suggest that many tumoral cell lines developing resistance to hypoxia are not able to govern subtle regulations of gene expression such as the waves of IRES regulation observed in HL-1 cells.

### VASH1, an ITAF of early hypoxia

We also consider the hypothesis that the important process of translational regulation observed in our study may be cardiomyocyte-specific. In such a case, IRES-dependent translation would depend on cell type specific ITAFs as well as the early response to hypoxia. These results are of great importance in regard to the acute stress response in ischemic heart that is necessary for recovery. In contrast, a delayed chronic response is known to be deleterious for heart healing (30). In agreement with this hypothesis, VASH1 expression is cell-type specific: described up to now as endothelial-specific, this protein is not expressed in tumoral cells (27). In the present study, we show that this cell-type specificity extends to cardiomyocytes. Consistent with our data, this protein has been described as a key actor of striated muscle angio-adaptation (31). VASH1 may thus have a role in the early hypoxic response in a limited number of cell types. The ITAF role of VASH1 identified here is physiologically relevant if one considers the VASH1 function in angiogenesis and stress tolerance (28). According to previous reports, VASH1 is induced during angiogenesis in endothelial cells and halts this process, while its overexpression also renders the same cells resistant to senescence and cell death induced by stress (28). Furthermore, it has been reported that VASH1 is induced after 3 hours of cell stress at the protein level but not at the transcriptional level in endothelial cells (32). This is in agreement with our observation in cardiomyocytes where VASH1, although downregulated in the transcriptome in early hypoxia, is more efficiently recruited in polysomes at the same time (Fig. 6).

It is noteworthy that VASH1 itself seems to be induced translationally by stress (Fig. 6) (32). In endothelial cells, Myashita et al report that the protein HuR upregulates VASH1 by binding to its mRNA. HuR may bind to an AU-rich element present in the 3’ untranslated region of the VASH1 mRNA. However, in other studies, HuR has also been described as an ITAF, thus it is possible that VASH1 itself may be induced by an IRES-dependent mechanism (10, 33, 34).

The anti-angiogenic function of VASH1 may appear inconsistent with its ability to activate the IRES of an angiogenic factor. However, our data also suggest that VASH1 might be an activator or an inhibitor for (lymph)angiogenic factor mRNA translation. Such a double role may explain the unique dual ability of VASH1 to inhibit angiogenesis and to promote endothelial cell survival (28, 32). This could result from the existence of different VASH1 isoforms of 44 kDa, 42 kDa, 36 kDa, 32 kDa and 27 kDa, resulting from alternative splicing and/protein processing (31, 35–37). Interestingly p42 and p27 are the main isoforms expressed in heart, while the p44 is undetectable (31, 37). One can expect that the ITAF function is carried by p42 which contains the two predicted RNA binding domains (Fig.5C, EV Fig. 4). VASH1 has been observed in both the nucleus and the cytoplasm, and no striking nucleocytoplasmic relocalization is visible in response to hypoxia, in contrast to other ITAFs such as hnRNPA1 or nucleolin which shuttle to cytoplasm upon stress (10, 13, 15, 38, 39). Interestingly, VASH1, appears as foci whose size increases in hypoxia, suggesting that it could be partly translocated to stress granules. This translocation has been reported for other ITAFs such as hnRNPA1 and polypyrimidine tract binding protein (PTB)(10, 40, 41).

### VASH1 regulates several, but not all IRESs

Although all IRESs of (lymph)angiogenic factor mRNAs are activated in early hypoxia, only FGF1 and VEGFD IRESs are regulated by VASH1. This suggests that other ITAFs are involved in activation of FGF2, VEGFA and VEGFC IRESs. Furthermore, the knock-down of VASH1 only partially silenced FGF1 and VEGFD IRES activities. This could result from the moderate efficiency of the knock-down, due to the strong stability of VASH1 protein, but it also suggests that VASH1 acts with other partners in the IRESsome. The double role of VASH1 as an activator or an inhibitor of IRES activity in hypoxia or in normoxia, respectively, also favors the hypothesis that VASH1 interacts with different partners in the IRESome. Thus, our study shows that IRESs of (lymph)angiogenic growth factor mRNAs, although they are all activated in early hypoxia, are regulated by different IRESome complexes whose composition is still to be discovered.

## MATERIALS & METHODS

### Lentivector construction and production

Bicistronic lentivectors coding for the *renilla* luciferase (LucR) and the stabilized firefly luciferase Luc+ (called LucF in the text) were constructed from the dual luciferase lentivectors described previously, which contained Luc2CP (2, 13). The LucR gene used here is a modified version of LucR where all the predicted splice donor sites have been mutated (sequence is available upon request). The cDNA sequences of the human FGF1, −2, VEGFA, -C, -D, c-myc and EMCV IRESs were introduced between the first (LucR) and the second cistron (LucF) (19, 42, 43). IRES sequences sizes are : 430 nt (FGF1), 480nt (FGF2), 302 nt (VEGFAa), 485 nt (VEGFAb), 419 nt (VEGFC), 507 nt (VEGFD), 363 nt (c-myc), 640 nt (EMCV)(2, 13, 16, 17, 19, 42). The two IRESs of the VEGFA have been used and are called VEGFAa and VEGFAb, respectively (16). The expression cassettes were inserted into the SIN lentivector pTRIP-DU3-CMV-MCS vector described previously (43). All cassettes are under the control of the cytomegalovirus (CMV) promoter.

Lentivector particles were produced using the CaCl_2_ method-based by tri-transfection with the plasmids pLvPack and pLvVSVg, CaCl_2_ and Hepes Buffered Saline (Sigma-Aldrich, Saint-Quentin-Fallavier, France), into HEK-293FT cells. Viral supernatants were harvested 48 hours after transfection, passed through 0.45 µm PVDF filters (Dominique Dutscher SAS, Brumath, France) and stored in aliquots at −80°C until use. Viral production titers were assessed on HT1080 cells with serial dilutions and scored for GFP expression by flow cytometry analysis on a BD FACSVerse (BD Biosciences, Le Pont de Claix, France).

### Cell culture, transfection and transduction

HEK-293FT cells and HT1080 cells were cultured in DMEM-GlutaMAX + Pyruvate (Life Technologies SAS, Saint-Aubin, France), supplemented with 10% fetal bovine serum (FBS), and MEM essential and non-essential amino acids (Sigma-Aldrich).

Mouse atrial HL-1 cardiomyocytes were a kind gift from Dr. William Claycomb (Department of Biochemistry & Molecular Biology, School of Medicine, New Orleans) (21). HL-1 cells were cultured in Claycomb medium containing 10% FBS, Penicillin/Streptomycin (100U/mL-100µg/mL), 0.1mM norepinephrine, and 2mM L-Glutamine. Cell culture flasks were pre-coated with a solution of 0.5% fibronectin and 0.02% gelatin 1h at 37°C (Sigma-Aldrich). To keep HL-1 phenotype, cell culture was maintained as previously described (21).

For hypoxia, cells were incubated at 37°C at 1%O_2_

HL-1 cardiomyocytes were transfected by siRNAs as follows: one day after being plated, cells were transfected with 10 nM of small interference RNAs from Dharmacon Acell SMARTpool targeting VASH1 (siVASH1) or non-targeting siRNA control (siControl), using Lipofectamine RNAiMax (Invitrogen) according to the manufacturer’s recommendations, in a media without penicillin-streptomycin and norepinephrine. Cells were incubated 72h at 37°C with siRNA (siRNA sequences are provided in EV Table 6).

For lentivector transduction, 6.10^4^ HL-1 cells were plated into each well of a 6-well plate and transduced overnight in 1 mL of transduction medium (OptiMEM-GlutaMAX, Life Technologies SAS) containing 5 µg/mL protamine sulfate in the presence of lentivectors (MOI 2). GFP-positive cells were quantified 48h later by flow cytometry analysis on a BD FACSVerse (BD Biosciences). HL-1 cells were transduced with an 80% efficiency. siRNA treatment on transduced cells was performed 72h after transduction (and after one cell passage). To achieve protein half-life measurement, HL-1 cardiomyocytes were treated with cycloheximide (InSolution CalBioChem) diluted in PBS at a final concentration of 10 µg/mL in well plates. Time-course points were taken by stopping cell cultures after 0h, 4h, 6h 8h 16h or 24h of incubation and subsequent capillary Western analysis of cell extracts.

### Reporter activity assay

For reporter lentivectors, luciferase activities *in vitro* and *in vivo* were performed using Dual-Luciferase Reporter Assay (Promega, Charbonnières-les-Bains, France). Briefly, proteins from HL-1 cells were extracted with Passive Lysis Buffer (Promega France). Quantification of bioluminescence was performed with a luminometer (Centro LB960, Berthold, Thoiry, France).

### Capillary electrophoresis

Diluted protein lysate was mixed with fluorescent master mix and heated at 95°C for 5 minutes. 3 µL of protein mix containing Protein Normalization Reagent, blocking reagent, wash buffer, target primary antibody (mouse anti-VASH-1 Abcam EPR17420 diluted 1:100; mouse anti-FGF1 Abcam EPR19989 diluted 1:25; mouse anti-P21 Santa Cruz sc-6546 (F5) diluted 1:10; rabbit anti eIF2*α* Cell Signaling Technology 9721 diluted 1:50; mouse anti-phospho-eIF2*α* Cell Signaling Technology 2103 diluted 1:50; rabbit anti-4EBP-1 Cell Signaling Technology 9452 diluted 1:50; rabbit anti-phospho-4EBP-1 Cell Signaling Technology 9451 diluted 1:50), secondary-HRP (ready to use rabbit “detection module”, DM-001), and chemiluminescent substrate were dispensed into designated wells in a manufacturer provided microplate. The plate was loaded into the instrument (Jess, Protein Simple) and proteins were drawn into individual capillaries on a 25 capillary cassette (12-230kDa)(SM-SW001). Data were analyzed on compass software provided by the manufacturer.

### RNA purification and cDNA synthesis

Total RNA extraction from HL-1 cells was performed using TRIzol reagent according to the manufacturer’s instructions (Gibco BRL, Life Technologies, NY, USA). RNA quality and quantification were assessed by a Xpose spectrophotometer (Trinean, Gentbrugge, Belgium). RNA integrity was verified with an automated electrophoresis system (Fragment Analyzer, Advanced Analytical Technologies, Paris, France).

500 ng RNA was used to synthesize cDNA using a High-Capacity cDNA Reverse Transcription Kit (Applied Biosystems, Villebon-sur-Yvette, France). Appropriate no-reverse transcription and no-template controls were included in the PCR array plate to monitor potential reagent or genomic DNA contaminations, respectively. The resulting cDNA was diluted 10 times in nuclease-free water. All reactions for the PCR array were run in biological triplicates.

### qPCR array

The DELTAgene Assay^TM^ was designed by Fluidigm Corporation (San Francisco, USA). The qPCR-array was performed on BioMark with the Fluidigm 96.96 Dynamic Array following the manufacturer’s protocol (Real-Time PCR Analysis User Guide PN 68000088). The list of primers is provided in EV Table 6. A total of 1.25 ng of cDNA was preamplified using PreAmp Master Mix (Fluidigm, PN 100-5580, 100-5581, San Francisco, USA) in the plate thermal cycler at 95°C for 2 min, 10 cycles at 95°C for 15sec and 60°C for 4 min. The preamplified cDNA was treated by endonuclease I (New England BioLabs, PN M0293L, Massachusetts, USA) to remove unincorporated primers.

The preamplified cDNA was mixed with 2x SsoFast EvaGreen Supermix (BioRad, PN 172-5211, California, USA), 50 μM of mixed forward and reverse primers and sample Loading Reagent (Fluidigm, San Francisco, USA). The sample was loaded into the Dynamic Array 96.96 chip (Fluidigm San Francisco, USA). The qPCR reactions were performed in the BioMark RT-qPCR system. Data was analyzed using the BioMark RT-qPCR Analysis Software Version 2.0.

18S rRNA was used as a reference gene and all data were normalized based on 18S rRNA level. Hprt was also assessed as a second reference gene but was not selected as its level was not stable during hypoxia. Relative quantification (RQ) of gene expression was calculated using the 2^-ΔΔCT^ method. When the RQ value was inferior to 1, the fold change was expressed as - 1/RQ. The oligonucleotide primers used are detailed in EV Table 6.

### Polysomal RNA preparation

HL-1 cells were cultured in 150-mm dishes. 15 min prior to harversting, cells were treated by cycloheximide at 100 μg/ml. Cells were washed three times in PBS cold containing 100 μg/mL cycloheximide and scraped in the PBS/cycloheximide. After centrifugation at 3,000 rpm for 2 min at 4°C, cells were lysed by 450μl hypotonic lysis buffer (5 mM Tris-HCL, pH7.5 ; 2.5 mM MgCl_2_ ; 1.5 mM KCl). Cells were centrifuged at 13,000 rpm for 5 min at 4°C, the supernatants were collected and loaded onto a 10-50% sucrose gradient. The gradients were centrifuged in a Beckman SW40Ti rotor at 39,000 rpm for 2.5 h at 4°C without brake. Fractions were collected using a Foxy JR ISCO collector and UV optical unit type 11. RNA was purified from pooled heavy fractions containing polysomes (fractions 19-27), as well as from cell lysate before gradient loading.

### Preparation of biotinylated RNA

The FGF1, VEGFA or EMCV IRESs was cloned in pSCB-A-amp/kan plasmid (Agilent) downstream from the T7 sequence. The plasmid were linearized and *in vitro* transcription was performed with MEGAscript T7 kit (Ambion), according to the manufacturer’s protocol, in the presence of Biotin-16-UTP at 1 mM (Roche), as previously described (26). The synthesized RNA was purified using RNeasy kit (Qiagen).

### BIA-MS experiments

BIA-MS studies based on surface plasmonic resonance (SPR) technology were performed on BIAcore T200 optical biosensor instrument (GE Healthcare), as described previously (**13, 26**). Immobilization of biotinylated IRES RNAs was performed on a streptavidin-coated (SA) sensorchip in HBS-EP buffer (10 mM Hepes pH 7.4, 150 mM NaCl, 3 mM EDTA, 0.005% surfactant P20) (GE Healthcare). All immobilization steps were performed at a flow rate of 2 *µ*l/min with a final concentration of 100 *µ*g/ml.

Binding analyses were performed with normoxic or hypoxic cell protein extracts at 100 *µ*g/ml over the immobilized IRES RNA surface for 120 sec at a flow rate of 30 *µ*l/min. The channel (Fc1) was used as a reference surface for non-specific binding measurements. The recovery wizard was used to recover selected proteins from cell protein extracts. This step was carried out with 0.1% SDS. Five recovery procedures were performed to get enough amounts of proteins for MS identification.

Eluted protein samples from BIA experiment were digested *in gel* with 1 *µ*g of trypsin (sequence grade, Promega) at 37°C OVN. Peptides were then subjected to LC-MS/MS analysis. The peptides mixtures were loaded on a YMC-Triart C18 150×300 *µ*m capillary column (particle diameter 3 *µ*m) connected to a RS3000 Dionex HPLC system. The run length gradient (acetonitrile and water) was 30 minutes. Then, on the AB Sciex 5600+ mass spectrometer, data were acquired with a data dependent analysis. Data were then loaded on Mascot software (Matrix Science) that attributes peptide interpretations to MS/MS recorded scans. The higher the score, the lower the probability of false positive (a score of 20 corresponds to a 5% probability of false positive).

### Surface Plasmon Resonance assays

For kinetic analysis, immobilization of biotinylated FGF1 IRES RNA was performed on a streptavidin-coated (SA) sensorchip in HBS-EP buffer (10 mM Hepes pH 7.4, 150 mM NaCl, 3mM EDTA, 0.005% surfactant P20) (GE Healthcare). Immobilization step was performed at a flow rate of 2 µl/min with a final concentration of 100 µg/ml. Total amount of immobilized FGF1 IRES RNA was 1500 RU.

Binding analyses were performed with recombinant protein VASH1 (Abnova H00022846-P01) at 100 µg/ml over the immobilized FGF1. This recombinant VASH1 contains the 27 kDa N-terminal part of the protein coupled to glutathione S-transferase. The channel (Fc1) was used as a reference surface for non-specific binding measurements.

A Single-Cycle Kinetics (SCK) analysis to determine association, dissociation and affinity constants (ka, kd, and KD respectively) was carried out by injecting different protein concentrations (16.25 nM-300 nM). Binding parameters were obtained by fitting the overlaid sensorgrams with the 1:1. Langmuir binding model of the BIAevaluation software version 3.0.

### Immunocytology

Cells were plated on glass coverslip and incubated for 4 h of normoxia or hypoxia. They were fixed with cold methanol at −20°C during 5 min, washed 3 times with PBS and permeabilized 1 min with 0.1% Triton. Then, cells were incubated 5 min with blocking solution (1% FBS, 0.5% BSA) and 30 min with anti-VASH1 antibody (1/50; abcam ab176114) and Alexa 488 conjugated anti-mouse secondary antibody. Images were acquired with LSM780 Zeiss confocal microscope, camera lens x60 with Z acquisition of 0.36 μM. A single plan is shown Fig. 6C.

Imaris software was used to represent vasohibin staining in Figure 6C. To differentiate vasohibin in the nucleus and cytoplasm, nucleus was delimitated with Dapi staining and all Vasohibin foci in the nucleus are shown in purple and in the cytoplasm in green.

Using imaris software, the mean of vasohibin foci was counted and the volume of vasohibin foci was quantified, a threshold was applied and all particles above 0,5 μm^3^ was selected and quantified.

### Statistical analysis

All statistical analyses were performed using one-way Anova with Tukey’s comparisons test or one-tailed Student’s t-test and are expressed as mean +-standard deviation, *p<0.05, **p<0.01, ***<0.001, ****<0.0001.

## ACKNOWLEDGMENTS

Our thanks go to J.J. Maoret and F. Martin from the Inserm UMR1048 GeT-TQ plateau of the GeT plateform Genotoul (Toulouse), F. Lopez and L. Tonini from the proteomic platform genotoul (Toulouse), J. Iacovoni from the Inserm UMR 1048 bioinformatics plateau, as well as L. van den Berghe and C. Segura from the Inserm UMR1037 vectorology plateau (Toulouse) and A. Lucas from the We-Met Functional Biochemistry Facility (Toulouse). We also thank V. Poinsot for helpful discussion and W. Claycomb for providing HL-1 cells.

This work was supported by Région Midi-Pyrénées, Association Française contre les Myopathies (AFM-Téléthon), Association pour la Recherche sur le Cancer (ARC), European funding (REFBIO), Fondation Toulouse Cancer Santé and Agence Nationale pour la Recherche AAPG2018-RIBOCARD. F.H. had fellowships from the Région Midi-Pyrénées and from the Ligue Nationale Contre le Cancer (LNCC). E.R.G. had a fellowship from AFM-Telethon. A.C. Godet had a fellowship from LNCC.

---

**Expanded View Figure 1.**
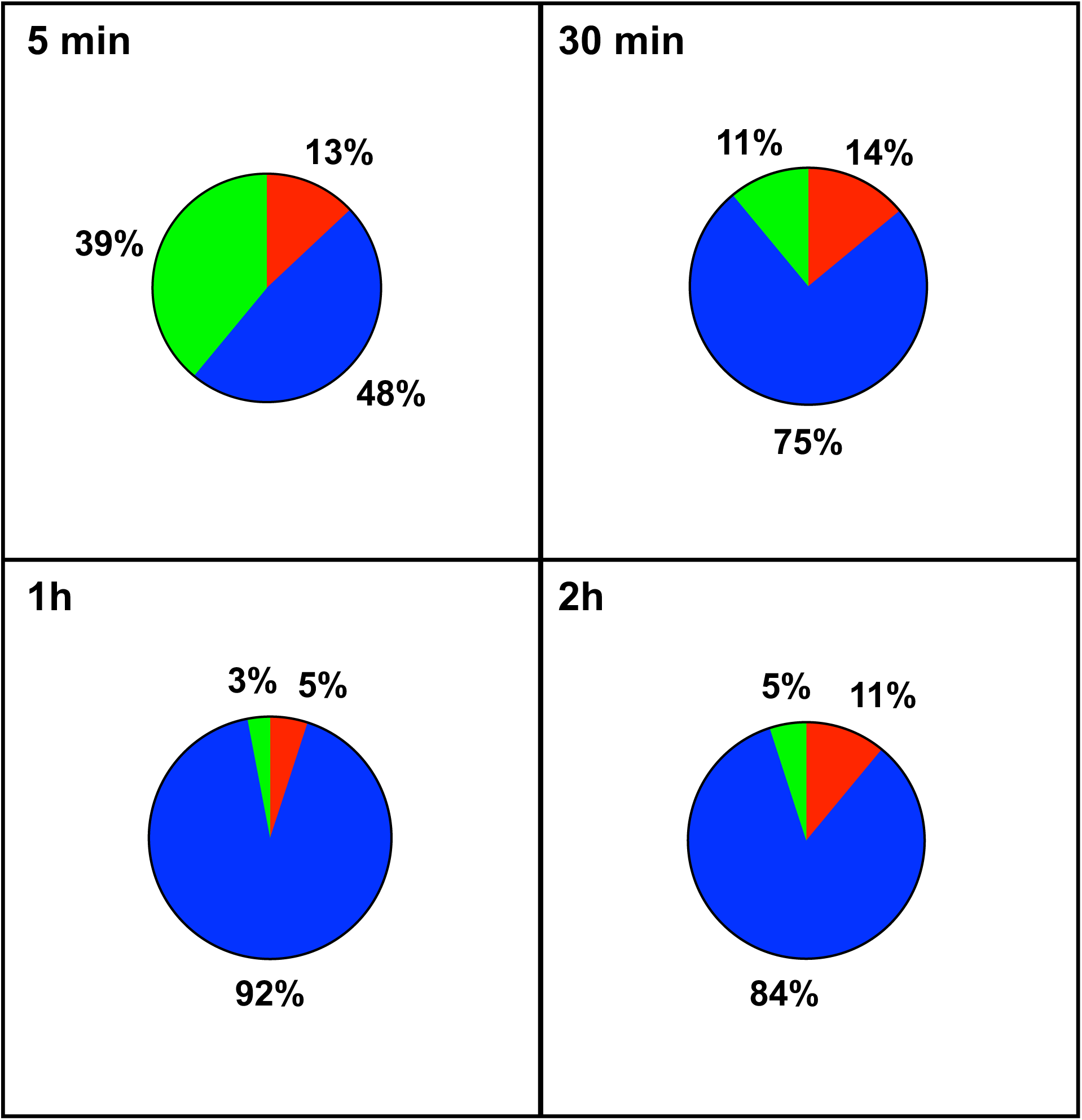
Transcriptome of IRES-containing mRNAs in hypoxic cardiomyocytes. Total RNA was purified from HL-1 cardiomyocytes submitted to increasing times from 5 min to 24 h of hypoxia at 1% O_2_, as well as from normoxic cardiomyocytes as a control. cDNA was synthesized and used for a Fluidigm deltagene PCR array dedicated to genes related to (lymph)angiogenesis or stress (EV Table 6). Relative quantification (RQ) of gene expression during hypoxia was calculated using the 2^-ΔΔCT^ method with normalization to 18S and to normoxia. The percentage of repressed (red), induced (green) and non-regulated (blue) mRNAs is shown for the earlier times of the kinetics. The later times are shown in Fig. 1. The detailed values for all the times of the kinetics are presented in EV Table 1.

**Expanded View Figure 2.**
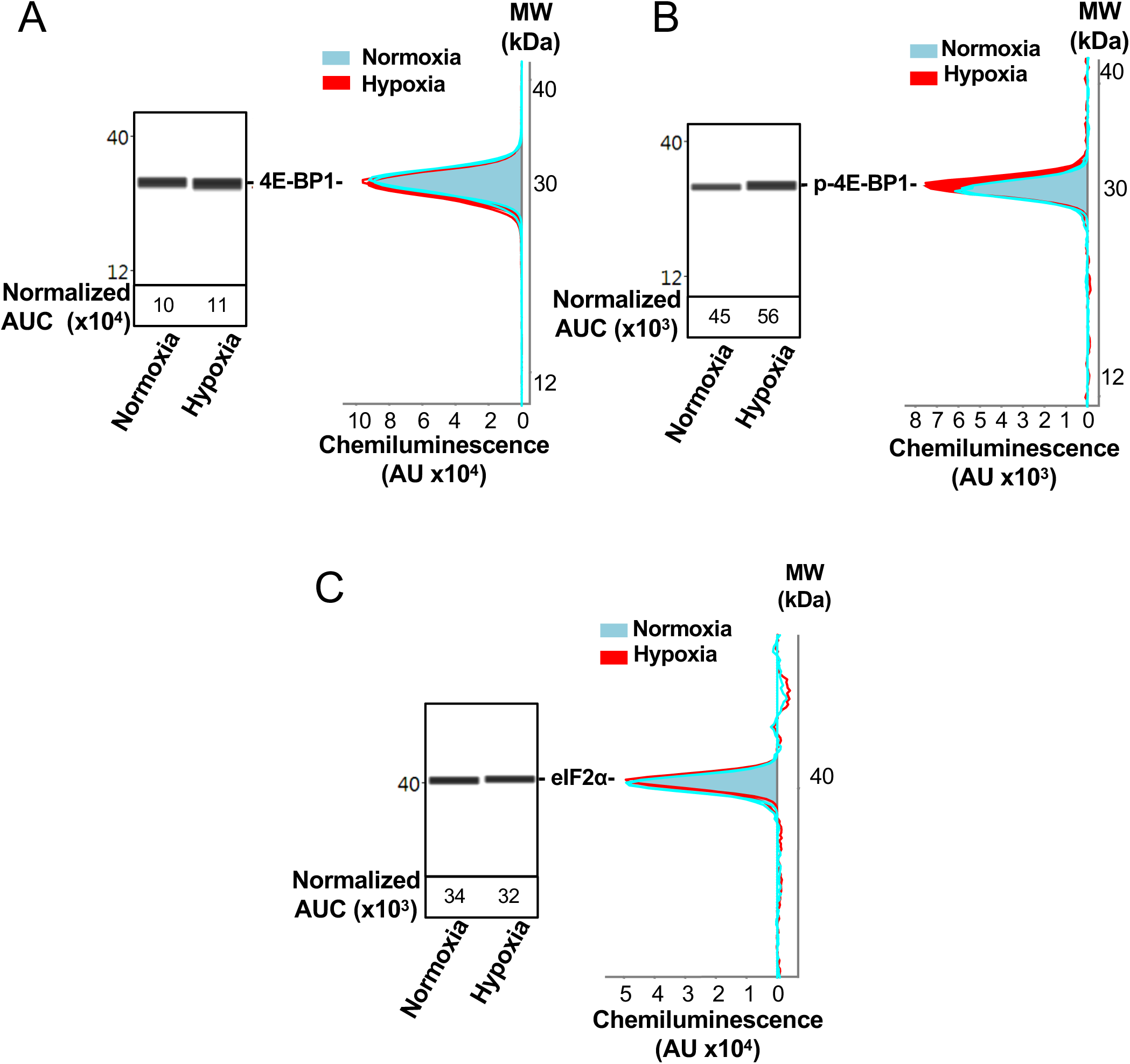
Capillary electrophoresis immunodetection of 4E-BP1 and eIF2α. A-C 4E-BP1 expression (A) and phosphorylation (B), as well as eIF2*α* expression (C) in normoxia and hypoxia (8 h) were analysed and quantified by capillary Simple Western, as described in Mat. & Meth. The quantified values, expressed in arbitrary units of luminescence (AUC) are normalized to total proteins. Analysis of eIF2*α* phosphorylation is shown in Figure 2.

**Expanded View Figure 3.**
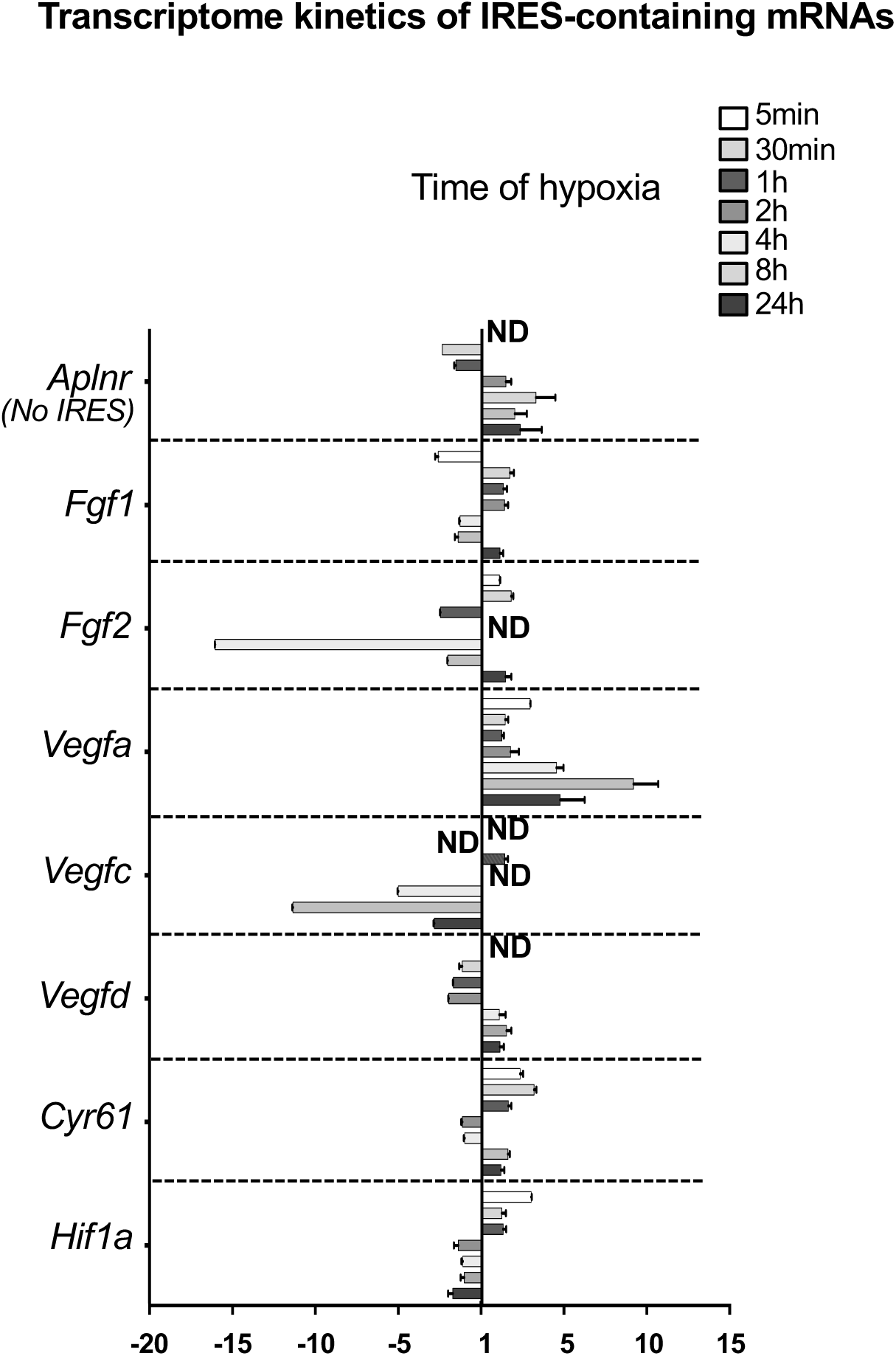
Transcriptome of IRES-containing mRNAs in hypoxic cardiomyocytes. RQ values for IRES-containing mRNA transcriptome kinetics extracted from the PCR arrays shown in EV Table 1. The gene *Aplnr* (apelin receptor) was chosen as a control without an IRES.

**Expanded View Figure 4.**
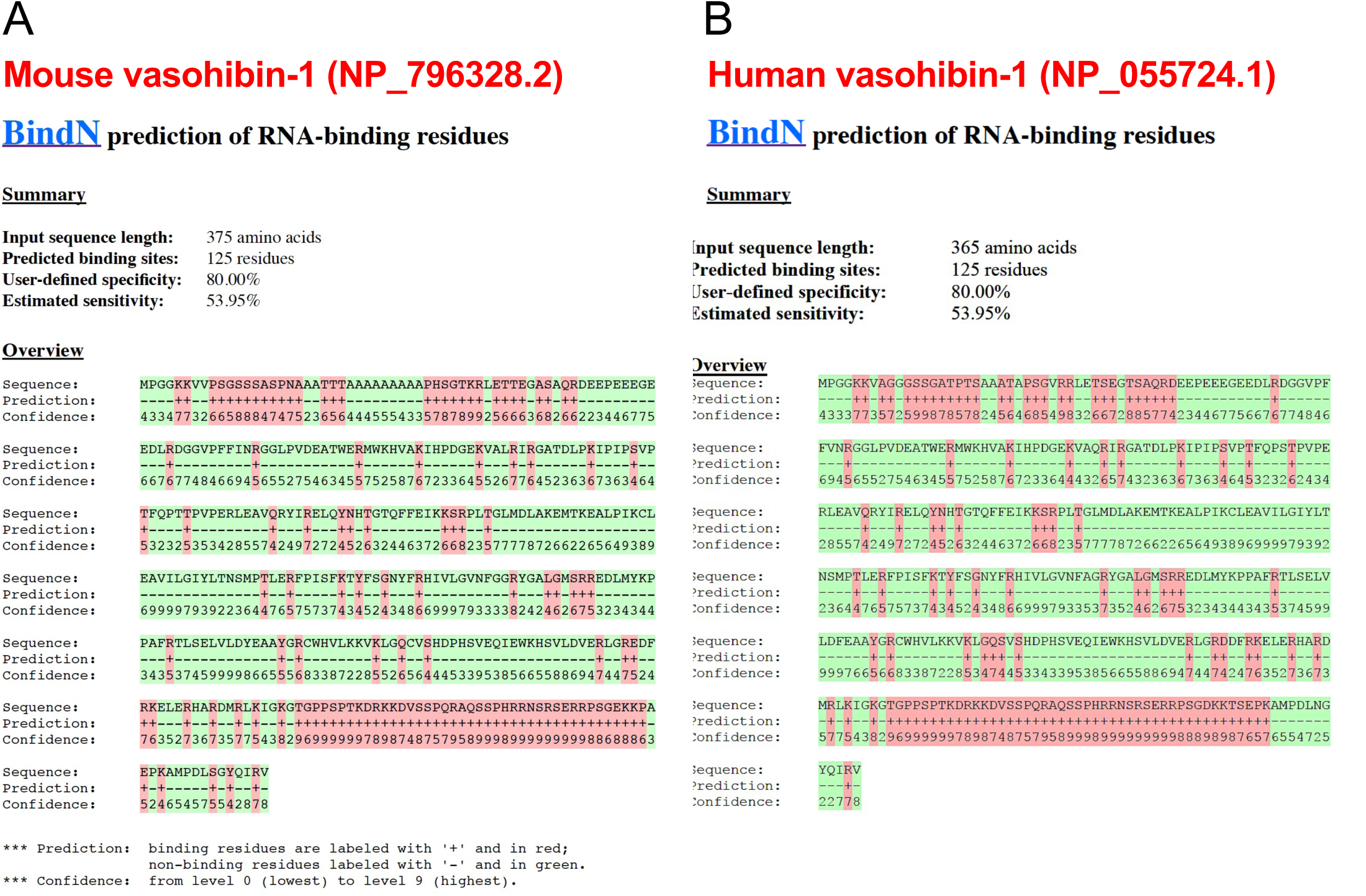
Conservation of predicted RNA binding domains in mouse and human vasohibin-1. A-B RNA binding domains in mouse (A) and human (B) VASH1 proteins were predicted using BindN software (https://omictools.com/bindn-2-tool)

**Expanded View Figure 5.**
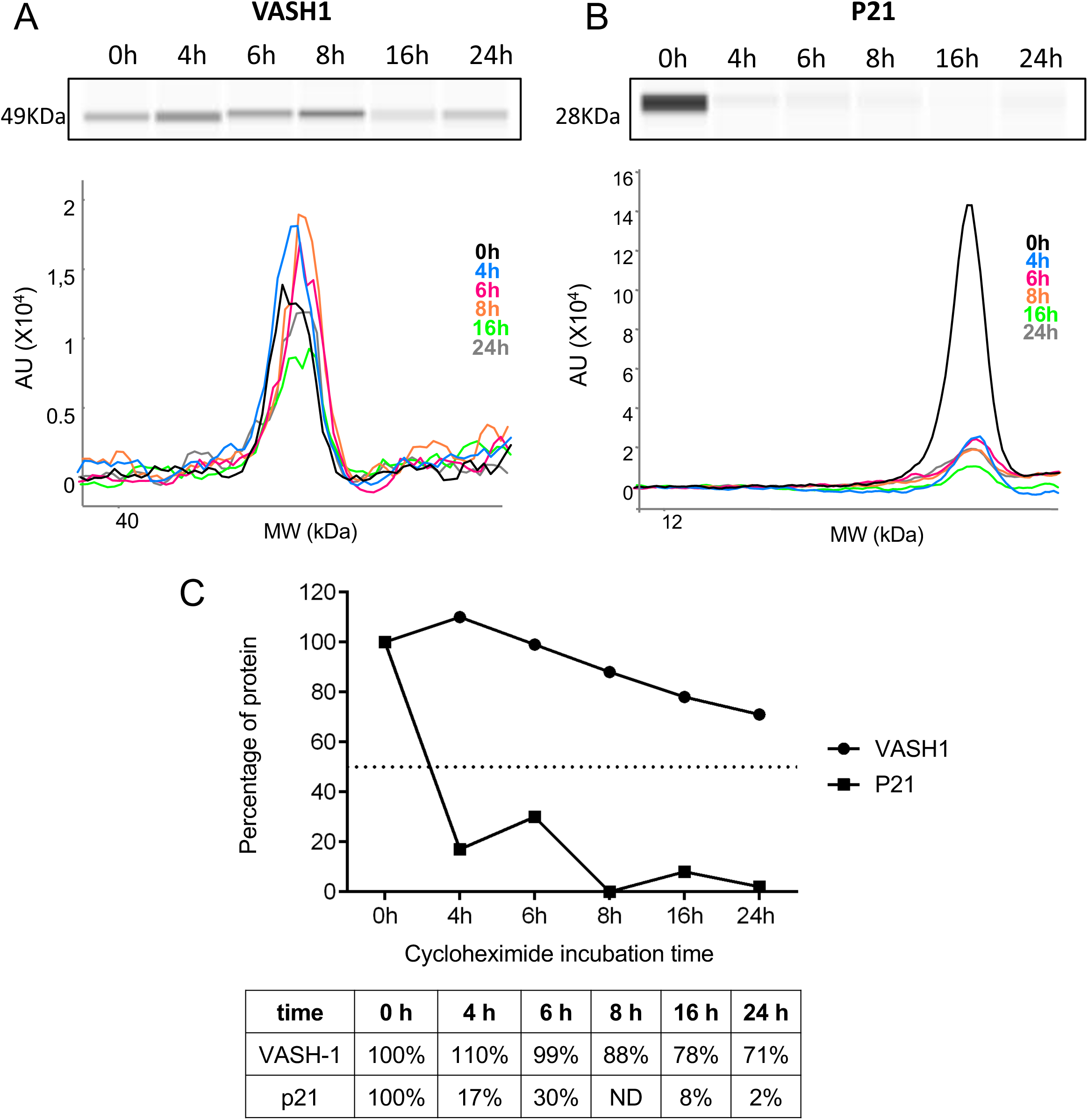
VASH1 half life is superior to 24h. A-C VASH1 half-life determination experiment were performed by blocking protein synthesis with cycloheximide at 10 µg/mL, with time-course points at 0 h, 4 h, 6 h, 8 h, 16 h and 24 h. VASH1 (A) and P21 (B) protein stability was measured by capillary Simple Western with normalization to 0 h time-course point. P21 was used as a control for its short half-life (C).

**EV Table 1.** Transcriptome of (lymph)angiogenic factor genes in hypoxic HL-1 cardiomyocytes. Total RNA was purified from HL-1 cardiomyocytes submitted to increasing times from 5 min to 24 h of hypoxia at 1% O2, as well as from normoxic cardiomyocytes as a control. cDNA was synthesized and used for a Fluidigm deltagene PCR array dedicated to genes related to (lymph)angiogenesis or stress (EV Table 6). Relative quantification (RQ) of gene expression in hypoxia was calculated using the 2^-ΔΔCT^ method with normalization to 18S and to normoxia. Standard deviation is indicated. When the RQ value is inferior to 1, the fold change is expressed as −1/RQ. ND means “non detected”.

Hantelys et al, EV Table 1
| Hypoxia time | 5min | 30min | 1h | 2h | 4h | 8h | 24h |
| --- | --- | --- | --- | --- | --- | --- | --- |
| Gene name | RQ | RQ | RQ | RQ | RQ | RQ | RQ |
| *Akt1* | 1.60 ±0.13 | 1.30 ±0.06 | 1.20 ±0.15 | 1.36 ±0.25 | -1.10 ±0.09 | 1.40 ±0.13 | 1.00 ±0.32 |
| *Ang* | 2.65 ±0.25 | 1.23 ±0.36 | -1.04 ±0.12 | 1.24 ±0.04 | 1.24 ±0.12 | 1.09 ±0.08 | 1.26 ±0.28 |
| *Angpt1* | 5.51 ±0.17 | 1.20 ±0.30 | 1.19 ±0.07 | 1.90 ±0.49 | 2.43 ±0.36 | 1.56 ±0.43 | -2.27 ±0.06 |
| *Angptl4* | 1.03 ±0.02 | -3.14 ±0.09 | 1.65 ±0.54 | 1.60 ±0.46 | 1.56 ±0.67 | 3.50 ±1.07 | -1.37 ±0.35 |
| *Anpep* | ND | -3.26 ±0.29 | 1.65 ±0.01 | -8.05 ±0.06 | -1.04 ±0.14 | 1.62 ±0.62 | 1.03 ±0.43 |
| *Apln (Apelin)* | 2.11 | -19.21 ±0.01 | -6.17 ±0.05 | -29.04 ±0.01 | 36.77 ±5.59 | 109.98 ±6.98 | 335.72 ±57.52 |
| *Aplnr* | ND | -2.36 ±0.03 | -1.57 ±0.10 | 1.44 ±0.32 | 3.25 ±1.70 | 1.98 ±0.72 | 2.31 ±1.29 |
| *Atp2a2 (SERCA2)* | 1.46 ±0.08 | 1.21 ±0.13 | 1.26 ±0.14 | 1.50 ±0.01 | 1.20 ±0.06 | 1.84 ±0.18 | 1.48 ±0.32 |
| *Bai1* | -1.52 ±0.19 | 2.90 ±0.91 | 1.17 ±0.06 | 1.65 ±0.52 | 1.36 ±0.13 | 2.05 ±0.60 | 1.04 ±0.30 |
| *Ccl2* | ND | -1.09 ±0.49 | 1.76 ±0.46 | -2.76 ±0.16 | 1.57 ±1.01 | 2.99 ±1.13 | 1.23 |
| *Ccl21a* | ND | -32.39 ±0.01 | 2.97 ±0.85 | -1.90 ±0.03 | 2.65 ±2.78 | -2.27 ±0.21 | 1.32 ±0.39 |
| *Col18a1* | 1.66 ±0.71 | 1.58 ±0.39 | 1.14 ±0.14 | 1.45 ±0.04 | -1.69 ±0.09 | 1.82 ±0.43 | 1.57 ±0.43 |
| *Col4a3* | 19.64 ±0.07 | 1.52 ±0.19 | 1.20 ±0.21 | 1.31 ±0.04 | 1.00 ±0.03 | 1.45 ±0.30 | 4.40 ±0.80 |
| *Ctgf* | 1.29 ±0.09 | 1.67 ±0.26 | 1.16 ±0.06 | 1.30 ±0.13 | 1.06 ±0.13 | 2.05 ±0.30 | 3.66 ±0.76 |
| *Cxcl1* | 1.32 ±0.64 | 1.11 ±0.27 | 1.67 ±0.19 | 1.58 ±0.12 | 1.01 ±0.09 | -1.43 ±0.08 | -2.04 ±0.13 |
| *Cxcl10* | 1.37 ±0.86 | 1.36 ±0.29 | 1.11 ±0.14 | 1.33 ±0.25 | 1.05 ±0.09 | -1.02 ±0.13 | -1.08 ±0.23 |
| *Cyr61* | 2.30 ±0.18 | 3.14 ±0.14 | 1.59 ±0.18 | -1.18 ±0.07 | -1.03 ±0.06 | 1.56 ±0.12 | 1.15 ±0.21 |
| *Edn1* | -2.43 ±0.28 | 2.93 ±0.31 | 1.72 ±0.32 | -1.4 ±0.24 | -1.39 ±0.06 | -1.33 ±0.09 | 1.04 ±0.26 |
| *Efna1* | 2.08 ±0.15 | 1.23 ±0.10 | -1.52 ±0.12 | -1.16 ±0.19 | 1.25 ±0.09 | 1.17 ±0.05 | 1.35 ±0.33 |
| *Efnb2* | 1.74 ±0.14 | 1.55 ±0.25 | 1.14 ±0.23 | -1.03 ±0.18 | 1.57 ±0.18 | 3.71 ±0.61 | 1.65 ±0.42 |
| *Egf* | ND | 1.35 ±0.14 | 1.32 ±0.30 | 1.56 ±0.44 | 1.56 ±0.11 | 2.14 ±0.53 | -1.61 ±0.35 |
| *Eng* | 2.96 ±0.94 | 1.55 ±0.39 | 1.18 ±0.19 | -1.02 ±0.14 | -1.11 ±0.15 | 2.27 ±0.33 | 1.65 ±0.34 |
| *Ephb4* | 2.22 ±0.13 | 1.67 ±0.16 | -1.03 ±0.14 | 1.10 ±0.13 | 1.01 ±0.12 | 1.45 ±0.09 | 1.57 ±0.21 |
| *ErbB2 (Her2)* | 1.87 ±0.19 | 1.85 ±0.26 | -1.01 ±0.18 | 1.16 ±0.18 | 1.32 ±0.11 | 1.00 ±0.18 | 1.34 ±0.09 |
| *F3* | 1.12 ±0.94 | 1.45 ±0.18 | 1.28 ±0.11 | -1.57 ±0.23 | -1.23 ±0.07 | -1.09 ±0.10 | -1.56 ±0.21 |
| *Fgf1 (aFGF)* | -2.63 ±0.18 | 1.69 ±0.23 | 1.29 ±0.22 | 1.37 ±0.21 | -1.32 ±0.05 | -1.43 ±0.20 | 1.08 ±0.20 |
| *Fgf2 (bFGF)* | 1.06 ±0.04 | 1.76 ±0.13 | -2.5 ±0.03 | ND | -16.05 ±0.02 | -2.05 ±0.07 | 1.43 ±0.35 |
| *Fgfr3* | 1.41 ±0.07 | 1.34 ±0.27 | -1.08 ±0.12 | 1.38 ±0.31 | 1.47 ±0.27 | 3.93 ±0.55 | 2.69 ±0.39 |

| *Hypoxia time* | 5min | 30min | 1h | 2h | 4h | 8h | 24h |
| --- | --- | --- | --- | --- | --- | --- | --- |
| *Gene name* | RQ | RQ | RQ | RQ | RQ | RQ | RQ |
| *Fn1* | 3.30 ±0.07 | 1.70 ±0.15 | -1.02 ±0.17 | 1.16 ±0.01 | -1.25 ±0.06 | 2.45 ±0.36 | 2.61 ±0.49 |
| *Hif1a* | 2.97 ±0.03 | 1.20 ±0.24 | 1.28 ±0.18 | -1.42 ±0.26 | -1.15 ±0.08 | -1.06 ±0.21 | -1.75 ±0.22 |
| *Hnrnpm* | 1.55 ±0.18 | 1.17 ±0.18 | 1.54 ±0.15 | 2.26 ±0.20 | 1.86 ±0.25 | 3.86 ±1.33 | 1.23 ±0.26 |
| *Hpse* | ND | 3.23 ±1.07 | 1.25 ±0.34 | 1.64 ±0.16 | 1.56 ±0.14 | 3.52 ±1.07 | 11.04 ±1.95 |
| *Id1* | -1.61 ±1.12 | 1.58 ±0.18 | 2.63 ±0.50 | 3.01 ±1.27 | 4.23 ±0.19 | 2.01 ±0.40 | 1.79 ±0.64 |
| *Ifna1* | ND | 1.72 ±0.17 | -1.09 ±0.06 | 1.73 ±0.07 | 2.19 ±0.33 | 3.11 ±0.25 | 1.31 ±0.49 |
| *Igf1* | 10.07 ±0.59 | 1.54 ±0.22 | 1.42 ±0.15 | 1.06 ±0.04 | 1.10 ±0.22 | -1.25 ±0.14 | -2.86 ±0.14 |
| *Igf1r* | ND | ND | ND | ND | -3.17 ±0.08 | 2.37 ±0.20 | 5.14 ±0.32 |
| *Itgav* | ND | 1.19 ±0.11 | -1.36 ±0.14 | 1.33 ±0.29 | 1.33 ±0.12 | 3.41 ±0.56 | 2.82 ±0.64 |
| *Itgb3* | -4.33 ±0.30 | 1.34 ±0.19 | -1.35 ±0.02 | 1.04 ±0.20 | 1.03 ±0.20 | 1.42 ±0.34 | -1.37 ±0.10 |
| *Jag1* | 2.13 ±0.09 | 1.80 ±0.24 | 1.03 ±0.13 | 1.66 ±0.46 | 1.46 ±0.06 | 2.26 ±0.37 | 1.76 ±0.21 |
| *Mdk* | 1.14 ±0.21 | 2.45 ±0.17 | -1.07 ±0.17 | -1.27 ±0.00 | 1.16 ±0.32 | 1.52 ±0.38 | -1.16 ±0.46 |
| *Mmp14* | -1.77 ±0.03 | 1.30 ±0.22 | 1.02 ±0.06 | -1.16 ±0.17 | 1.06 ±0.11 | 1.70 ±0.24 | 1.95 ±0.15 |
| *Mmp2* | 1.46 ±0.18 | 1.96 ±0.50 | 1.20 ±0.05 | 1.68 ±0.44 | 1.14 ±0.33 | 1.94 ±0.66 | -1.23 ±0.37 |
| *Neat1* | 4.03 ±0.48 | -1.17 ±0.23 | -1.59 ±0.05 | 2.65 ±0.00 | 1.89 ±0.15 | 2.96 ±1.02 | -1.16 ±0.29 |
| *Nos3* | 9.29 | -4.01 ±0.10 | 2.00 ±0.54 | -2.58 ±0.26 | -1.30 ±0.14 | -1.02 ±0.40 | -1.32 ±0.19 |
| *Nrp1* | 2.46 ±0.06 | 1.32 ±0.17 | 1.04 ±0.15 | -1.25 ±0.25 | -1.10 ±0.07 | 1.47 ±0.25 | 1.10 ±0.16 |
| *Nrp2* | 2.50 ±0.07 | 1.55 ±0.05 | 1.02 ±0.12 | 1.02 ±0.12 | 1.14 ±0.19 | 1.72 ±0.32 | 1.83 ±0.39 |
| *P54nrb* | 2.96 ±0.09 | 1.58 ±0.42 | 1.21 ±0.20 | 1.42 ±0.29 | 1.22 ±0.13 | 1.66 ±0.30 | 1.11 ±0.23 |
| *PAI1 (SerpinE1)* | -1.73 ±0.12 | 2.18 ±0.54 | 1.46 ±0.15 | 1.74 ±0.65 | 5.75 ±0.57 | 15.76 ±2.01 | 13.18 ±1.02 |
| *Pdgfa* | 3.65 ±0.43 | 1.45 ±0.18 | 1.29 ±0.26 | 1.59 ±0.36 | 1.71 ±0.11 | 1.97 ±0.35 | 1.54 ±0.34 |
| *Pecam1* | 1.48 | -11.55 ±0.02 | -1.09 ±0.24 | 2.86 ±1.08 | 2.09 ±0.77 | 2.70 ±1.72 | 2.60 ±0.86 |
| *Pf4* | ND | -39.36 ±0.02 | -1.79 ±0.33 | -2.20 ±0.18 | 1.80 ±0.40 | 1.48 ±0.55 | 1.01 ±0.28 |
| *Pgf* | ND | 2.17 ±0.14 | -1.36 ±0.19 | -7.89 ±0.09 | 4.47 ±0.89 | 1.85 ±0.29 | 3.22 ±0.85 |
| *Plau (upa)* | 1.13 ±0.29 | -1.04 ±0.24 | -1.36 ±0.16 | -1.31 ±0.13 | 1.25 ±0.30 | 1.58 ±0.26 | 1.24 ±0.24 |
| *Plg* | ND | -2.71 ±0.26 | -1.05 | 2.45 ±0.63 | -1.35 ±0.19 | -3.33 ±0.16 | -2.94 ±0.25 |
| *Prox1* | 7.13 ±0.20 | 1.48 ±0.23 | 1.26 ±0.14 | -1.13 ±0.14 | -1.08 ±0.13 | 1.27 ±0.29 | -1.72 ±0.07 |

| **Hypoxia time** | **5min** | **30min** | **1h** | **2h** | **4h** | **8h** | **24h** |
| --- | --- | --- | --- | --- | --- | --- | --- |
| **Gene name** | **RQ** | **RQ** | **RQ** | **RQ** | **RQ** | **RQ** | **RQ** |
| *Psf/Sfpq* | 1.45 ±0.17 | 1.37 ±0.09 | 1.30 ±0.19 | 1.22 ±0.08 | 1.10 ±0.15 | 1.55 ±0.59 | -1.43 ±0.16 |
| *Pspc1* | 1.00 ±0.80 | 1.34 ±0.11 | 1.50 ±0.01 | 1.64 ±0.58 | 1.67 ±0.19 | 1.37 ±0.31 | 1.59 ±0.23 |
| *SerpinF1* | ND | -31.89 ±0.02 | -9.09 ±0.01 | -1.91 ±0.33 | -2.38 ±0.11 | -1.14 ±0.17 | 1.02 ±0.37 |
| *Sphk1* | ND | 1.91 ±0.80 | 1.70 ±0.04 | -1.51 ±0.19 | 1.30 ±0.11 | 2.78 ±1.20 | 1.17 ±0.43 |
| *Tek* | 1.97 ±1.55 | 1.98 ±0.23 | 1.55 ±0.41 | 1.48 ±0.20 | 1.24 ±0.14 | -1.25 ±0.09 | 1.04 ±0.13 |
| *Tgfa* | ND | -2.01 ±0.13 | -7.94 ±0.05 | -6.9 ±0.03 | 2.91 ±0.63 | 3.50 ±1.29 | 4.05 ±1.71 |
| *Tgfb1* | -5.26 ±0.02 | 1.22 ±0.23 | -1.45 ±0.14 | 1.03 ±0.10 | -1.02 ±0.22 | 1.45 ±0.24 | 1.05 ±0.25 |
| *Tgfb2* | -2.45 ±0.08 | 1.50 ±0.31 | 1.22 ±0.07 | 1.37 ±0.12 | 1.78 ±0.24 | 2.00 ±0.39 | 1.43 ±0.40 |
| *Tgfbr1* | -2.03 ±0.57 | 1.33 ±0.16 | 1.28 ±0.25 | 1.48 ±0.14 | 1.86 ±0.34 | 2.23 ±0.77 | -1.27 ±0.16 |
| *Thbs1* | 2.65 ±0.09 | 1.88 ±0.12 | 1.19 ±0.09 | 1.47 ±0.33 | 1.12 ±0.07 | 1.30 ±0.22 | 1.44 ±0.08 |
| *Thbs2* | 2.15 ±0.12 | 1.52 ±0.10 | 1.03 ±0.16 | 1.20 ±0.18 | -1.20 ±0.01 | 1.21 ±0.15 | -1.22 ±0.21 |
| *Timp1* | -2.20 ±0.04 | 1.13 ±0.27 | -1.32 ±0.29 | -1.96 ±0.17 | -1.28 ±0.12 | -1.22 ±0.11 | -2.33 ±0.19 |
| *Timp2* | 2.21 ±0.04 | 1.46 ±0.22 | 1.14 ±0.01 | 1.13 ±0.09 | 1.67 ±0.13 | 1.28 ±0.12 | -1.69 ±0.15 |
| *Timp3* | ND | 2.70 ±0.43 | 1.40 ±0.23 | 1.98 ±0.46 | 1.09 ±0.16 | 1.04 ±0.29 | 1.11 ±0.17 |
| *Vash1* | ND | ND | ND | ND | -3.01 ±0.14 | 1.56 ±0.30 | 1.10 ±0.08 |
| *Vegfa* | 2.90 ±0.02 | 1.40 ±0.18 | 1.20 ±0.12 | 1.72 ±0.50 | 4.48 ±0.43 | 9.12 ±1.48 | 4.71 ±0.69 |
| *Vegfb* | 1.38 ±0.04 | 1.33 ±0.10 | -1.2 ±0.11 | 1.02 ±0.04 | -1.09 ±0.07 | 1.04 ±0.22 | -1.59 ±0.10 |
| *Vegfc* | ND | ND | 1.38 ±0.18 | ND | -5.03 ±0.05 | -11.37 ±0.03 | -2.86 ±0.16 |
| *Vegfd* | ND | -1.19 ±0.17 | -1.73 ±0.22 | -1.99 ±0.23 | 1.05 ±0.37 | 1.48 ±0.29 | 1.08 ±0.24 |
| *Vegfr2 (Kdr)* | -2.05 ±0.05 | 1.25 ±0.13 | 1.07 ±0.04 | 1.43 ±0.21 | 1.31 ±0.10 | 2.29 ±0.38 | -1.01 ±0.30 |

**EV Table 2.**
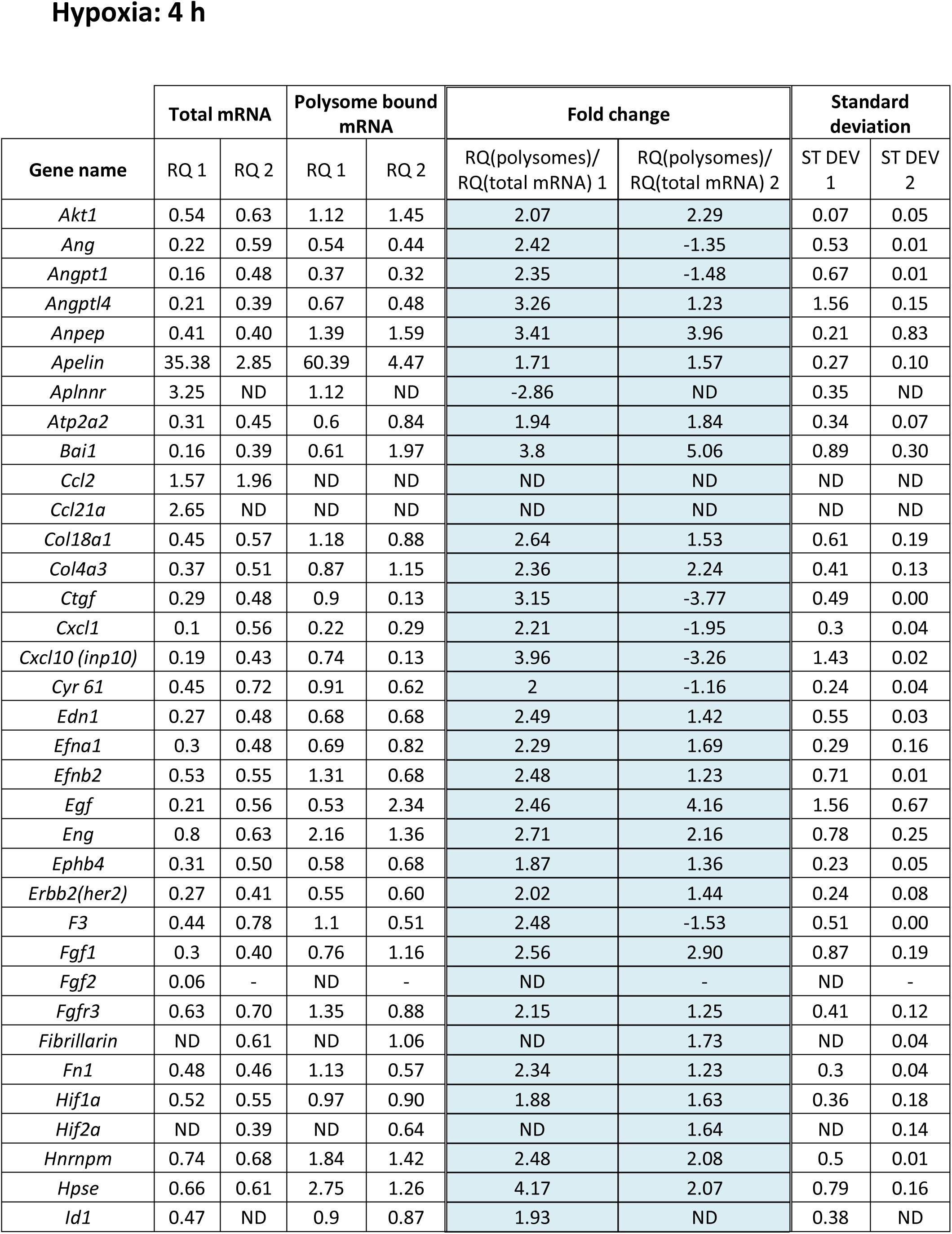
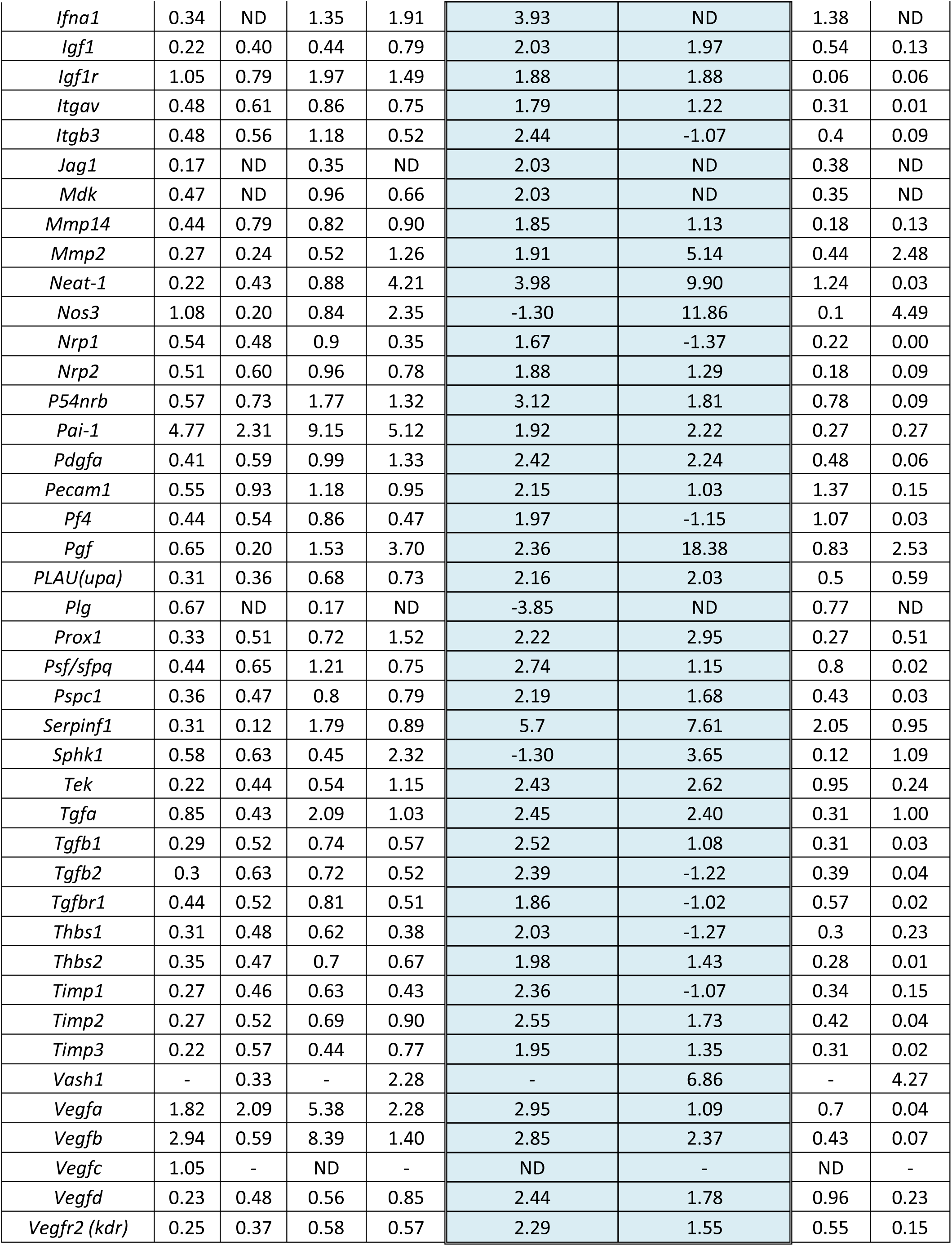
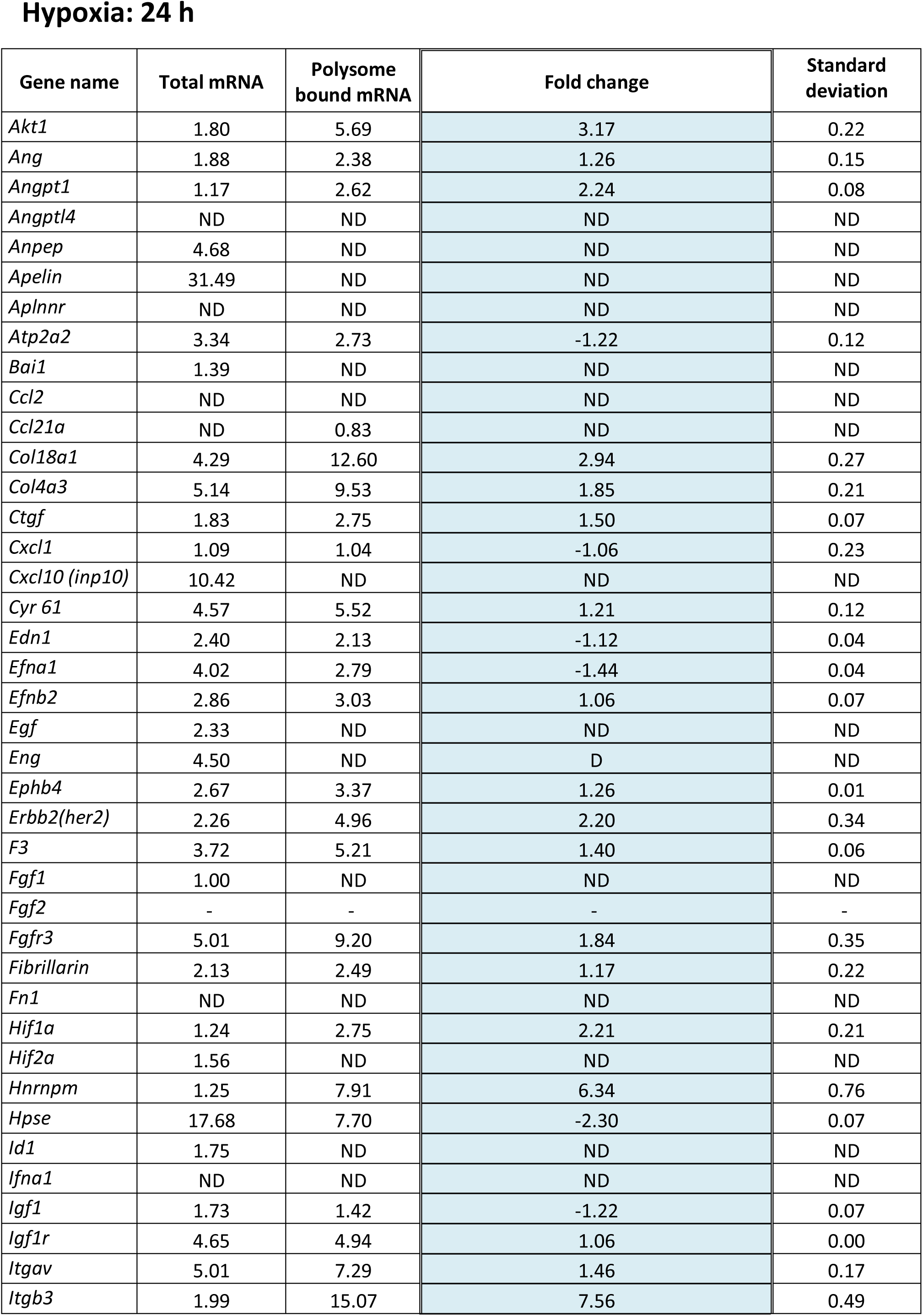
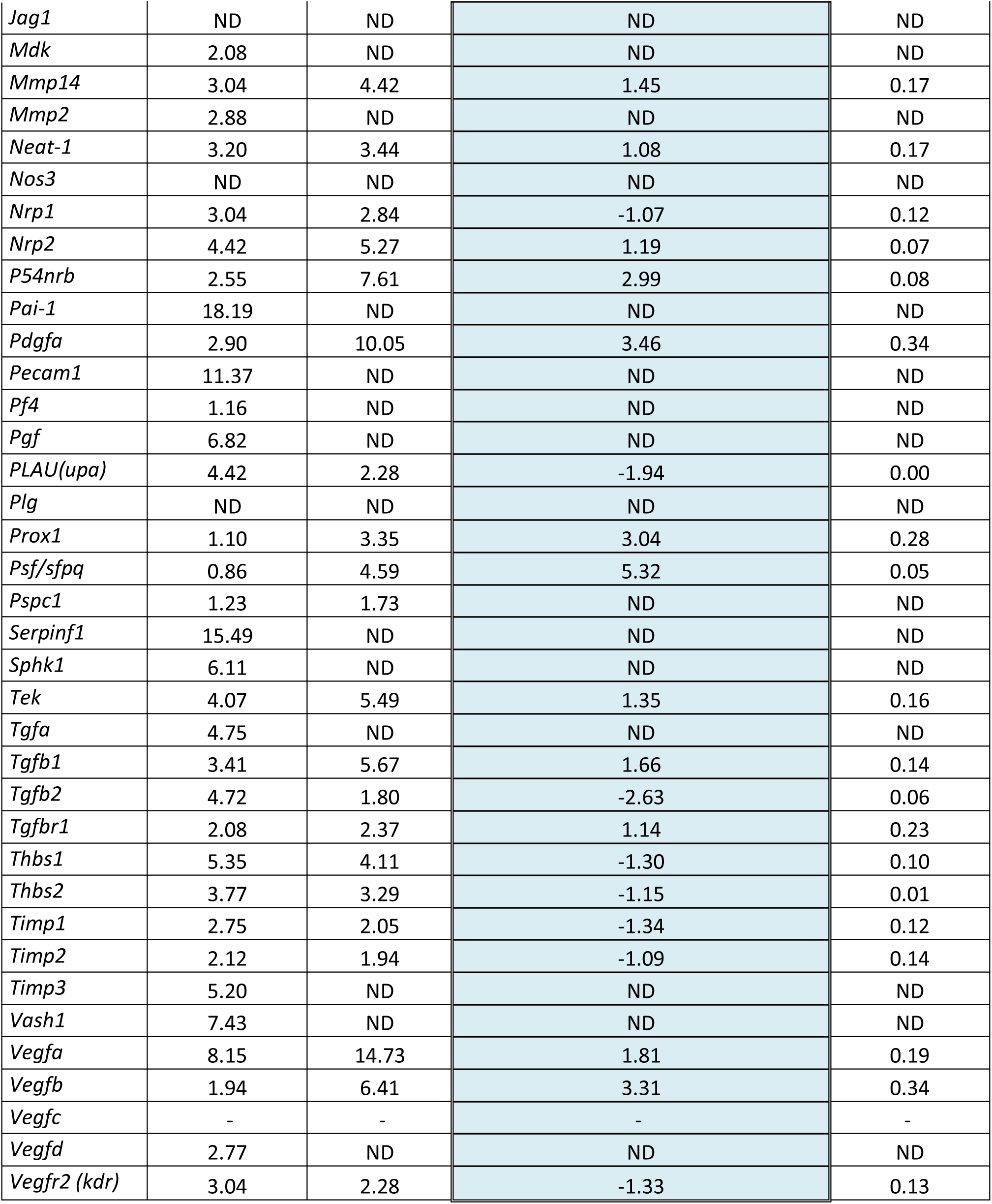
Translatome of (lymph)angiogenic factor genes in hypoxic HL-1 cardiomyocytes. Polysomes were purified on sucrose gradient from HL-1 cardiomyocytes either in normoxia or after 4 h or after 24 h of hypoxia at 1% O2, as described in Materials and Methods. RNA was purified from polysome-bound and from cell lysate (before gradient loading). cDNA and PCR array was performed as in Figure 1 and in EV Table 1. Relative quantification (RQ) of gene expression in hypoxia was calculated using the 2^-ΔΔCT^ method (polysomal RNA/total RNA normalized to normoxia). The 4 h time of hypoxia array was repeated in two independent arrays (RQ1 and RQ2). The values presented in Figures 2 and 3 correspond to RQ1 values. For each array, gene expression analysis was performed in three replicates. Standard deviation is indicated. When the RQ value is inferior to 1, the fold change is expressed as −1/RQ. ND means “non detected”. “-“ means that the gene was not included in the array.

**EV Table 3.**
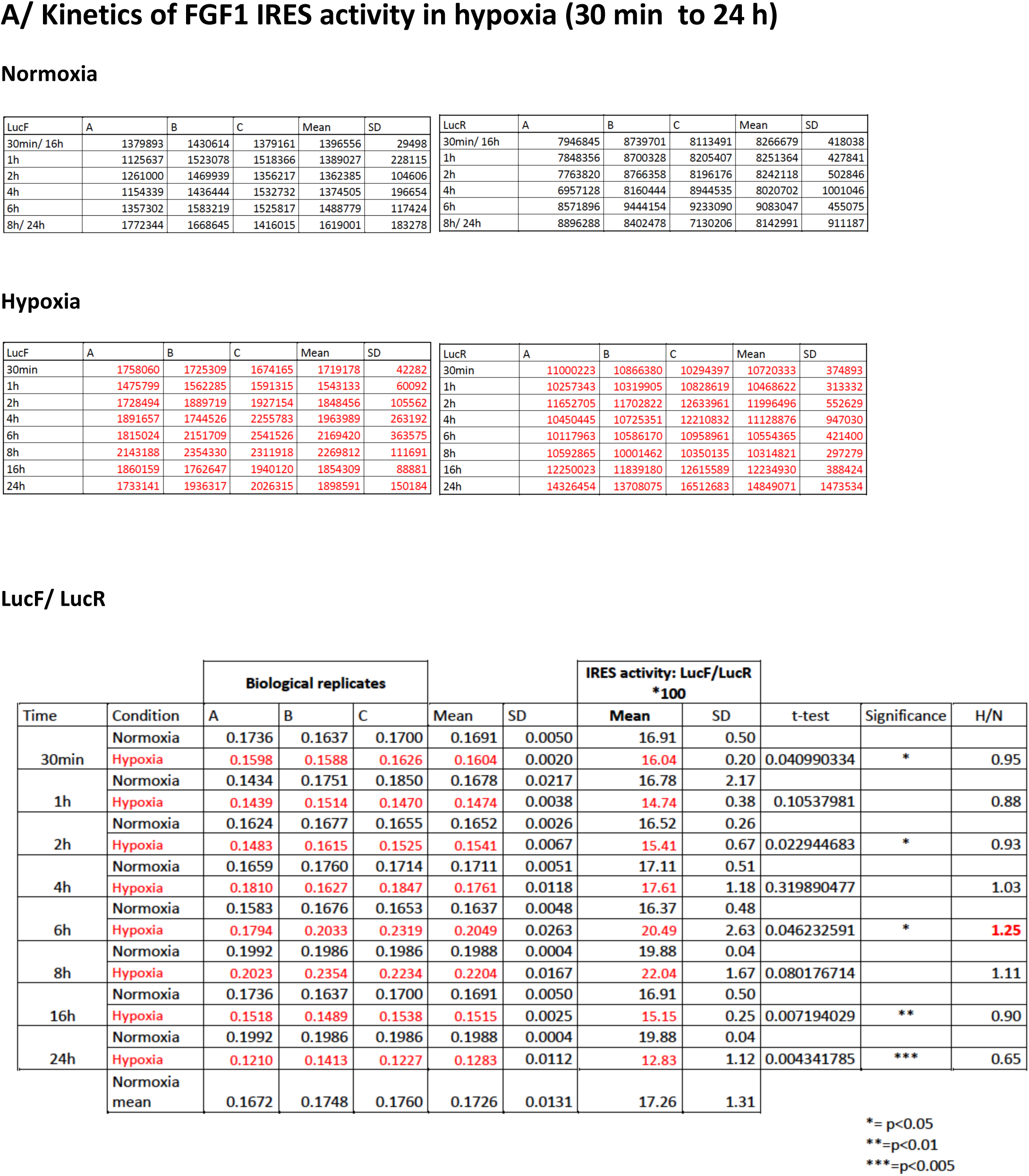
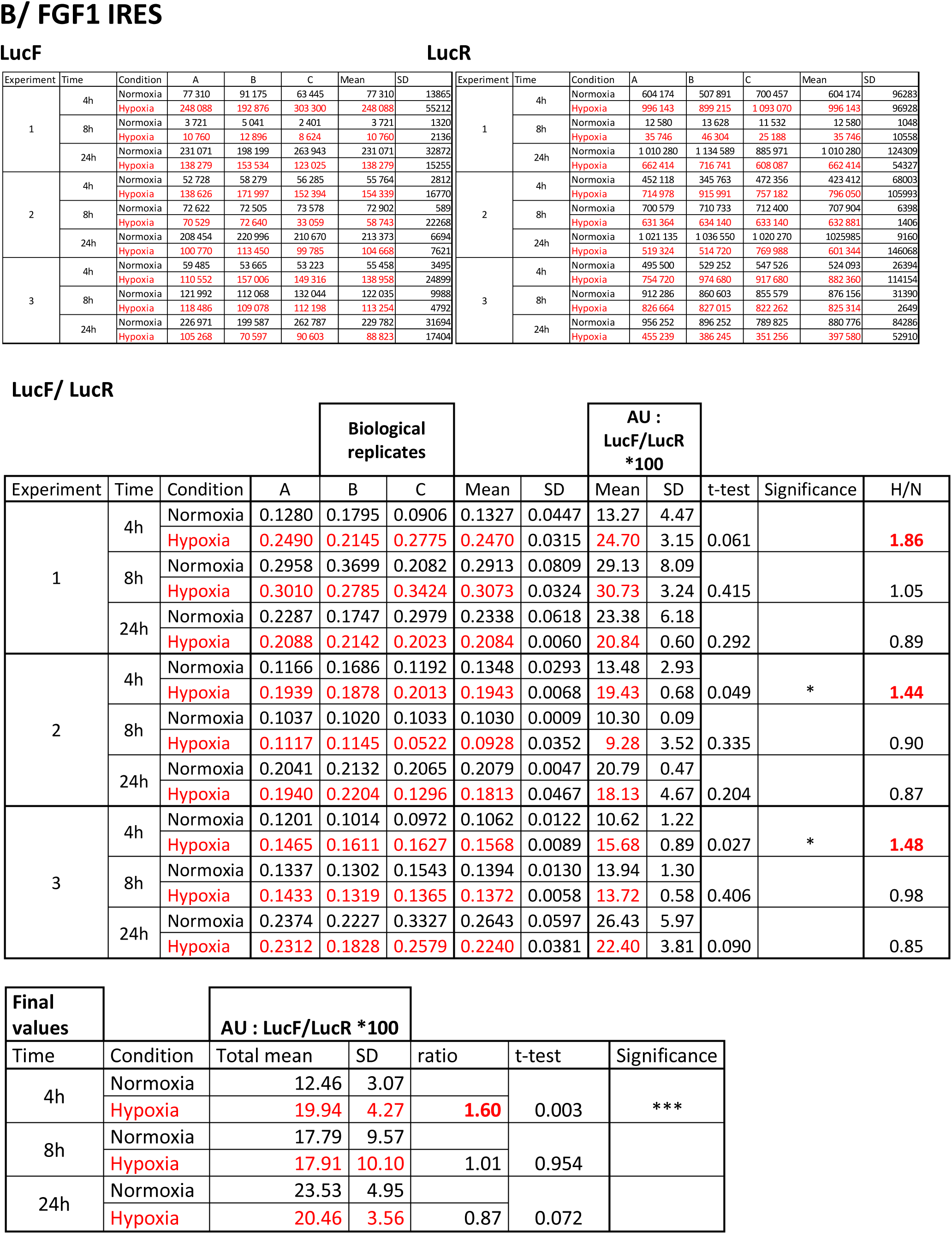
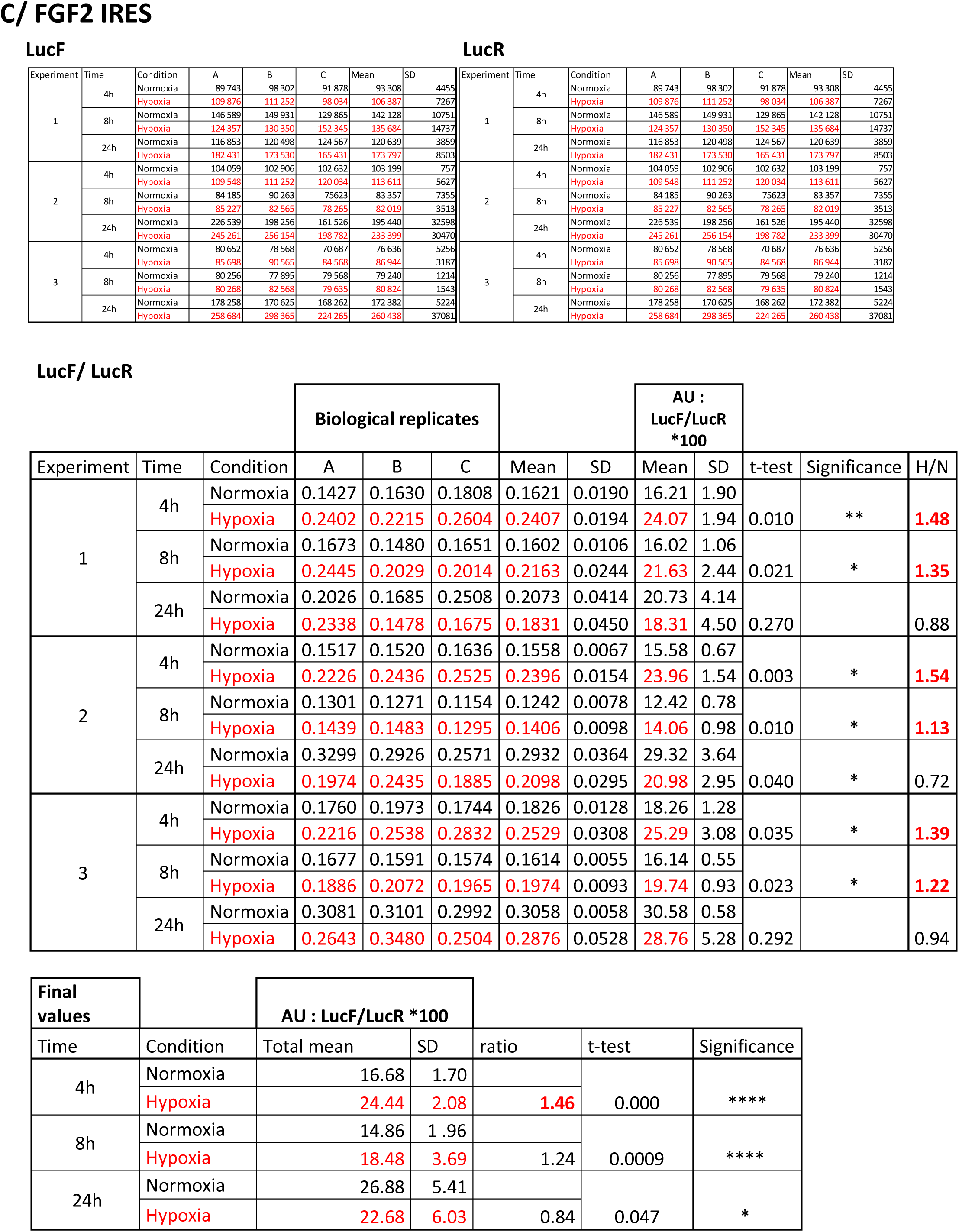
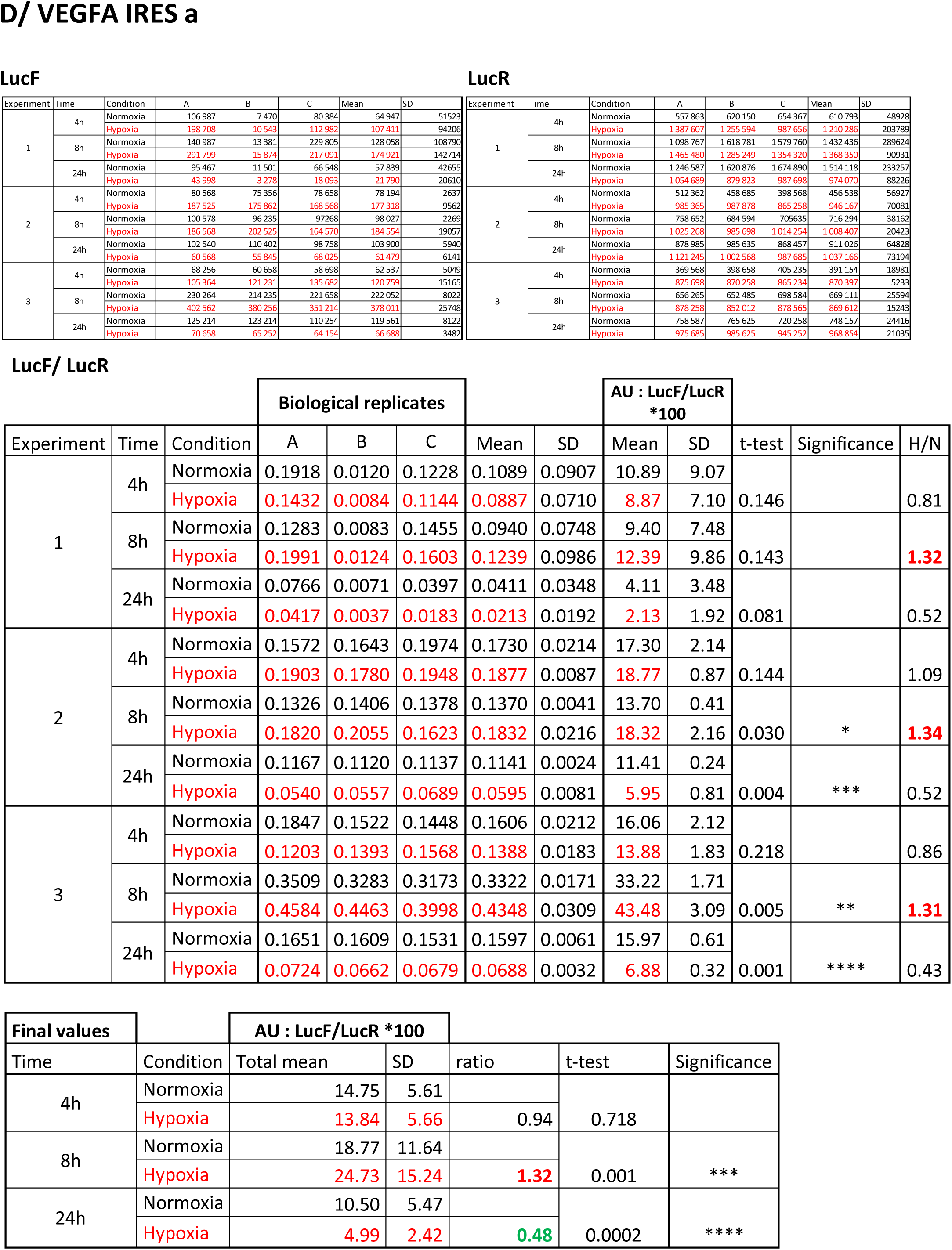
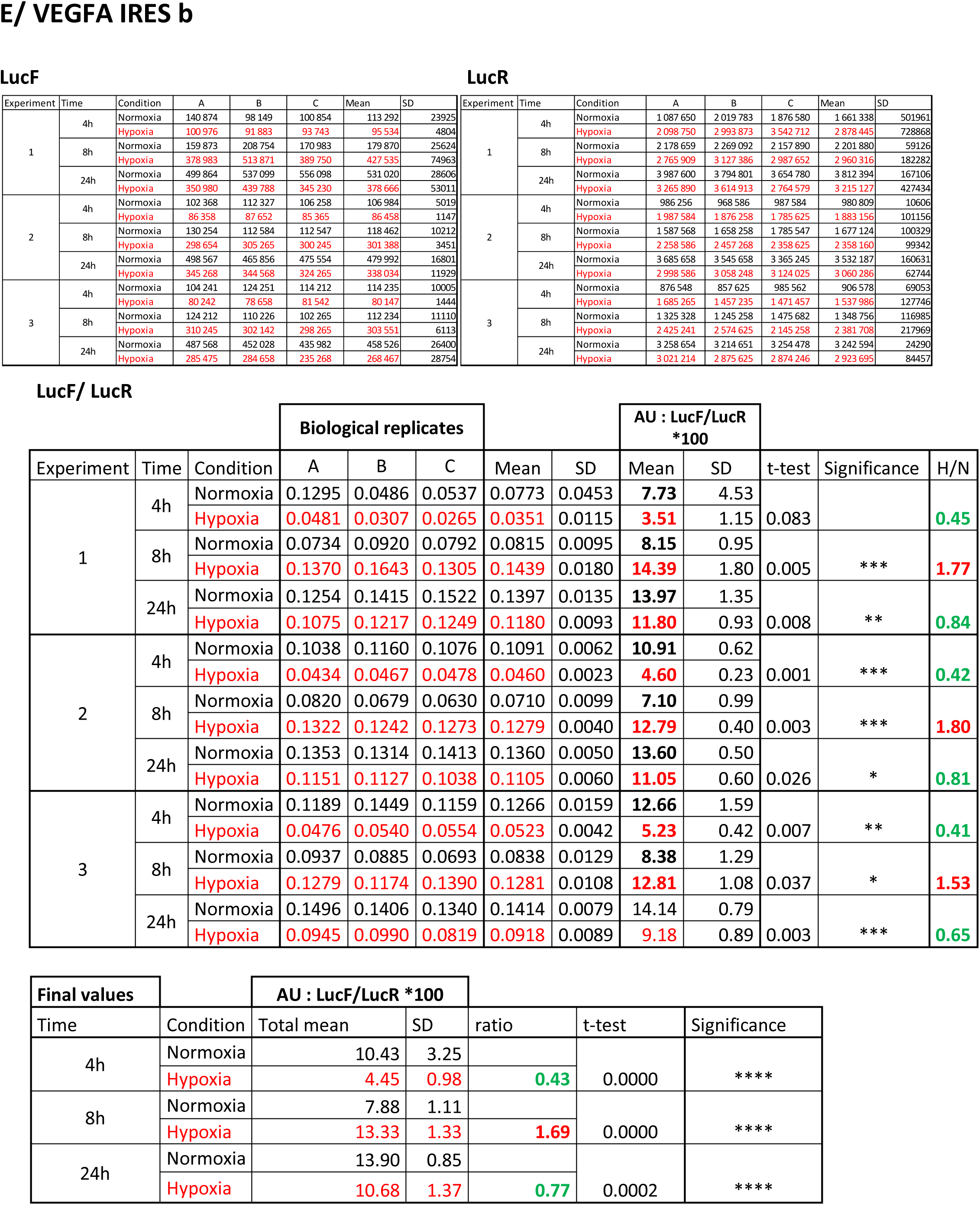
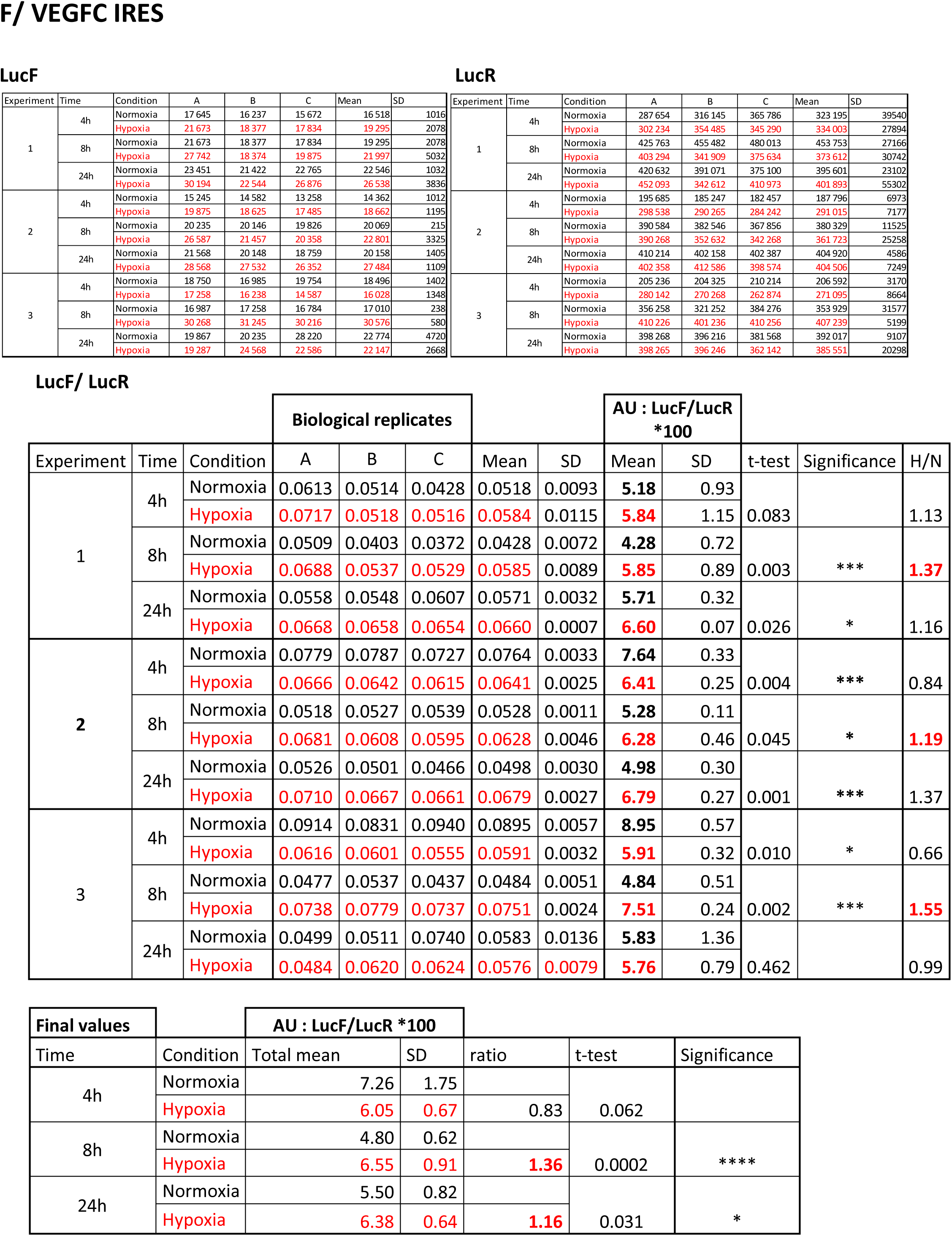
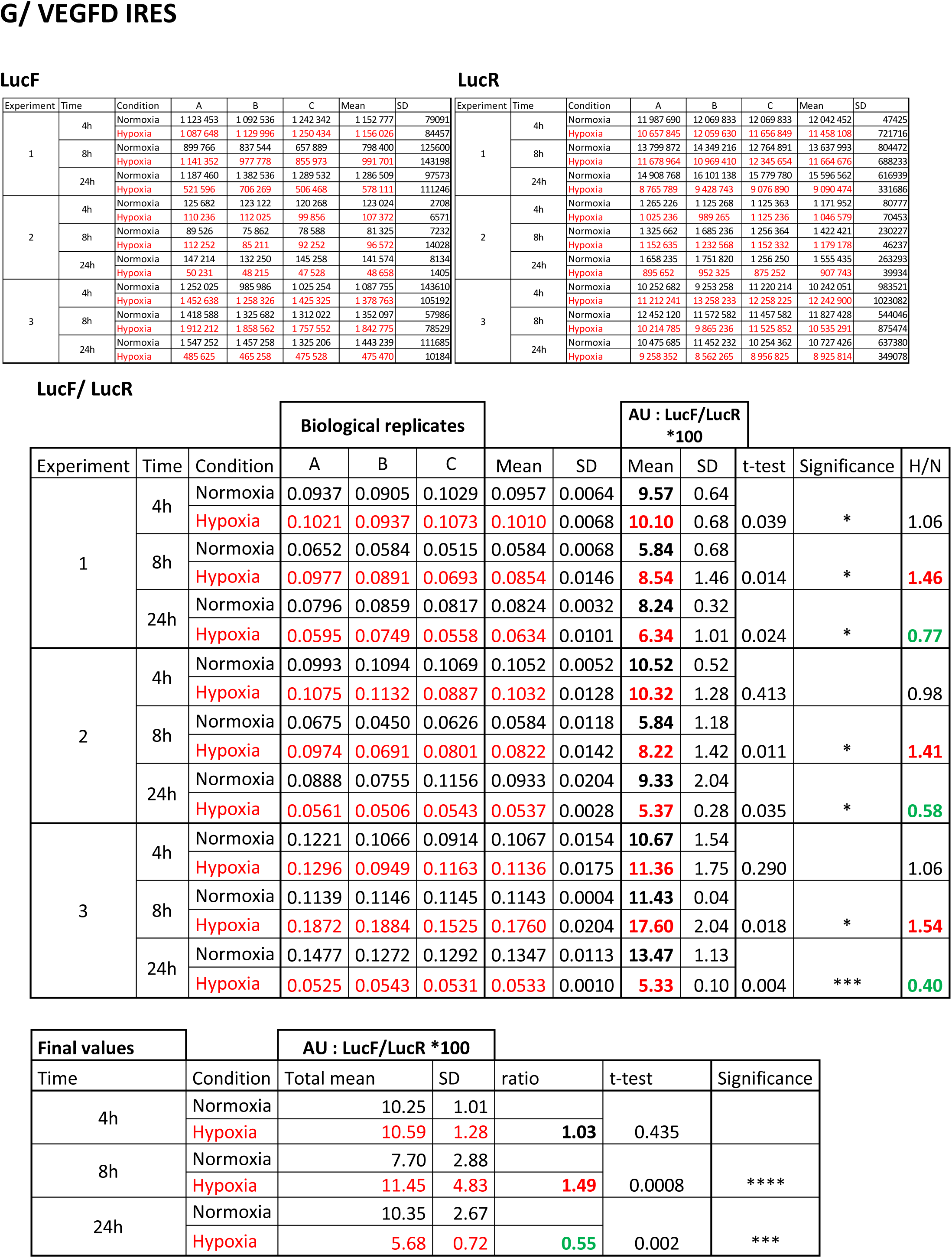
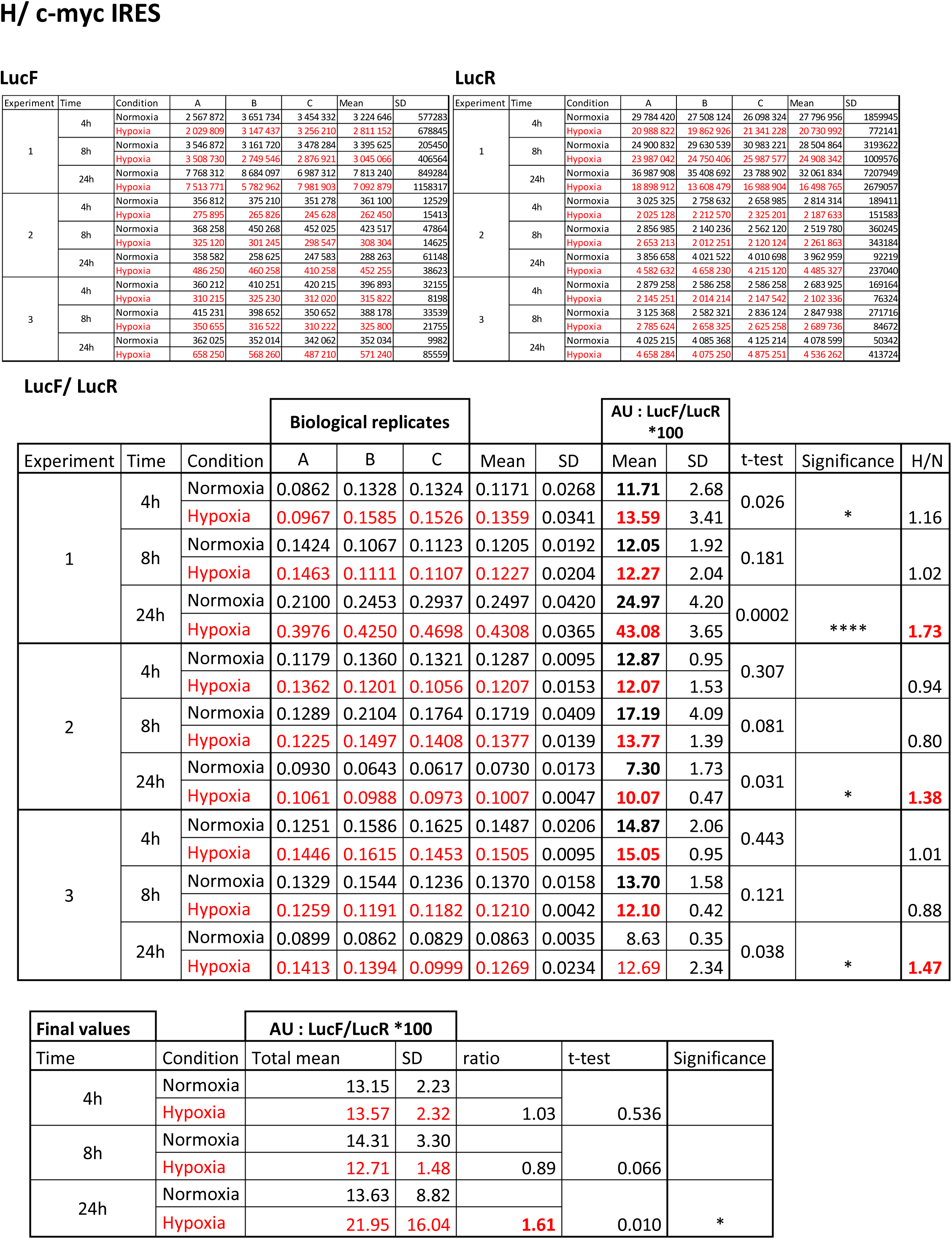
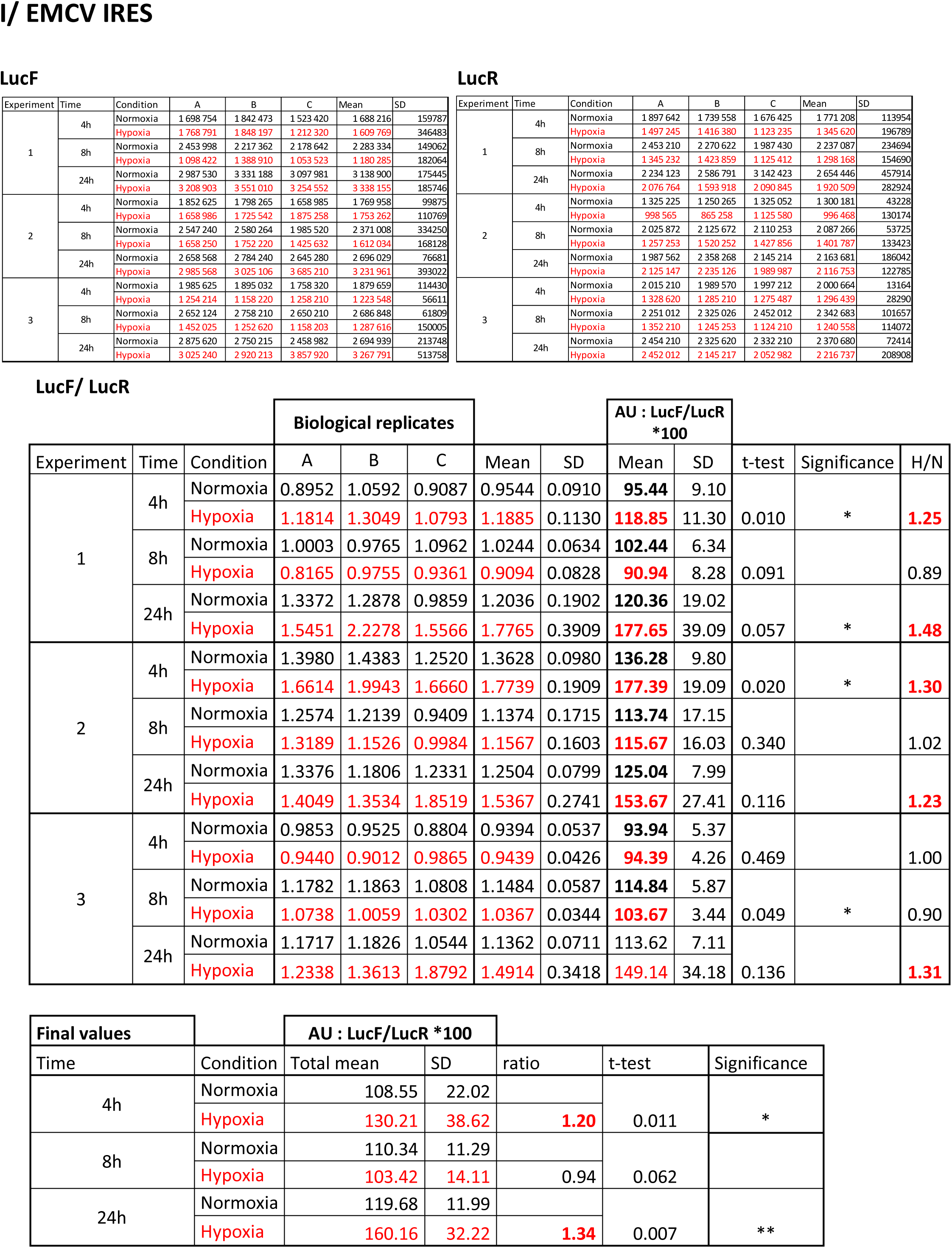
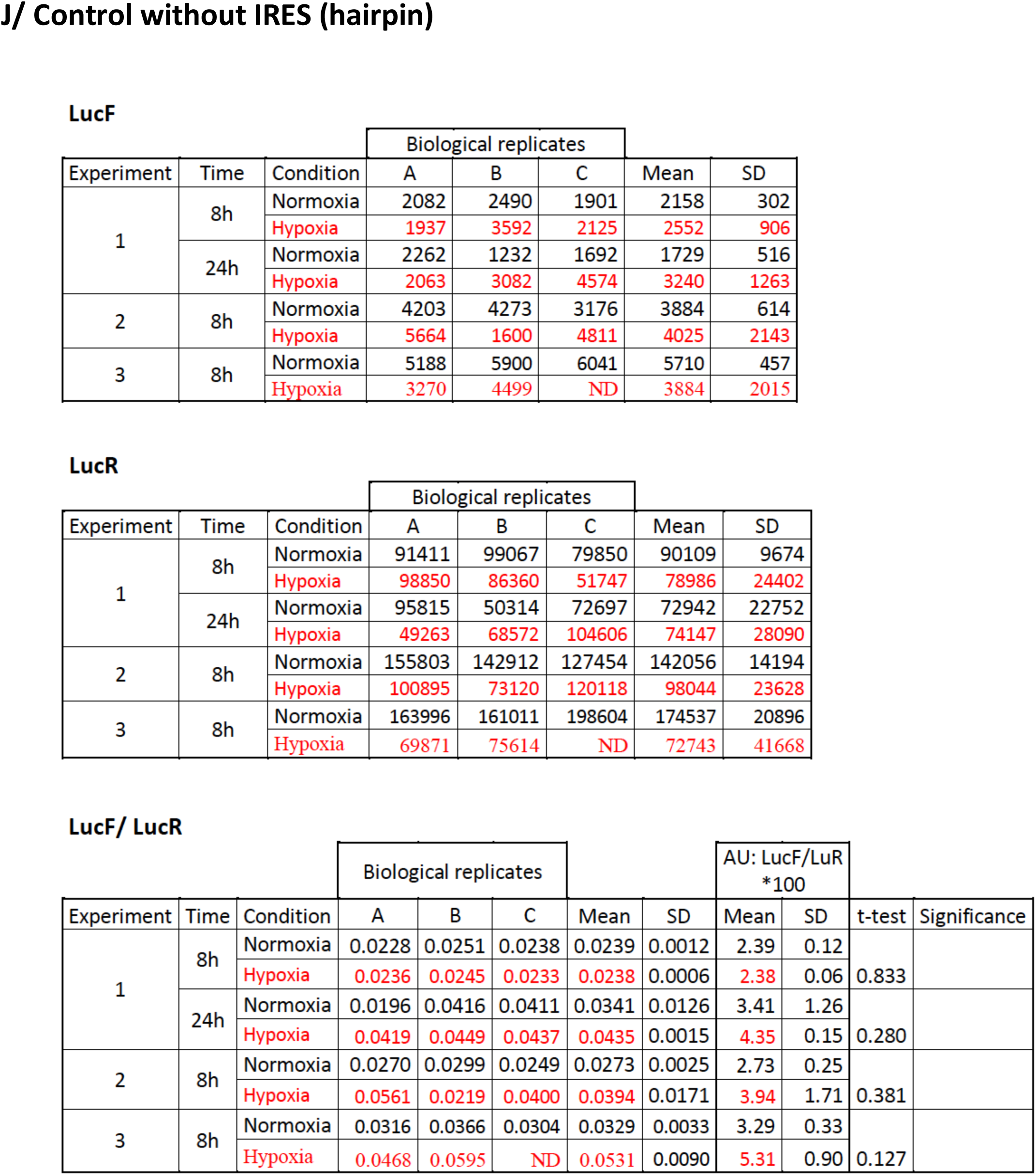
IRES activities at different times of hypoxia in HL-1 cells. Luciferase activity values and IRES activities corresponding to the experiments presented Figure 4. A/ Kinetics of FGF1 IRES activity from 30 min to 24 h B-I/ Activities of the different IRES at 4 h, 8 h and 24 h of hypoxia J/ Negative control with a lentivector containing a hairpin (no IRES) between the two luciferase cistrons. Biological replicates are indicated as A, B and C, whereas independent experiments are indicated as 1, 2, 3. Means, standard deviation (SD) and t-test of IRES activities were calculated. The panels “final values” correspond to means of all experiments (nine values) which are reported in the histograms of Figure 4. P-value significance is indicated: *p<0.05, **p<0.01, ***<0.001, ****p<0.0001.

**EV Table 4.**
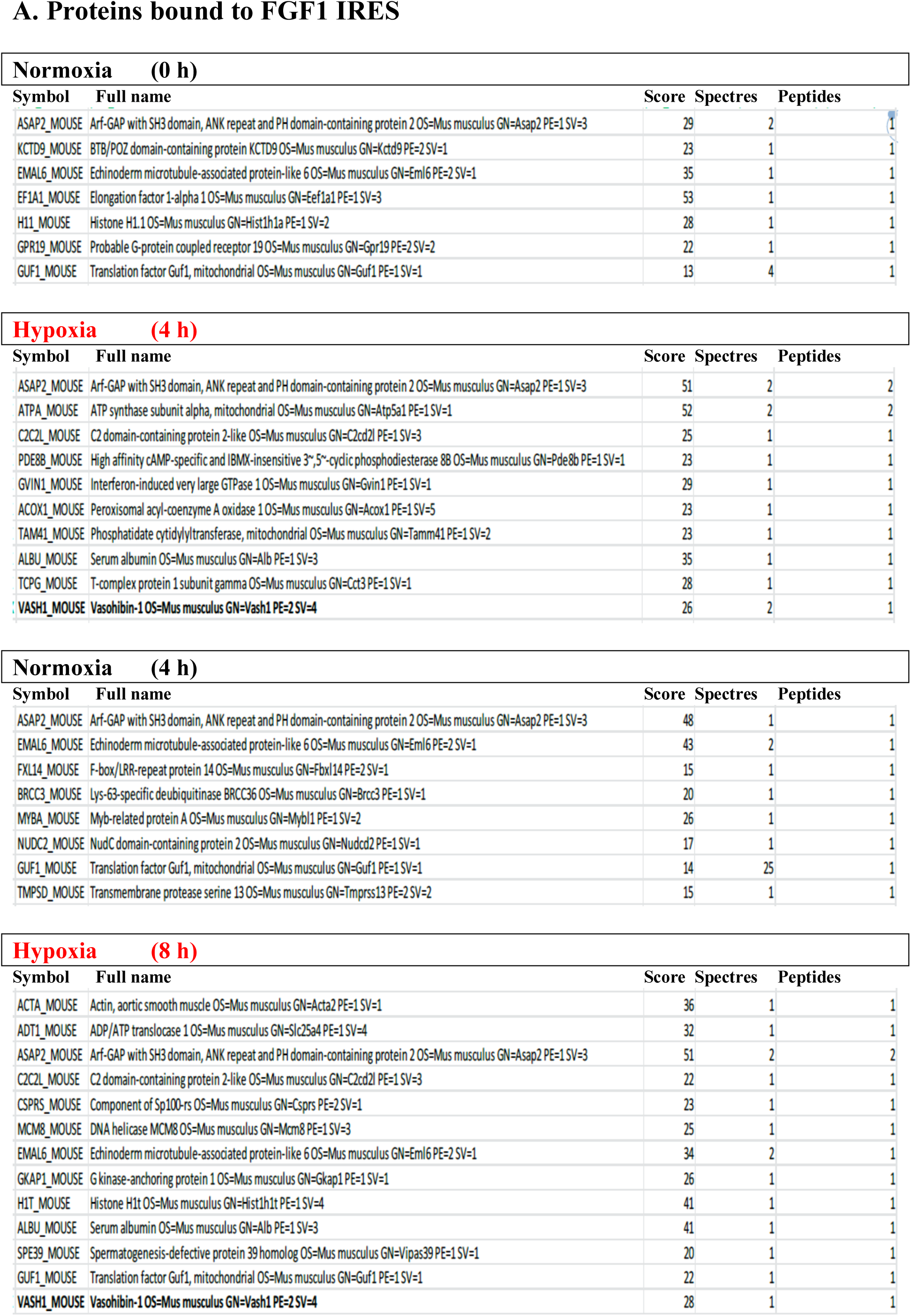
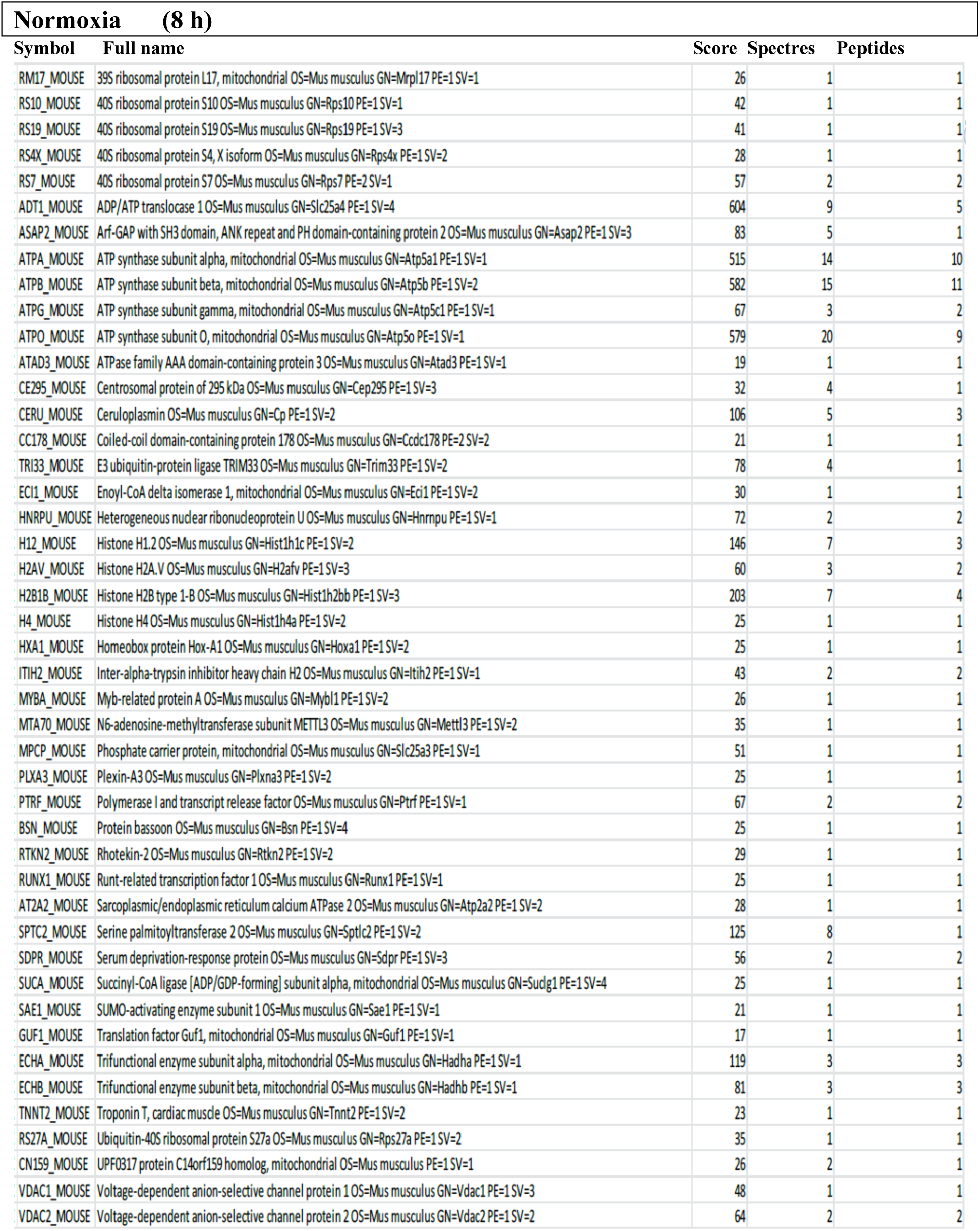
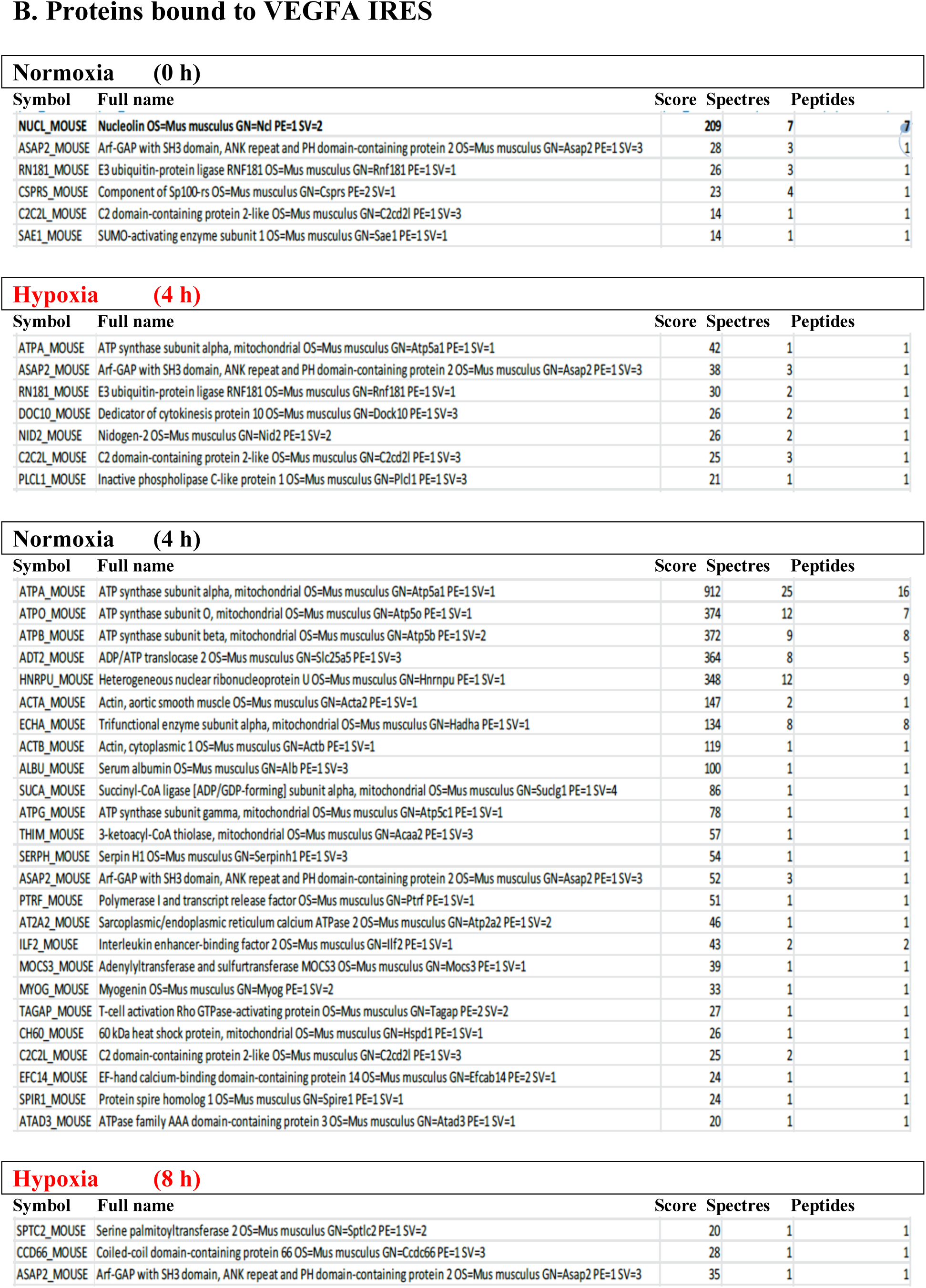
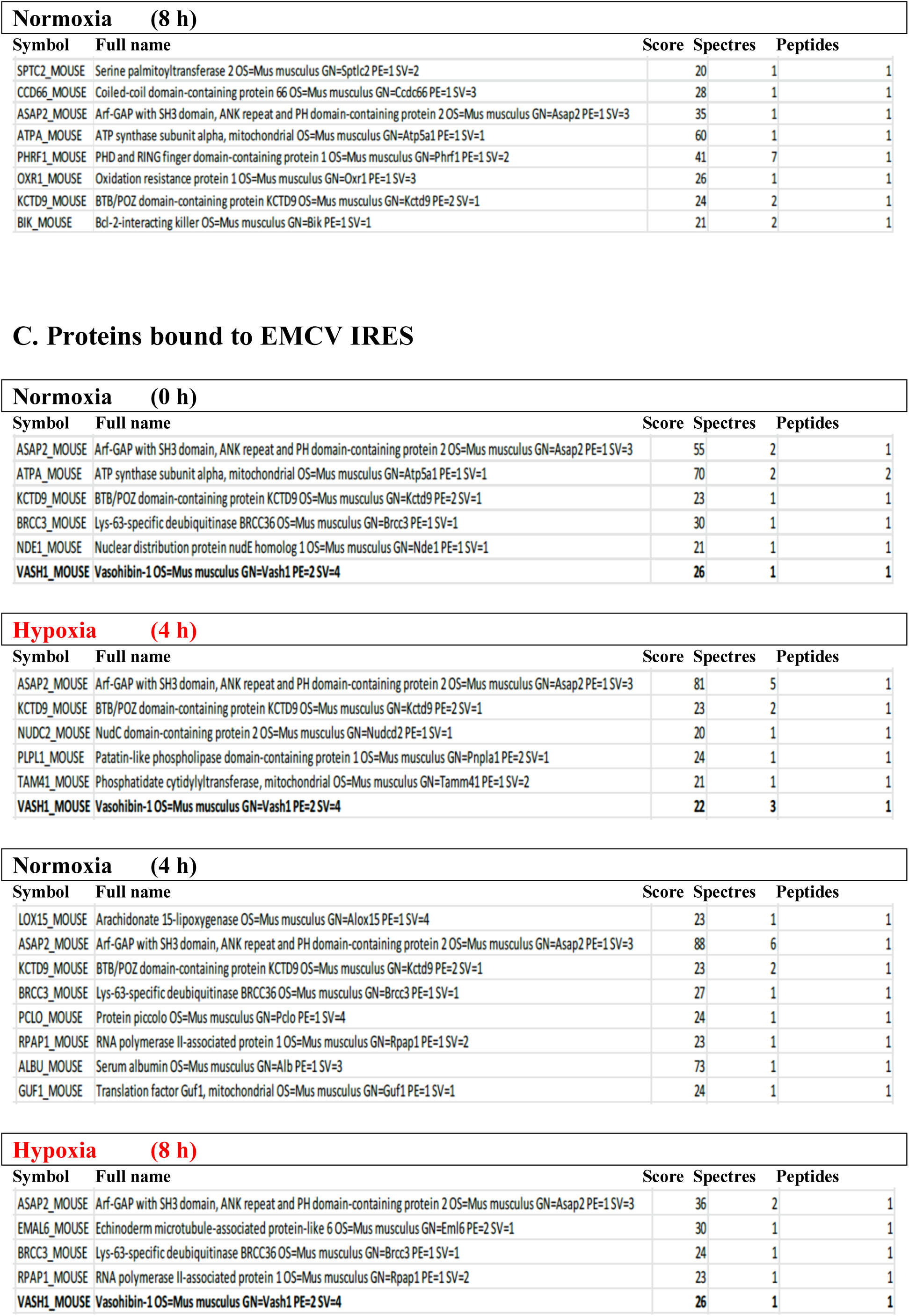
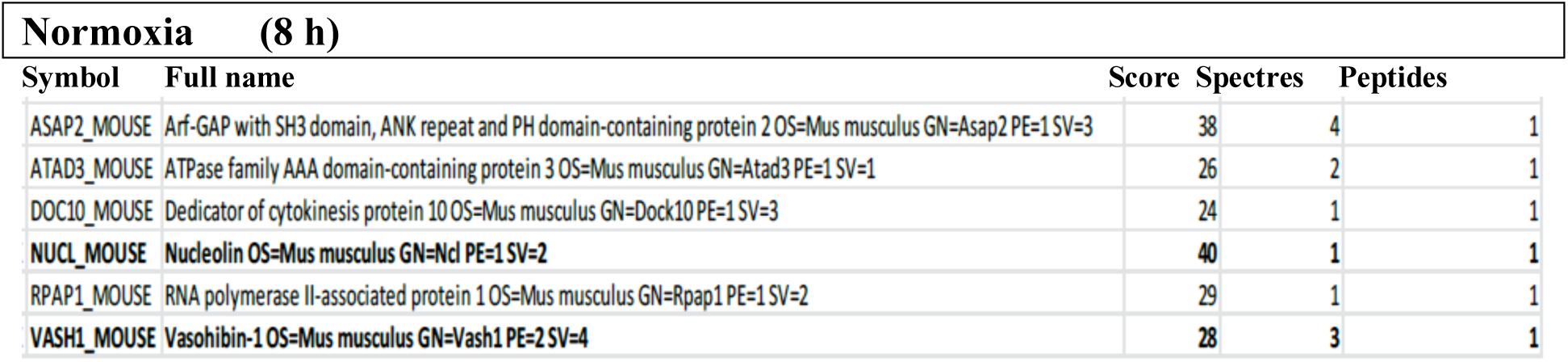
BIA-MS analysis of IRES-bound proteins in hypoxic cardiomyocytes. Total cell extracts from normoxic or hypoxic HL-1 cardiomyocytes were injected into the BIAcore T200 optical biosensor device where biotinylated IRES RNAs had been immobilized. The list of bound proteins identified by mass spectrometry (LC-MS/MS) after tryptic digestion is shown for FGF1 (A), VEGF-Aa (B) or EMCV (C) IRESs, respectively. The score and the number of spectra and peptides identified are indicated. For each time of hypoxia, cells were cultivated the same time in normoxia as a control (Normoxia 4h and 8h).

**EV Table 5.**
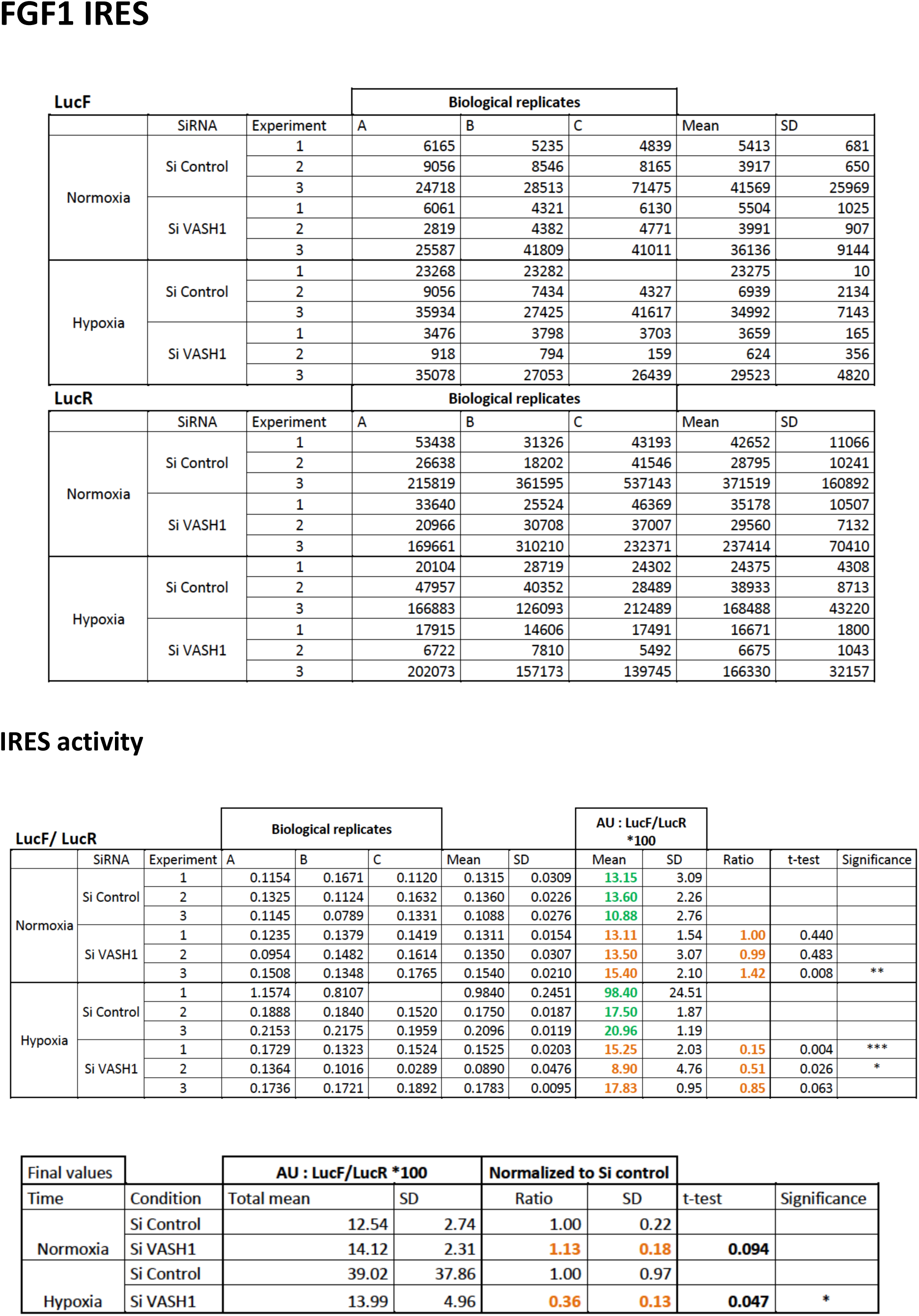
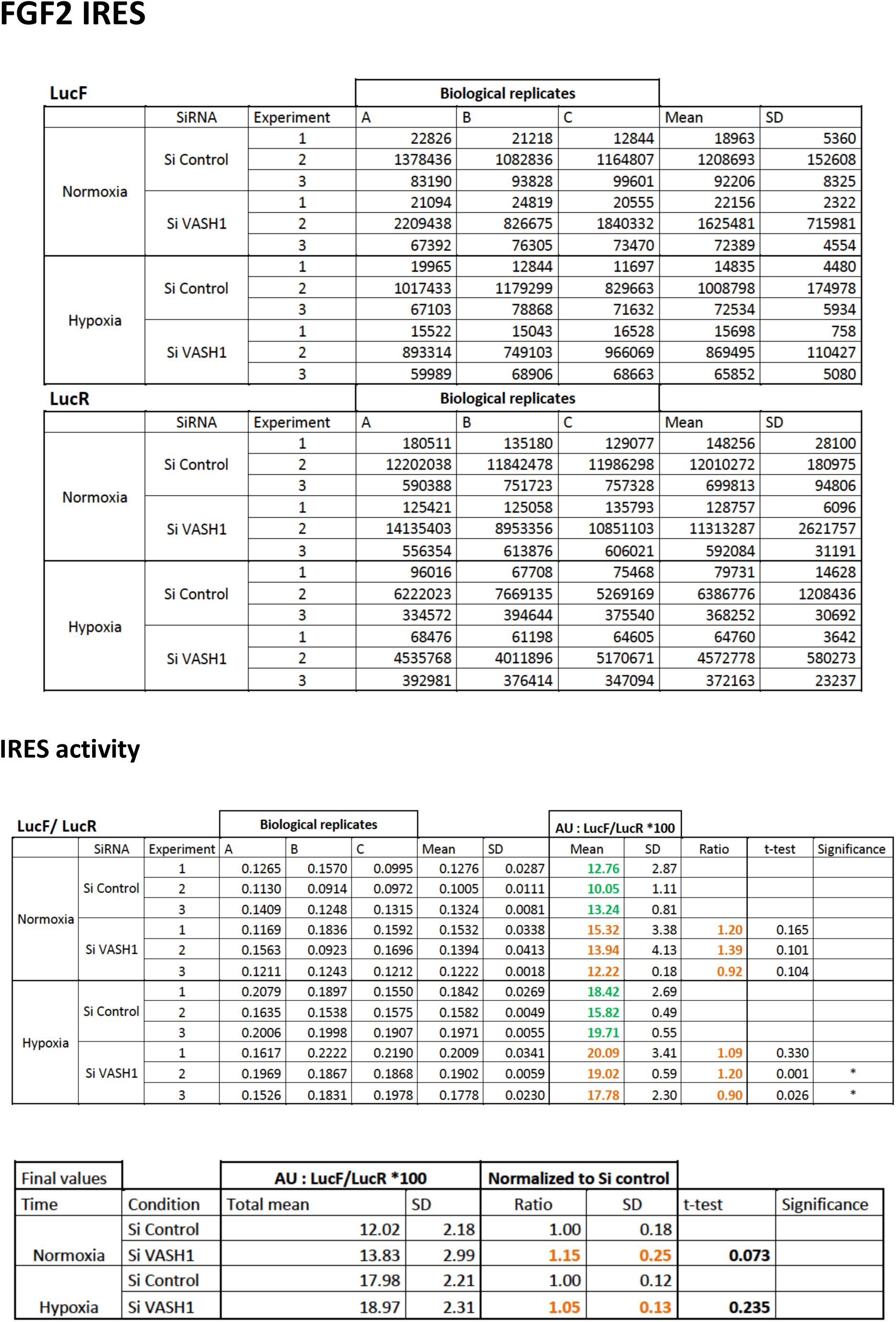
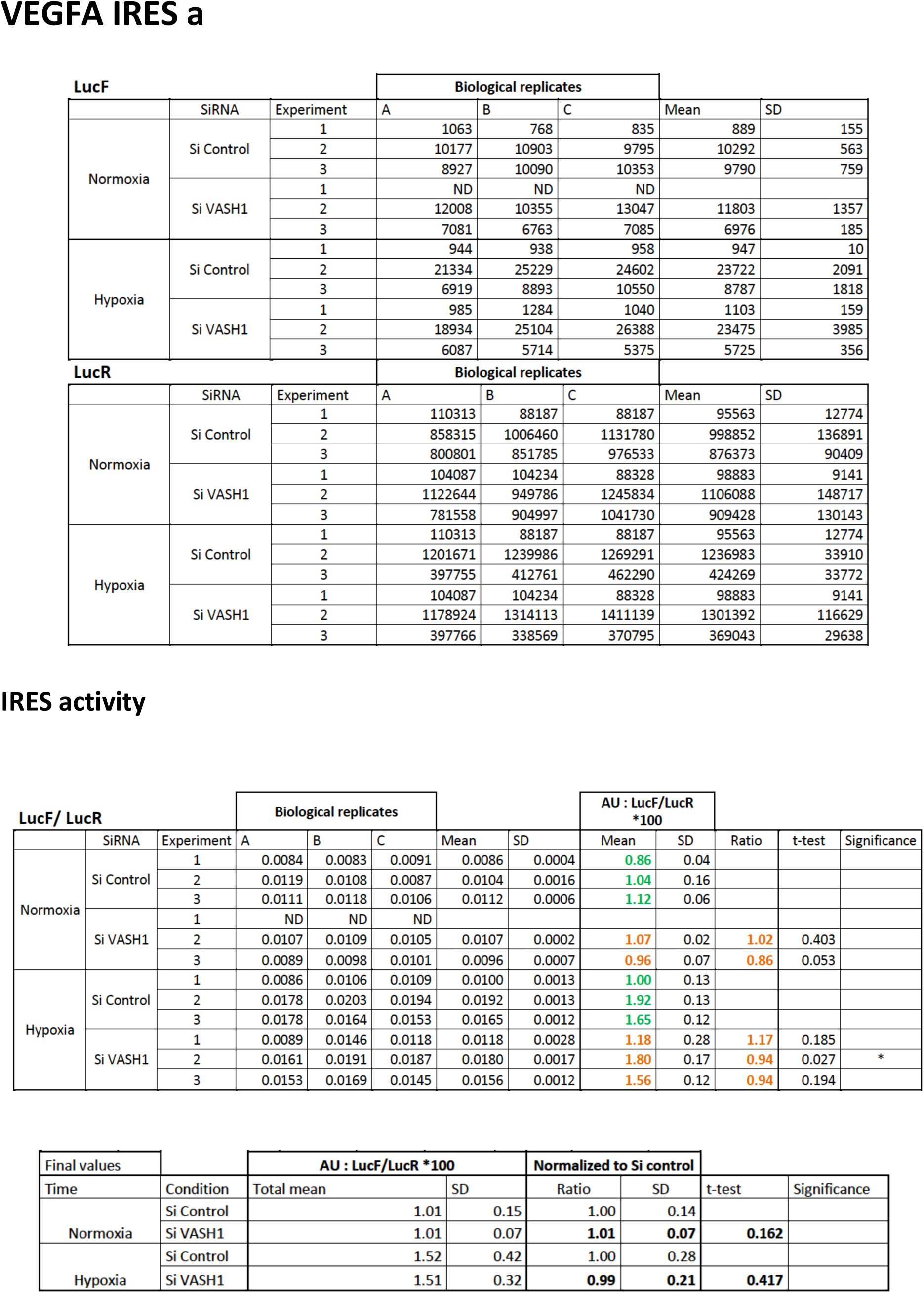
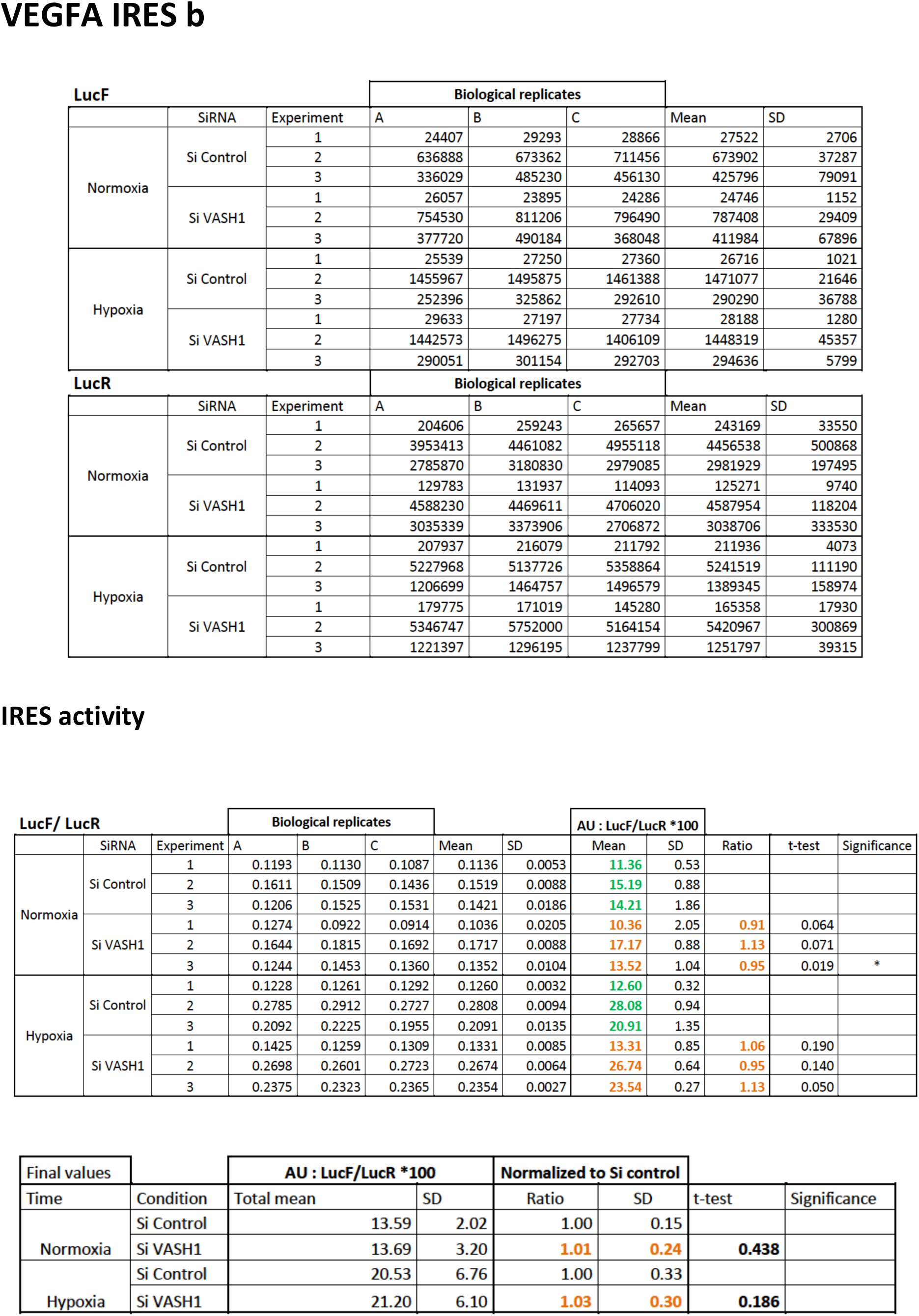
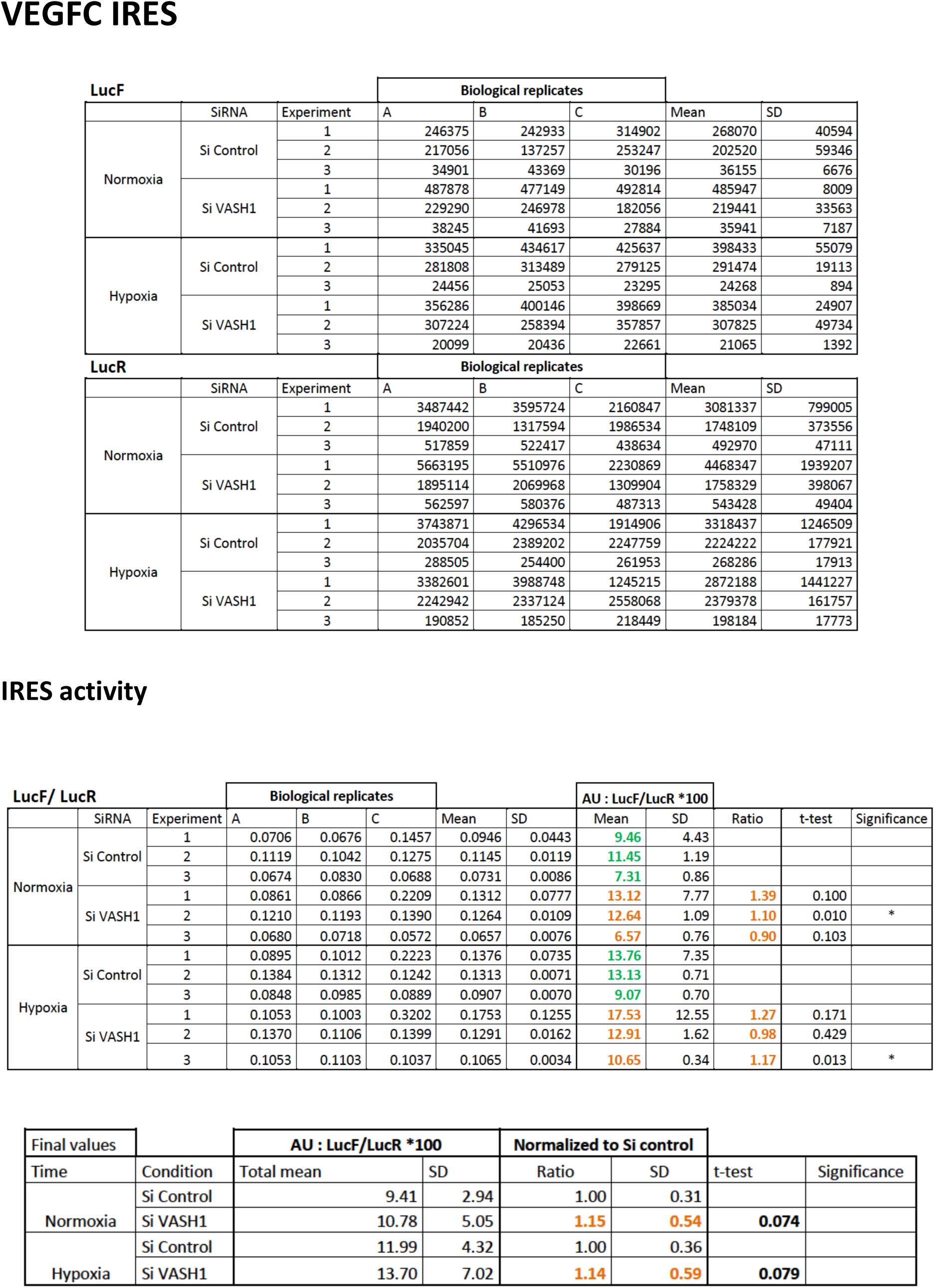
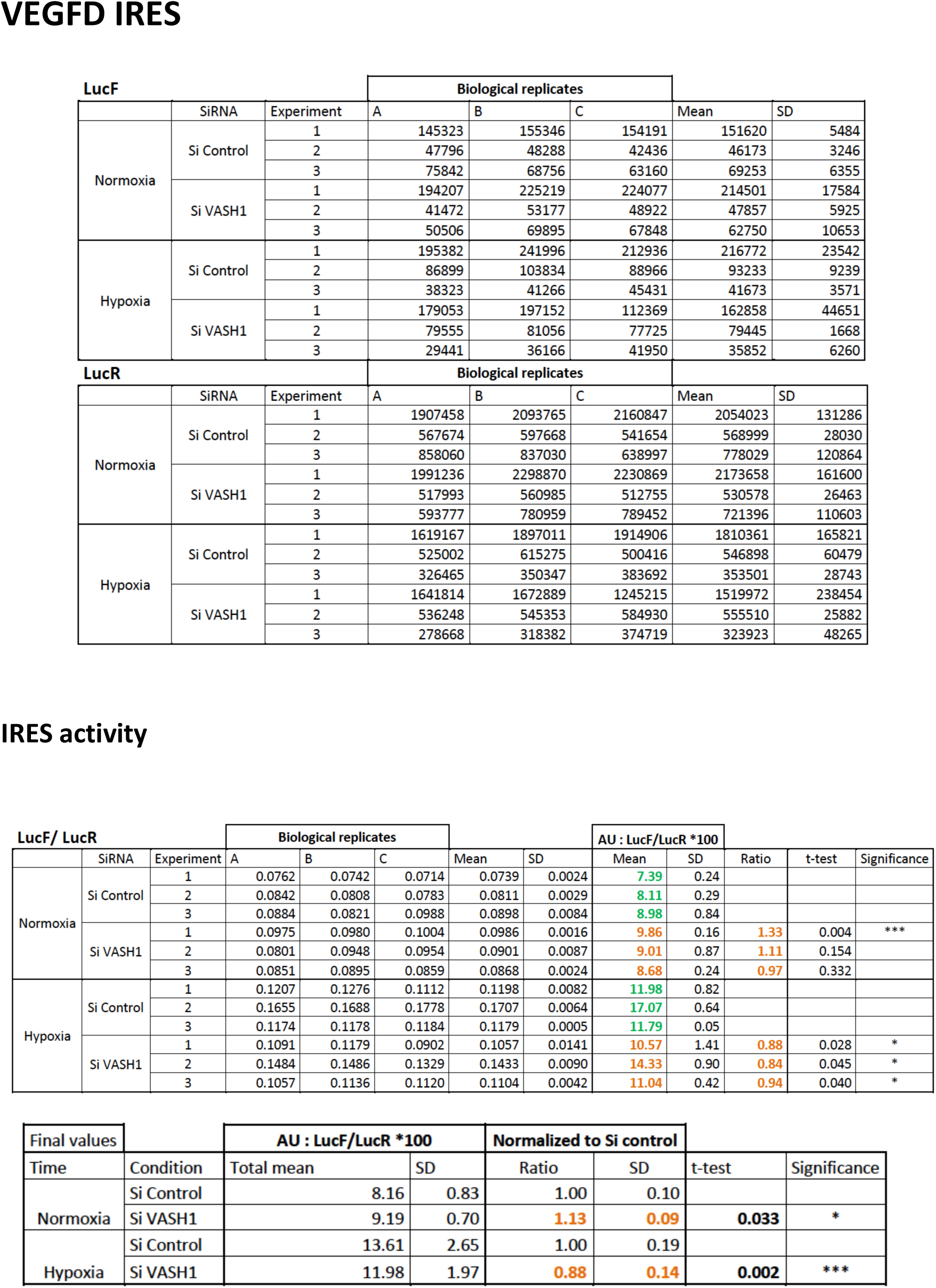
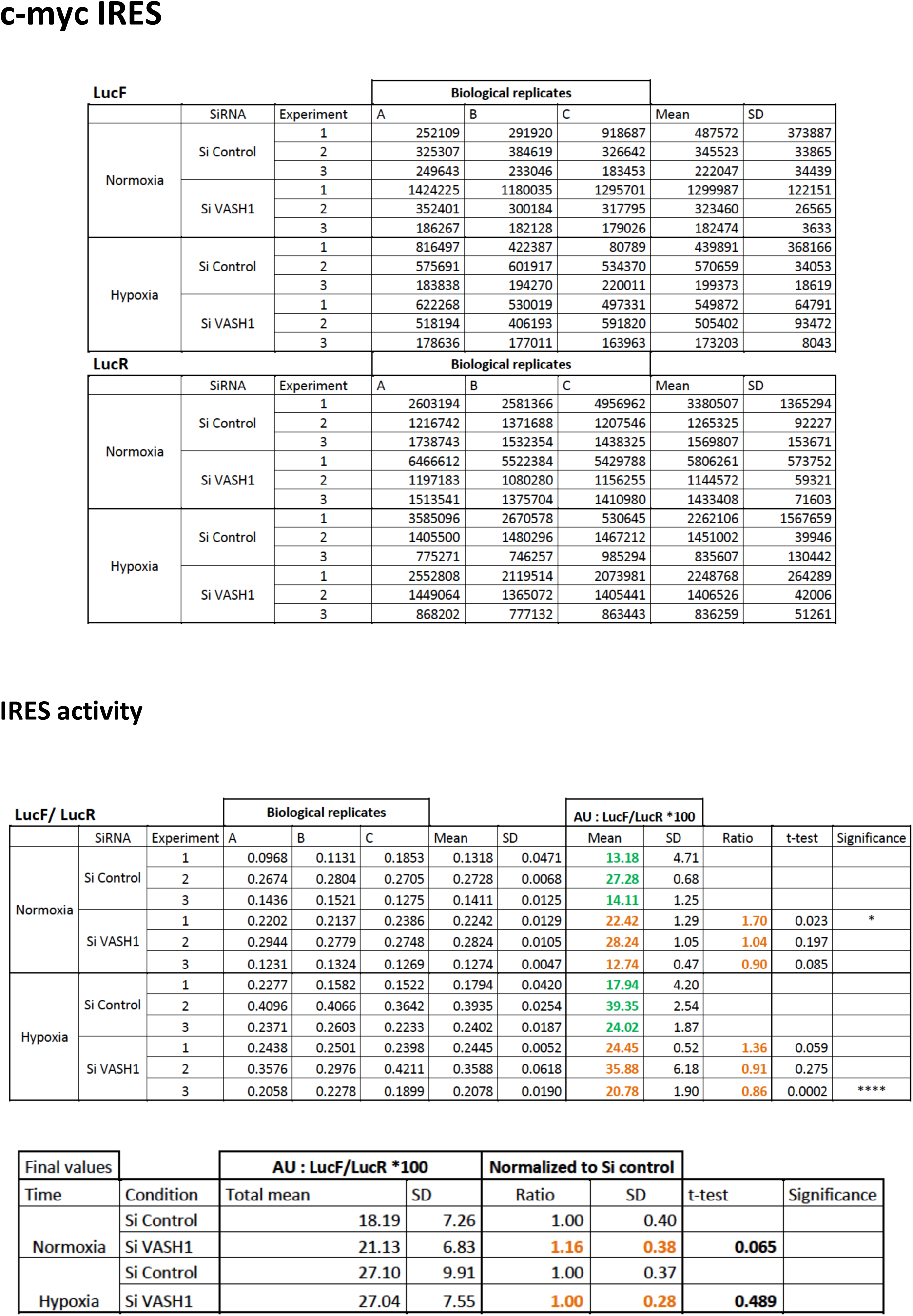
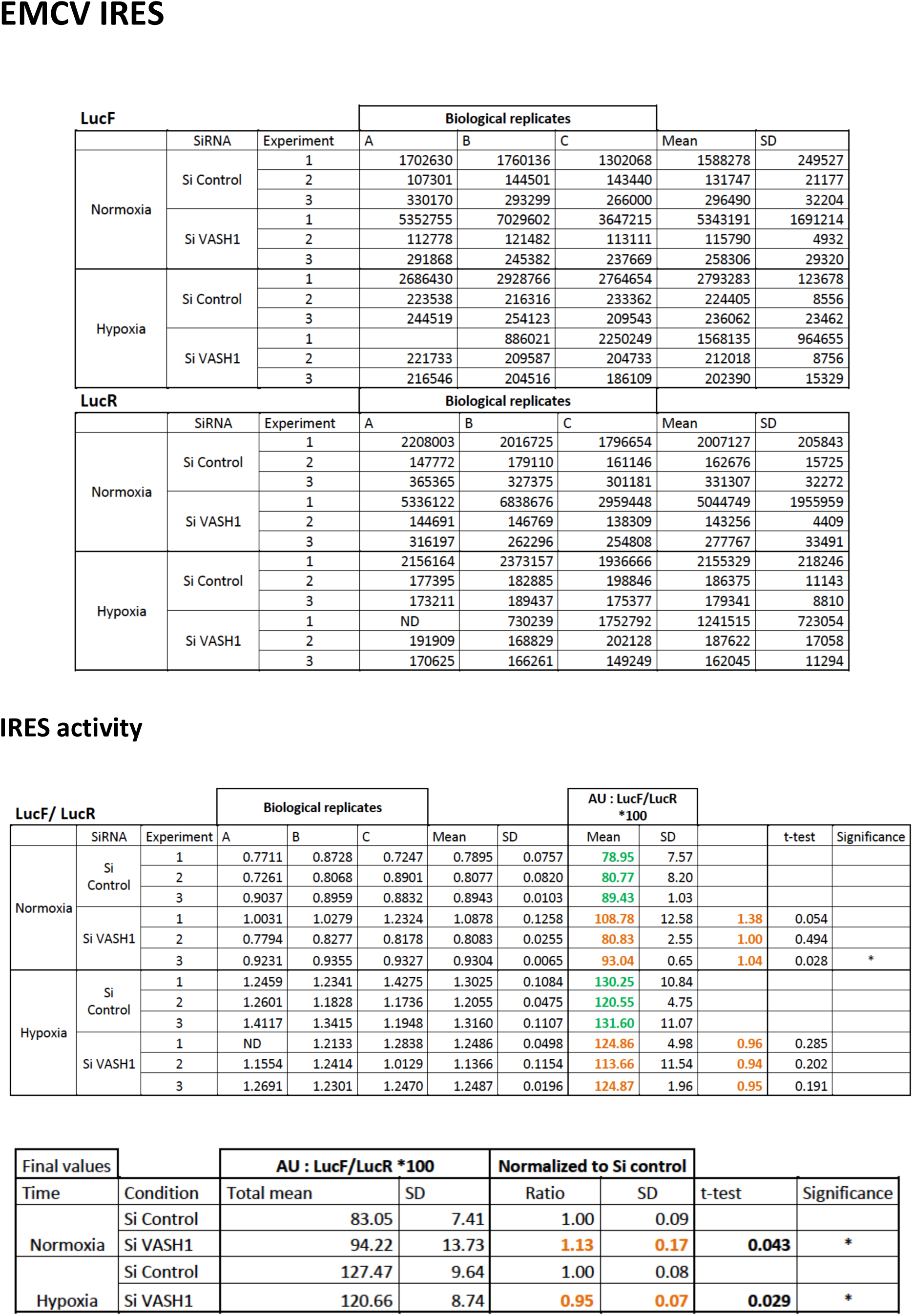
Knock-down of VASH1 in HL-1 cells. HL-1 cells transduced by the different IRES-containing lentivectors were transfected with siRNA siVASH of SiControl and submitted to 8 h of hypoxia. Luciferase activity and IRES activities (ratio LucF/LucR x 100) were measured. The values correspond to the experiments presented Figure 4. Biological replicates are indicated as A, B and C, whereas independent experiments are indicated as 1, 2, 3. Means, standard deviation (SD) and t-test of IRES activities were calculated. The panels “final values” correspond to means of all experiments (nine values) which are reported in the histograms of Figure 7. P-value significance is indicated: *p<0.05, **p<0.01, ***<0.001, ****p<0.0001.

**EV Table 6.**
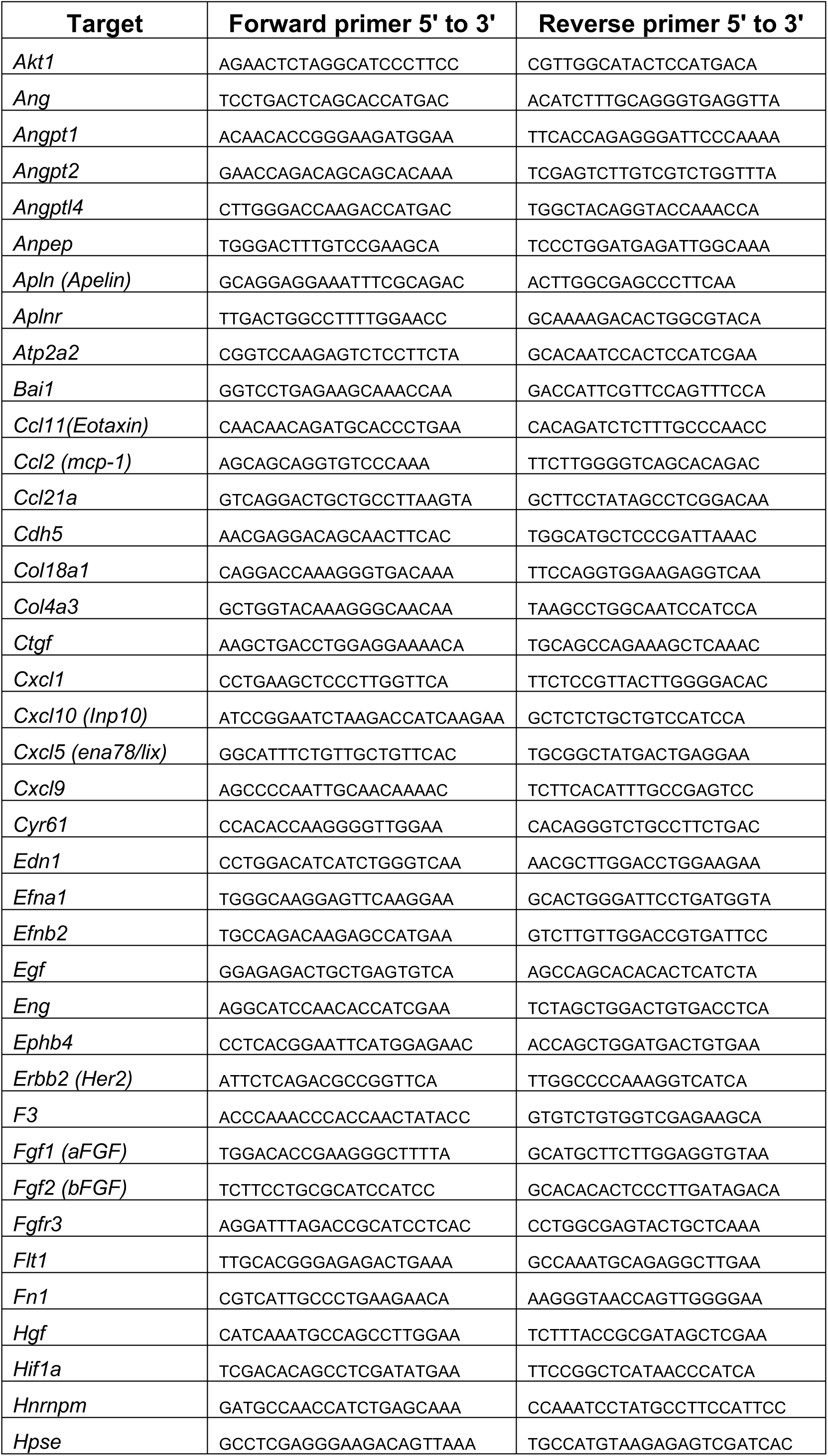
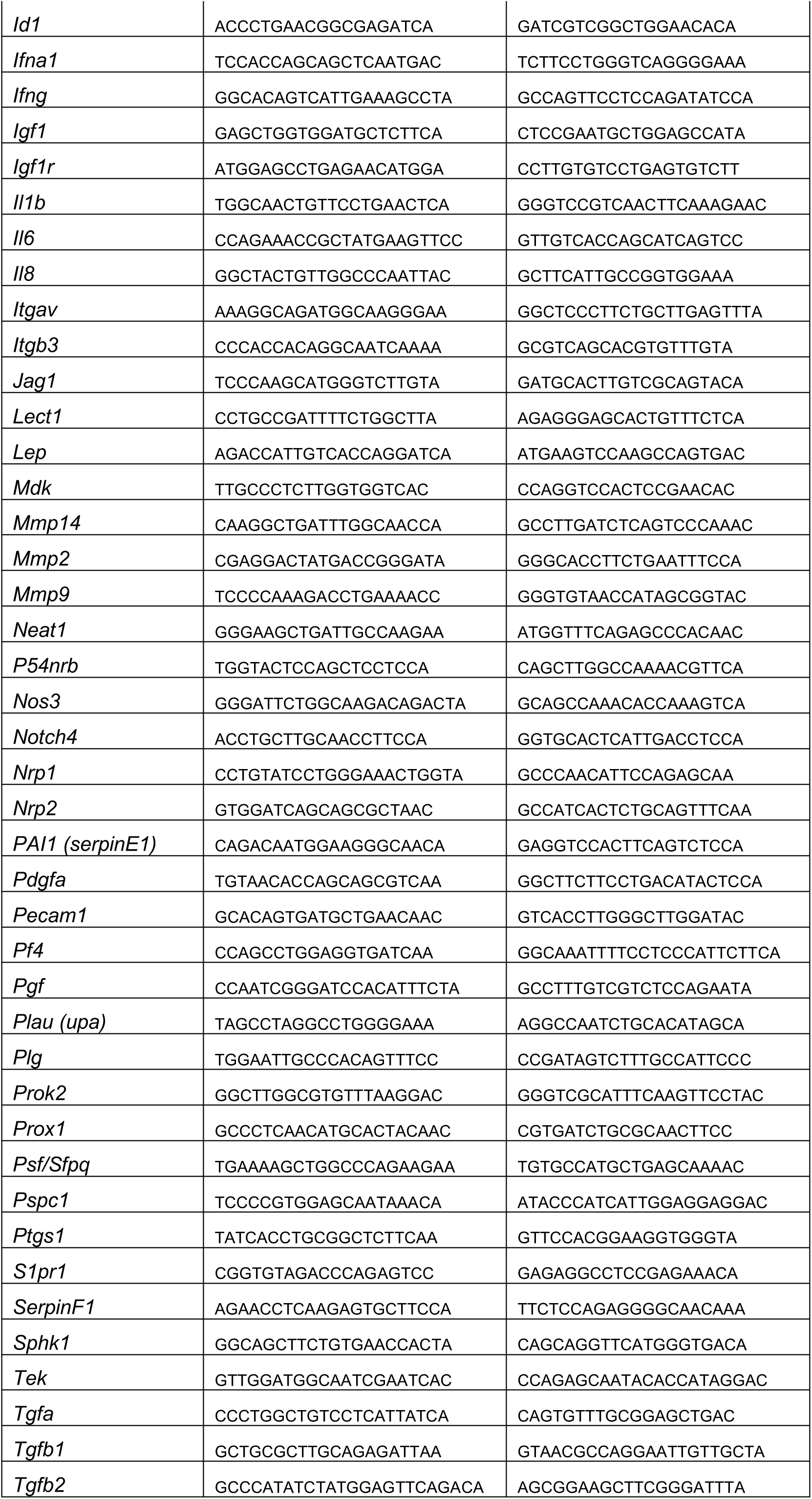
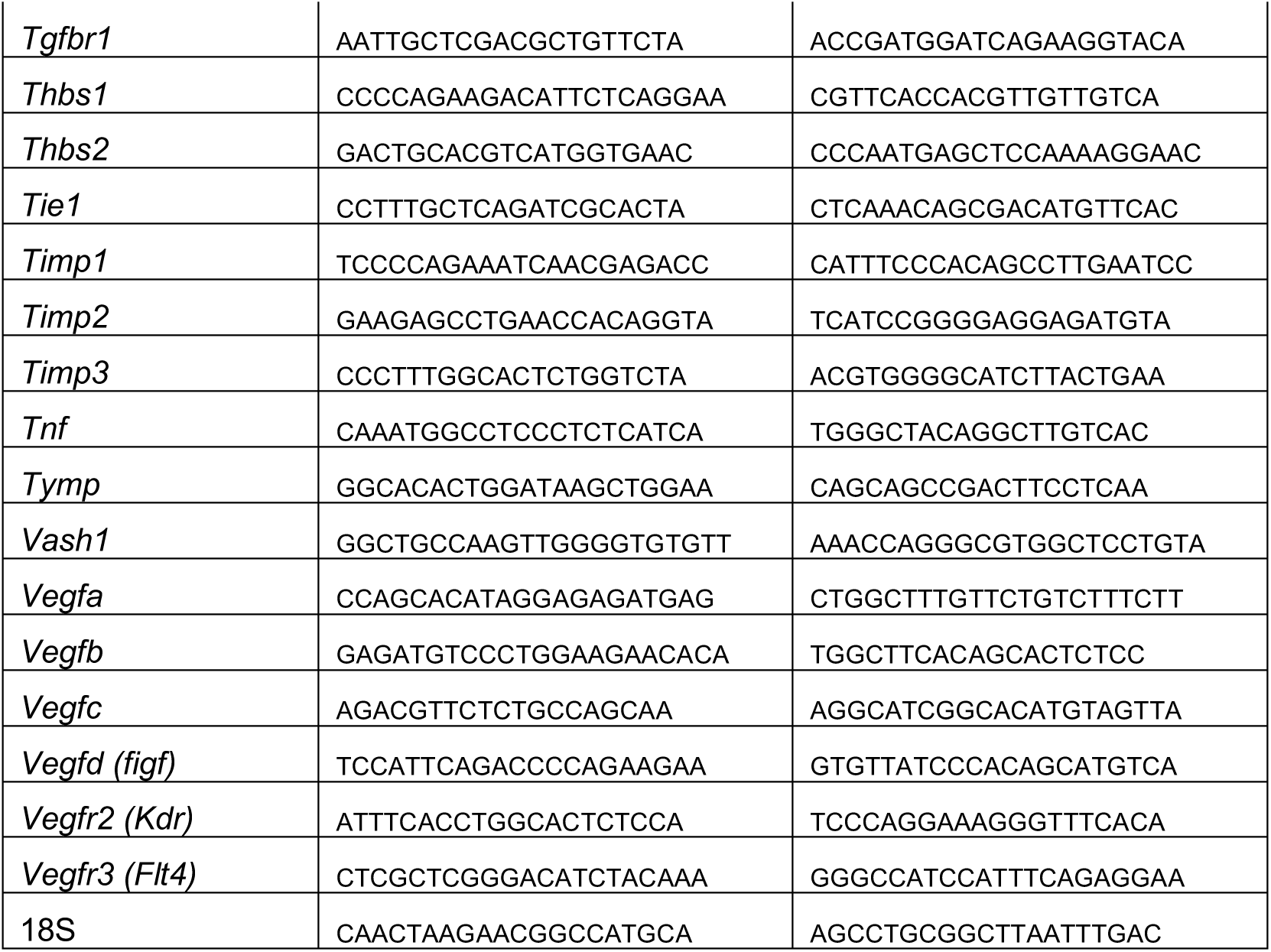
List of genes and primer couples used in the Fluidigm Deltagene PCR array.

**EV Table 7.**
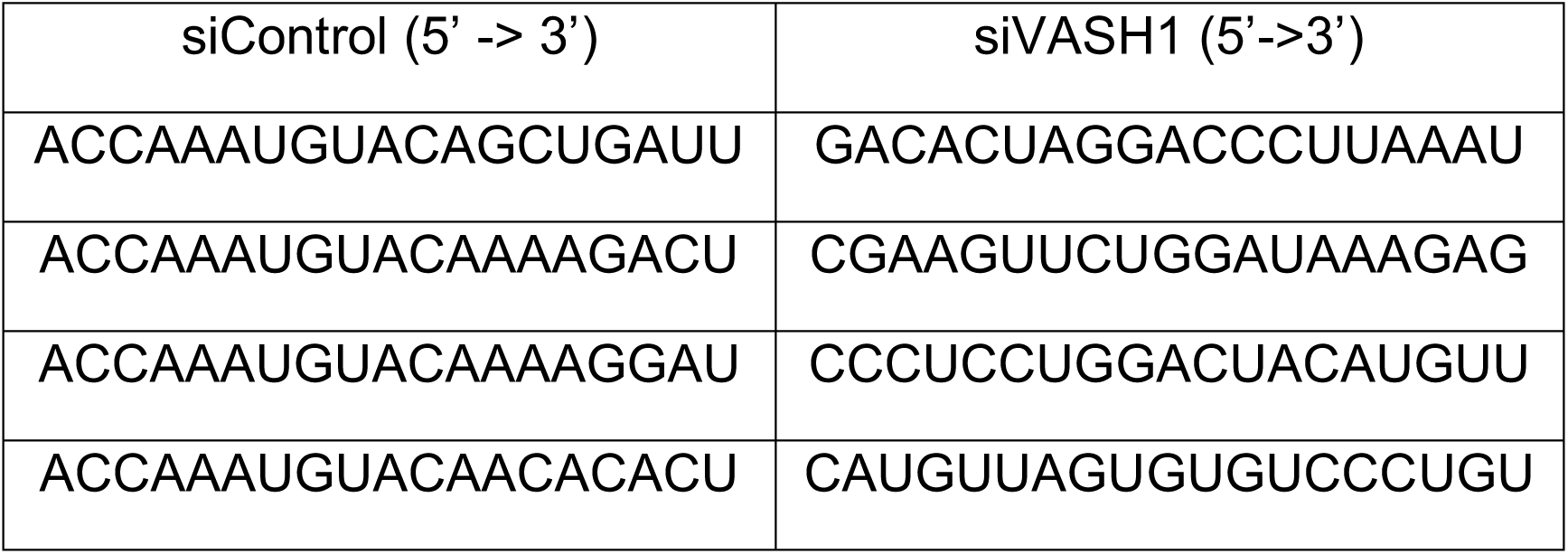
SiRNA sequences. The sequences of the four siRNAs present in the siControl and siVASH1 Smartpools are indicated.

